# Structures of tRNA-bound CRISPR-Cas13 reveal universal HEPN RNase mechanisms

**DOI:** 10.64898/2025.12.03.690598

**Authors:** Roland W. Calvert, Brooke K. Hayes, Her Xiang Chai, Yongyi Peng, Cheng Huang, Hariprasad Venugopal, Cyntia Taveneau, Jovita D’Silva, Joseph Rosenbluh, Chen Davidovich, Chris Greening, Rebecca S. Bamert, Gavin J. Knott

## Abstract

Ribonucleases (RNases) are ubiquitous drivers of RNA metabolism across life. Higher eukaryotes and prokaryotes nucleotide-binding (HEPN) domain-containing RNases form a nuclease superfamily whose mechanisms of substrate selection and catalysis remain poorly understood. Here, we report substrate preferences for diverse Cas13 HEPN RNases, revealing precise cleavage of tRNA anticodons and acceptor stems. Using cryo-EM, we present a series of activated *Leptotrichia buccalis* Cas13a (LbuCas13a) structures engaged with substrate tRNA. We show LbuCas13a captures tRNA through shape- and sequence-specific recognition, cleaving U-rich anticodons independently of tRNA modifications. Leveraging these insights, we reprogram Cas13 specificity and engineer variants with accelerated RNase activity. These findings establish the molecular basis for Cas13 tRNase activity and indicate that the HEPN RNase superfamily obeys universally conserved mechanisms of substrate recognition and catalysis.

## Main Text

Higher Eukaryotic and Prokaryotic Nucleotide-binding (HEPN) RNases are a superfamily of widespread endoribonucleases characterized by dimeric α-helical domains bearing RφX_3_H HEPN-motifs (*1, 2*). HEPN RNases are central to key eukaryotic pathways, including pre-rRNA maturation (Las1L-PNK (*3, 4*)), the unfolded protein response (Ire1 (*5, 6*)), and human innate immunity (RNase L (*7*)), and have widespread roles in microbial immunity (*8–10*). Despite the functional representation of the HEPN RNase superfamily across all domains of life, the mechanisms driving HEPN substrate selection and catalysis have remained elusive. CRISPR-Cas13, model HEPN RNases, are microbial RNA guided adaptive immune systems that restrict foreign genetic material (*11–13*). Cas13 systems are highly diverse in sequence and structural architecture with numerous subtypes and recently described ancestral systems (*14–17*). Across all subtypes, base pairing between CRISPR RNA (crRNA) and foreign activator RNA (aRNA) drives conformational changes that align and subsequently activate a pair of intramolecular HEPN domains (*12, 13, 18*). The activated HEPN endoribonuclease cleaves aRNA (*cis*-cleavage) and off-target RNA (*trans*-cleavage or collateral activity), driving host dormancy and restricting bacteriophage replication (*19, 20*). Programmable *cis*-cleavage enables CRISPR-mediated RNA knockdown (*21, 22*). In contrast, *trans*-cleavage activity has been leveraged in CRISPR-based RNA diagnostics and phage engineering (*13, 23, 24*). There is emerging evidence to suggest that the Cas13 HEPN RNase may exhibit a collateral preference for tRNA (*25*) akin to the viral dsRNA activated human HEPN RNase L (*26, 27*). However, current mechanistic models are either derived from enzymatically inactive complexes or from HEPN RNases lacking cognate substrates (*18, 28, 29*).

To shed light on the mechanisms within the HEPN RNase superfamily, we profiled substrate preferences across representative Cas13a and Cas13d enzymes to reveal site-specific cleavage of tRNA anticodons or acceptor stems. Using wild type *Leptotrichia buccalis* (Lbu) Cas13a as a model system (*30–32*), we present the structural and mechanistic basis for HEPN RNase activity using six high resolution cryo-EM structures. Our analysis of the catalytic center coordinating non-cleavable tRNA uncovers the basis of substrate recruitment, nucleotide specificity, and resolves the long-standing questions around HEPN RNase catalytic mechanisms. Finally, we reveal determinants for RNA substrate specificity and the core catalytic mechanisms driving HEPN RNase activity across biology.

### Diverse Cas13 HEPN RNases site-specifically cleave tRNA

CRISPR-Cas13 HEPN RNases are described as non-specific endoribonucleases with short mono- or di-nucleotide preferences (*24, 33*). *Leptotrichia shahii* (Lsh) Cas13a was recently reported to have anticodon tRNase (ACNase) activity, suggesting that diverse Cas13 HEPN RNases may harbor more complex substrate RNA preferences (*25*). To explore this, and avoid the confounding role of cellular RNases, we treated extracted total RNA from *E. coli* with recombinant Cas13 enzymes, including LbuCas13a, *Lachnospiraceae bacterium* (Lba) Cas13a, and *Ruminococcus flavefaciens* (Rfx) Cas13d (**Fig. S1-2**). Calculating fold-change between enzyme-treated and untreated samples revealed changes in coverage reflecting RNA fragmentation by activated Cas13 (**Fig. 1A**). Across all enzyme treatments, tRNA displayed the most statistically significant changes in comparison to rRNA and mRNA (**Fig. 1A, S3**). In addition, rRNA was well-protected from Cas13 activity within intact ribosomes (**Fig. S4**). Examining total RNA cut sites revealed consensus single (LbuCas13a: NꜜU, LbaCas13a: AꜜN) or di-nucleotide (RfxCas13d: UꜜU) cleavage signatures (**Fig. 1B, S5**). Within tRNA specifically, LbuCas13a and RfxCas13d cut sites occurred within the anticodon (**Fig. 1C-D, S6-7**). LbaCas13a displayed no ACNase activity, but highly site-specific activity within tRNA acceptor stems (AꜜCCA) (**Fig. 1C-D, S8**). To validate LbuCas13a and RfxCas13d site-specific ACNase activity, we investigated activity on a synthetic tRNA^Lys(UUU)^ substrate. When incubated with activated LbuCas13a, we observed tRNA^Lys(UUU)^ cleavage into long-lived tRNA halves (**Fig. S9**). Moreover, long-lived tRNA halves were generated from the anticodon stem-loop (ASL^Lys(UUU)^) alone (**Fig. S10A, S11-12**), suggesting unmodified ASL are sufficient for precise anticodon cleavage. In further agreement with RNA-seq, we observed no tRNase activity on ASL^Phe(GAA)^, whereas ASL^Asp(GUC)^, ASL^Lys(UUU)^, and ASL^His(GUG)^ were cleaved within the anticodon (**Fig. S10A and S10B**). Using deoxy-blocked nucleotides, we precisely mapped preferred ASL^Lys(UUU)^ cut sites to U34ꜜU35 and U35ꜜU36, consistent with RNA-seq (**Fig. S10C**). Similarly, RfxCas13d cleaved ASL^Lys(UUU)^ within the anticodon at positions U34ꜜU35 and U35ꜜU36 (**Fig. S13**).

**Figure 1.**
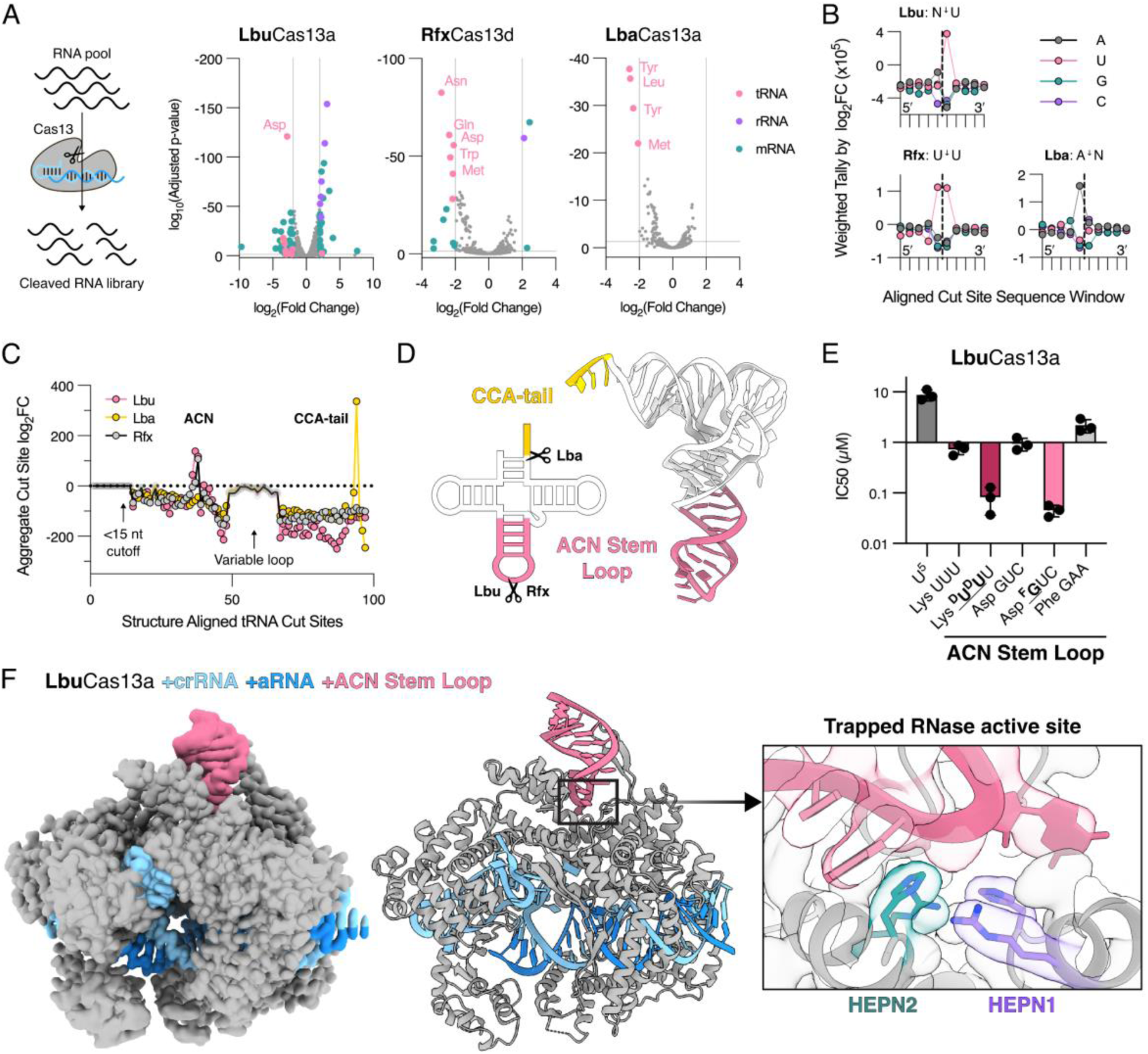
Cas13 preferentially cleaves tRNA. (**A**) RNA sequencing of total *E. coli* RNA with active LbuCas13a, LbaCas13a, and RfxCas13d. Volcano plots show differential transcriptome coverage by Cas13 activity at T=30 min (n=2: adj. P=0.05, |log_2_(FC)|=2). (**B**) FC-weighted nucleotide tallies at T=30 min of cut-aligned sequence windows show Cas13-specific nucleotide signatures. (**C**) Aggregate FC of aligned tRNA cut sites by structure from differential 5′ end analysis (5′ ends= 1-14 are undetectable): LbuCas13a (pink), LbaCas13a (yellow) and RfxCas13d (grey). (**D**) Cut sites (scissors) mapped onto a 2D (left) and 3D (right) tRNA (6UGG (*61*)): ACN stem loop in pink, CCA-tail yellow. (**E**) IC50 values for U^5^ and tRNA ACN stem loop (wildtype or non-cleavable) substrates determined by competition assays with LbuCas13a. (**F**) Structure of LbuCas13a:crRNA:aRNA (grey, light blue and dark blue respectively) bound to tRNA^Asp^ ACN stem loop (pink). Left) EMReady post-processed cryoEM map of the structure, Middle) Flat cartoon of structure, Right) Zoom of HEPN active site showing LbuCas13a and tRNA^Asp^ ACN (pink) interactions. LbuCas13a is shown with the cryoEM map superimposed. Crucial HEPN1 (R472, H477, green) and HEPN2 (R1048, H1053, purple) residues shown as sticks and colored by heteroelement.

The discrete ACNase activity suggested a Cas13a and Cas13d HEPN RNase preference for structured U-rich ASLs. To explore this, we indirectly measured substrate binding to the activated LbuCas13a HEPN using RNase competition assays (**Fig. 1E, S14-20**). Consistent with a preference for structured U-rich ASLs, both ASL^Asp(GUC)^ and ASL^Lys(UUU)^ outcompeted labelled U-rich homoribopolymers (**Fig. 1E**). Incorporation of non-cleavable nucleotides (2ʹ-fluoro modified ASL^Asp(fGUC)^ or 2ʹ-deoxyuridine modified ASL^Lys(dUdUU)^) further improved the IC_50_, suggesting a mode of HEPN RNase competitive inhibition (**Fig. 1E**). In contrast, and consistent with a mechanism of both structure and sequence-based recognition, the non-cleavable ASL^Phe(GAA)^ lacking NꜜU within the anticodon was less effective (**Fig. 1E**). Given that synthetic ASL substrates recapitulated LbuCas13a cleavage of modified full-length tRNA, we solved a series of cryo-EM structures at every stage of wildtype LbuCas13a activity to comprehensively understand the mechanism of HEPN RNase substrate capture and cleavage (**Table S1, Fig. S21-25**). We reconstituted Lbu-crRNA (2.3Å resolution), activated Lbu-crRNA-aRNA poised for substrate recognition (2.6Å resolution), inactive Lbu-crRNA engaged with ASL^Lys(dUdUU)^ (2.5Å resolution), activated Lbu-crRNA-aRNA engaged with ASL^Lys(dUdUU)^ (3.1Å resolution), and activated Lbu-crRNA-aRNA engaged with ASL^Asp(fGUC)^ (**Fig. 1F**, 2.6Å resolution). In all states, we clearly resolved the LbuCas13a protein, crRNA, aRNA, as well as the conserved active site HEPN1 (H472, R477) and HEPN2 (R1048, H1053) residues in the apo-state or engaged with substrate ASL^Lys(dUdUU)^ or ASL^Asp(fGUC)^ **(Fig. S25**).

### Cas13 HEPN RNase accessory elements facilitate substrate RNA capture

The cleavage profiles for LbuCas13a, RfxCas13d, and LbaCas13a demonstrated a diversity of substrate preferences despite having the same conserved active site. We hypothesized that this bias is imposed by structural elaboration of the HEPN fold with bespoke “accessory” elements surrounding the active site (*2*). Our structural data shows both ASL^Lys(dUdUU)^ and ASL^Asp(fGUC)^ engage deep within the active site groove formed between the juxtaposed LbuCas13a HEPN1 and HEPN2 domains bearing accessory β-sheets and α-helices, respectively (**Fig. 1F, 2A**). Comparing structures of Lbu-crRNA bound to ASL^Lys(dUdUU)^ with activated Lbu-gRNA-aRNA bound to ASL^Lys(dUdUU)^ revealed an ordered transition of the substrate towards the HEPN active site (**Fig. 2A**). Strikingly, the asymmetry of the HEPN accessory elements imposed a preferred ASL orientation with the substrate engaged along the minor groove contacted by the HEPN1 β-stranded accessory element we call the “finger” (**Fig. 2A**) (**Movie S1-2**). Across all states, the bound ASL makes several sequence independent contacts with HEPN accessory elements through charged contacts along the phosphodiester backbone of the ASL (**Fig. 2B**). Consistent with these structural observations, changing the sequence without altering the ASL secondary structure had no impact on activity (**Fig. S11D**). Furthermore, our RNA-seq analysis revealed multiple tRNAs cleaved by the activated LbuCas13a HEPN RNase, consistent with shape recognition of the stem region (**Fig. S6-8**). To confirm the importance of the HEPN accessory elements in guiding substrate engagement, we purified a panel of LbuCas13a constructs with charge-neutralized accessory elements on HEPN2 (Helix A: K1004A, K1011A, K1014A and K1017A; Helix B: K1036A, K1039A, Q1040A and K1042A), HEPN1 (Finger C: K409A, N414A, K415A and K419A), or with deleted Finger (Δ409-421) (**Fig. 2C, S26**). Across a panel of RNA substrates, we observed that any alteration to the HEPN accessory elements compromised RNase activity (**Fig 2D, S26-27**), suggesting that the ASL interactions represent a general mechanism of substrate tRNA or ssRNA capture and engagement.

**Figure 2.**
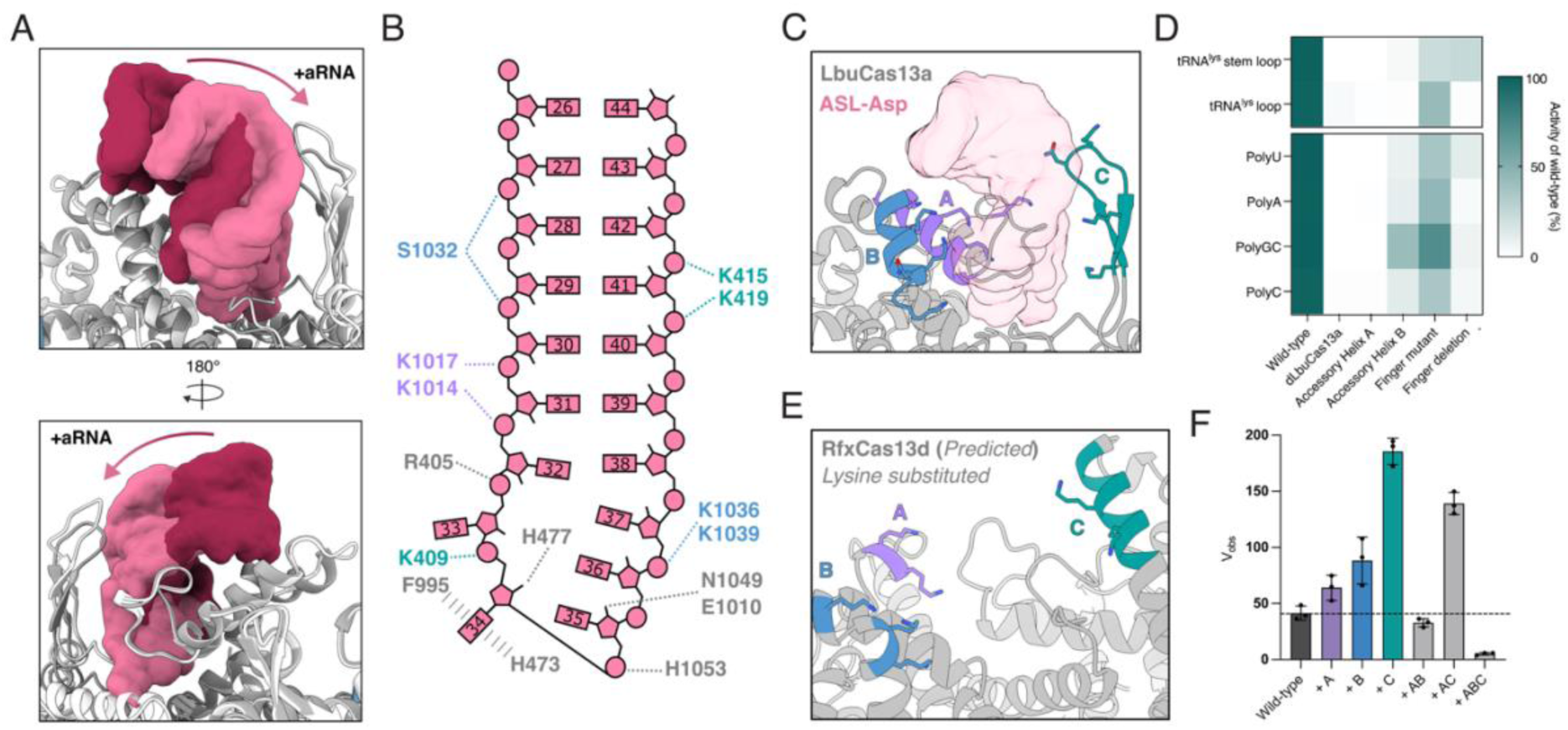
tRNA substrate capture by Cas13 is mediated by accessory elements. (**A**) Movement of ASL (surfaces) in the active site of LbuCas13a (cartoon). Binary LbuCas13a (LbuCas13a:crRNA, light grey) with ASL^Lys(dUdUU)^ map (red). Ternary LbuCas13a (LbuCas13a:crRNA:aRNA, dark grey) with ASL^Asp(fGUC)^ map (pink). (**B**) Contact map illustrating LbuCas13a sidechain residues near ASL^Lys(dUdUU)^ and ASL^Asp(fGUC)^ phosphodiester (circles), ribose (pentagons), or bases (rectangles). (**C**) Ternary LbuCas13a (flat cartoon, grey) bound to tRNA^Asp(fGUC)^ (transparent pink). Accessory helix A (K1004, K1011, K1014, K1017) purple, accessory helix B (K1036, K1039, Q1040, K1042) blue, and accessory finger green (K409, N414, K415, K419). (**D**) LbuCas13a accessory element mutants. Alanine mutations as listed in (B). Finger deletion (Δ409-421) colored green in (B); dLbuCas13a (H477N, H1053N). Activity (n=3) shown as percentage of wildtype for tRNA ASL^Lys(UUU)^ and disordered tRNA^Lys^ loop (top panel), and ribopolymer reporters (bottom panel: U^5^, A^5^, CGCGC and C^5^). (**E**) Ternary RfxCas13d AlphaFold3 model (RfxCas13d:crRNA:aRNA) shown as grey flat cartoon. Equivalent accessory elements in purple, blue and green. (**F**) Observed velocity (n=3, error bars SD) for RfxCas13d mutations with U^5^ reporter. Wildtype in dark grey. Mutations for A: A188K, A191K (purple), B: E753K, E754K (blue) and C: N897K, E898K, E901K (green). Grey bars stacked mutations of previous.

To explore the broader role of accessory elements in Cas13 ACNase activity, we generated a high-confidence prediction of the active RfxCas13d-crRNA-aRNA complex (**Fig. S28**). Examining the HEPN RNase groove revealed an arrangement of structurally analogous HEPN1 and HEPN2 accessory elements (**Fig 2E**). Comparing the model RfxCas13d HEPN to activated LbuCas13a revealed a similar topological, but less charged landscape of HEPN1 and HEPN2 accessory elements (**Fig. S29**). Supercharging either Helix A or Helix B (but not both) with positively charged amino acid substitutions increased activity against a suite of substrates (**Fig. 2F, S30-33**). Consistent with the role of accessory elements in governing global Cas13 HEPN RNase substrate specificity, the tRNA acceptor stem cleaving LbaCas13a possesses a divergent set of accessory elements around the composite HEPN RNase active center (**Fig. S34**). Collectively, these structural and functional data support a model where Cas13 HEPN1 and HEPN2 accessory elements work cooperatively to recruit and guide structured tRNA and unstructured RNA substrates towards the HEPN active site.

### Substrate nucleotide recognition is divergent across Cas13 HEPN RNases

Various HEPN RNases are reported to have nucleotide sequence preferences (*29, 33, 34*). Cas13 systems are allosterically activated by crRNA-aRNA hybridization, with previous studies reporting the crRNA and HEPN domain re-arrangements that juxtapose the HEPN motifs for catalysis (*18, 28, 35*). Our structure of ASL^Lys(dUdUU)^ loaded but inactive LbuCas13-crRNA revealed an incomplete engagement with the HEPN where access to the nuclease is occluded by a HEPN1 substrate gating loop (aa396-406) (**Fig. 3A**). Re-arrangement of the substrate gating loop upon aRNA binding facilitates ASL access into the HEPN active site via induced fit whereby the anticodon triplet is unstacked, positioning the scissile phosphate above the juxtaposed HEPN motifs (**Fig. 3B**). The structure of LbuCas13a engaged with ASL^Asp(fGUC)^ revealed the unstacked anticodon is exclusively read out by HEPN2 N1055 with base specific contacts to the Watson-Crick face of U35 (**Fig. 3A-B**). Consistent with base specific uridine read out, pseudouridine modification or changes to the surrounding loop sequence through base substitutions or inversions had no effect on HEPN RNase activity (**Fig. S10D, S35**). Mutagenesis to proximal residues in the substrate gating loop (N400A, N400G, R405A) or HEPN2 (Q1007A, N1049A) did not alter the substrate preference profile (**Fig. 3C**). In contrast, and consistent with N1055 being the sole determinant of the NꜜU preference, N1055A, N1055G, or N1055D ablated LbuCas13a HEPN RNase activity against U^5^ homoribopolymers (**Fig. 3C**). By expanding the HEPN2 specificity pocket through N1055A or N1055G substitutions, we effectively relaxed the HEPN RNase specificity to now accommodate diverse short non-U containing RNA substrates, albeit at a cost to overall catalytic efficiency (**Fig. 3C, S36-37**). Consistent with the programmability of the HEPN2 mediated substrate preference, N1055D substitution retained the pocket volume but altered the hydrogen bonding network to select for C^5^ homoribopolymers (**Fig. 3C**).

**Figure 3.**
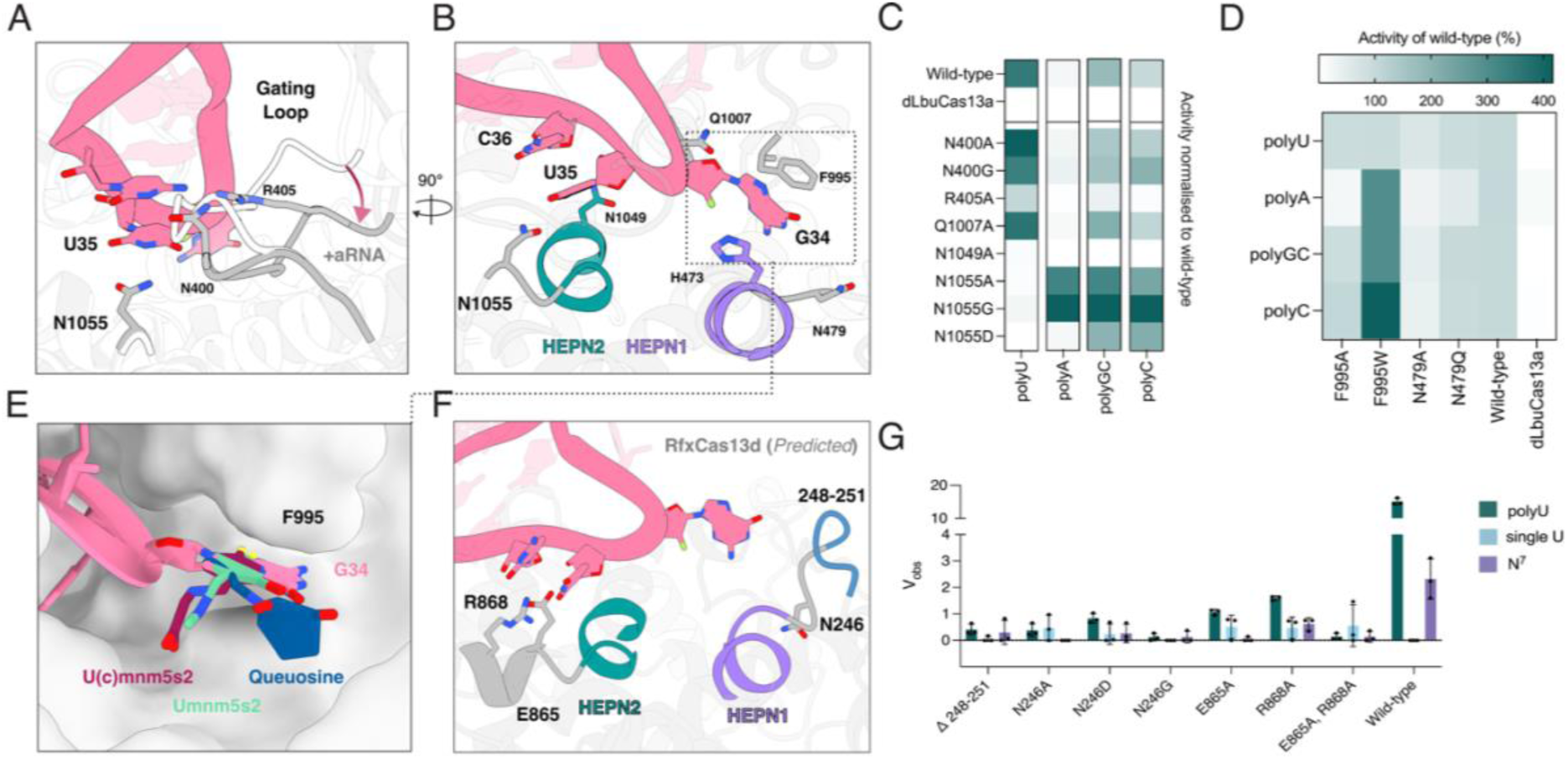
LbuCas13a substrate gating and LbuCas13a/RfxCas13d nucleotide specificity. Proteins shown in grey, tRNA pink. Key residues and ACN nucleotides shown as sticks, colored by heteroelement; dLbuCas13a (H477N, H1053N). (**A**) and (**B**) Interactions between LbuCas13a and ASL^Asp(fGUC)^ (**A)** movement of gating loop between binary (light grey) and ternary (dark grey) states. (**B)** 90° rotation of (A), HEPN1 purple, HEPN2 green. (**C)** Activity (n=3, normalized to wildtype) of LbuCas13a substrate recognition mutants measured by ribopolymer reporters (U^5^, A^5^, CGCGC and C^5^). (**D**) Activity (n=3) of LbuCas13a G-flip base stacking mutants, against substrates as in (C). **(E**) G-flip binding pocket for LbuCas13a (surface representation) with ASL^Asp(fGUC)^ (cartoon, G34 stick). Wobble base (position 34) modifications overlaid: U(c)mnm^5^s^2^ (purple), Umnm^5^s^2^ (green) and Queuosine (blue). (**F**) AlphaFold3 ternary RfxCas13d model (grey cartoon): HEPN1 purple, HEPN2 green. Substrate ASL^Asp(fGUC)^ overlaid in pink, the blue loop (248-251) was deleted. (**G**) Activity (n=3, error bars SD) of RfxCas13d substrate recognition mutants. Reporters as listed (U^5^, single U (AAUAA) and N^7^).

HEPNs are metal ion-independent nucleases, suggesting that they distort the substrate RNA to mediate cleavage (*36*). Substrate ASL engagement within the LbuCas13a HEPN RNase results in an induced fit rearrangement of the anticodon triplet (**Fig. 3B**). Within the LbuCas13a-ASL^Asp(fGUC)^ bound structure, we observed a flexing of the tRNA substrate within the pocket correlating to the appearance and disappearance of discrete density describing the wobble base G34 (**Fig. S38, Movie S1**). Modelling these two states revealed a clear step of base-specific read out that precedes the wobble position G34 base flipping towards HEPN1 and positioning of the scissile phosphate above the HEPN motif active site (**Fig. S39**). Once flipped, the HEPN1 pocket accommodates G34 in a wide non-specific pocket where the purine stacks between F995 and H473 (**Fig. 3B**). While both H473N and F995A substitution are tolerated, the activity profile of the HEPN RNase could be improved through F995W substitution (**Fig. 3D, S40**), consistent with the generic role of this pocket in accommodating bases of any type. Furthermore, aligning chemical structures of tRNA wobble bases (position 34) bearing modifications revealed that all could be accommodated with no re-arrangements (**Fig. 3E**), consistent with the modification agnostic activity of the LbuCas13a HEPN RNase.

Our RNA-seq analysis revealed a di-nucleotide UꜜU preference for RfxCas13d that drives site specific cleavage of a restricted pool of *E. coli* tRNA anticodons (**Fig. 1B-D**). Using our RfxCas13d AlphaFold3 prediction (**Fig. S28**) and superimposing the unstacked ASL^Asp(fGUC)^ suggested discrete HEPN1 and HEPN2 nucleotide binding pockets either side of the catalytic center (**Fig. 3F**). Mutations to either of these features abolished HEPN RNase activity (**Fig. 3G, S30, S41**), consistent with a strict requirement for di-nucleotide read out prior to catalysis. Taken together, our data demonstrate how Cas13 HEPN RNases mediate base specific contacts within the anticodon triplet to engage the substrate RNA above the HEPN RNase active site for cleavage.

### Catalytic mechanism of the Cas13 HEPN RNase

HEPN RNases function via a dimeric juxtaposed arrangement of RφX_3_H HEPN-motifs. While the HEPN residues involved are well defined, the role of each residue in catalysis remains elusive (*2*). To understand how catalysis proceeds we investigated our high-resolution structure of activated LbuCas13a binding ASL^Asp(fGUC)^ where the scissile phosphate is engaged with the catalytic tetrad (**Fig. 4A**). In this configuration, the conserved HEPN1 H477 contacts 2′-F of G34, HEPN2 H1053 contacts the bridging 5′-O of U35, and the conserved Arg residues (R472 and R1048) coordinates the scissile phosphate (**Fig. 4A**). Intriguingly, we observed H473 in the apo-HEPN state had dual-occupancy comparable to Las1 (*3*) (**Fig. S42**) and substrate binding biases this equilibrium towards an active state (**Fig. 4B**). Given the strict preferred orientation of ASL engagement mediated by HEPN accessory elements, we reasoned that HEPN1 H477 likely mediates attack on the 2′-OH to initiate the reaction and generate a 2′3′ cyclic phosphate (cP) product. The positioning and strict conservation of HEPN2 H1053 suggested it may participate by reducing the cP to 3′-P via hydrolysis — a two-step mechanism analogous to the highly catalytically efficient RNase A or RNase T1 (*37, 38*). To explore this, we validated the end chemistry of the products generated by LbuCas13a HEPN RNase activity using HPLC-MS, which revealed almost exclusively cP products (**Fig. 4C, S43-44**). In contrast, substrate cleavage by RNase T1 produced detectable 3′-P alongside the cP final product **(Fig. 4C, S45)**. Collectively, our structural and biochemical data support a coherent catalytic model for the LbuCas13a HEPN RNase (**Fig. 4D**). Activated LbuCas13a exhibits an aligned composite HEPN active site with all RφX_3_H residues in proximity. Substrate RNA is read out by HEPN2 N1055, which specifically binds U along the Watson-Crick face. Following readout, the preceding ribose is locked into a suitable pre-catalytic state through base flipping, stabilized within a sequence-nonspecific HEPN1 pocket. HEPN1 H477 activates the 2′-OH to initiate an in-line nucleophilic attack on the scissile phosphate to generate a pentavalent reaction intermediate stabilized by the remaining conserved HEPN His/Arg residues (R472, R1048, and H1053). Once the transition state is resolved and the phosphodiester is broken, the fragment products with 2′3′-cP and 5′-OH ends are released, and the active site residues are regenerated by the solvent to the ground state for another catalytic cycle (**Fig. 4D**).

**Figure 4.**
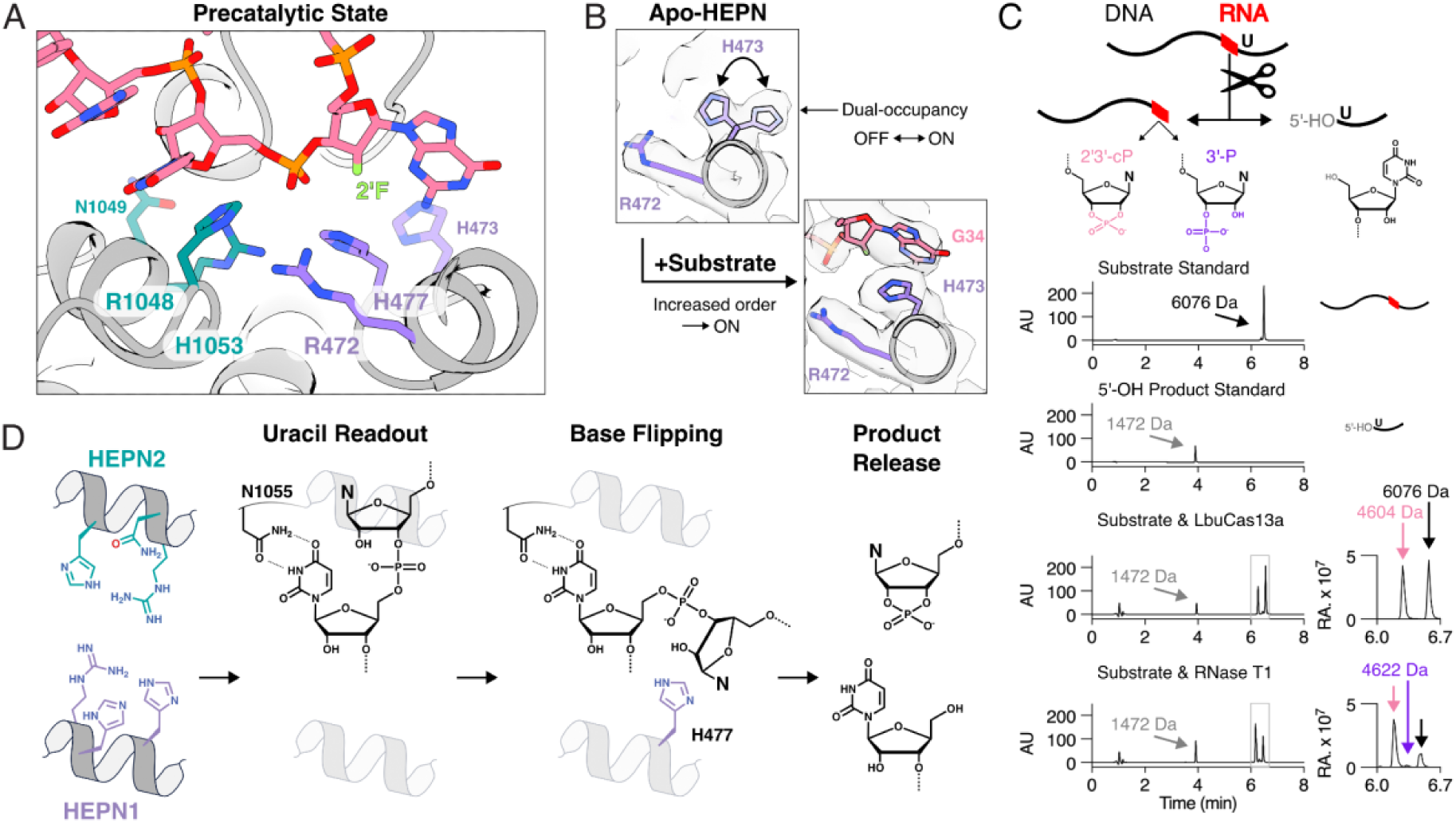
Catalytic mechanism of RNA cleavage by HEPN active site. LbuCas13a (grey, cartoon) with HEPN1 purple and HEPN2 green; ASL^Asp(fGUC)^ (pink). Residues involved in catalysis and ACN residues (G34, U35, C36) shown as sticks, colored by heteroelement. (**A**) Precatalytic state of LbuCas13a with U35 phosphate of tRNA^Asp^ in proximity to the HEPN active site. (**B**) H473 exhibits dual occupancy in apo-state (top panel), but not in the substrate-bound state (lower panel) (**C**) LbuCas13a cleavage of a single RNA base (red) within a DNA polymer (black) to determine the 3′ product. (Top) Cleavage to generate a 5′-OH product (1472 Da), and either a 3′-P (4622 Da) or 2′3′-cP (4604 Da) product. (Bottom left) HPLC chromatograms (absorbance at 280nm; AU), in order: substrate (6076 Da); 5′-OH product (1472 Da); substrate +LbuCas13a (showing substrate, and product masses); substrate +RNase T1 (produces 3′-P). (Bottom right) relative abundance based on LC-MS. (**D**) Catalytic mechanism of LbuCas13a. Substrate U is recognized by N1055 then the adjacent 5′ nucleotide is flipped for HEPN catalysis, initiated by H477 acting as the general base with R472, R1048 and H1053 stabilizing the reaction coordinate. The resulting fragments have 2′3′-cP and 5′-OH ends.

## Discussion

In this study, we uncovered site specific tRNase activity across diverse CRISPR-Cas13 systems. We determined six LbuCas13a cryo-EM structures spanning the multistate process of HEPN RNase activation, substrate tRNA recruitment, and cleavage. We proposed a complete model for LbuCas13a HEPN RNase activity that includes the mechanisms of tRNA shape recognition, anticodon readout, substrate remodeling, and the chemistry of catalysis (**Fig. 4C**, **Fig. 5A**). Our findings are applicable to the broader HEPN RNase superfamily and provide key mechanistic insights into the function of tRNA cleaving RfxCas13d (**Fig. 5B**) and eukaryotic tRNA cleaving RNase L and Ire1 (**Fig. 5C-D**).

**Figure 5.**
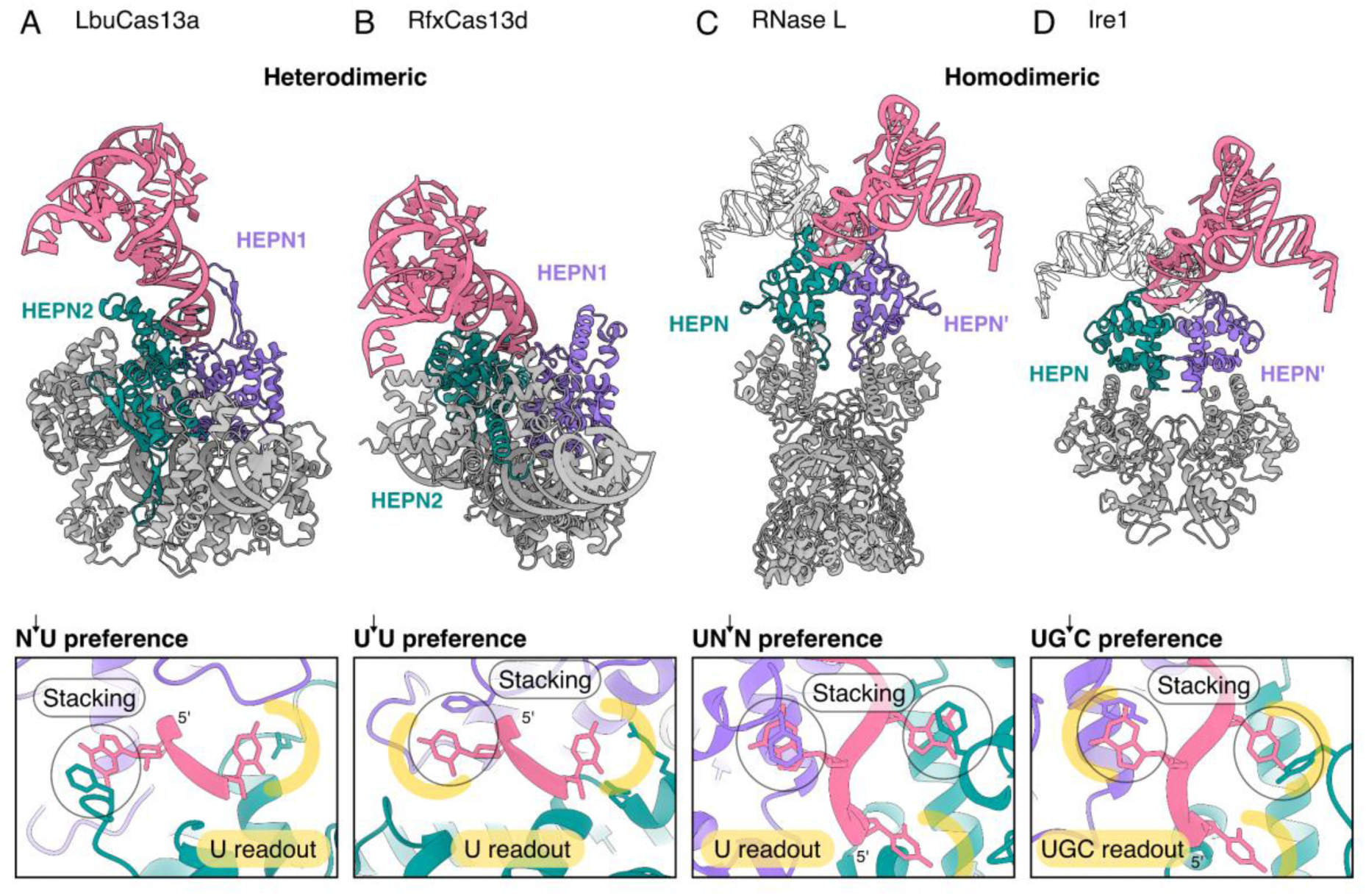
Models of tRNA anticodon capture and cleavage by diverse HEPN RNases. Analysis of HEPN domains in (**A**) LbuCas13a, (**B**) RfxCas13d, (**C**) human RNase L, and (**D**) *S. cerevisiae* Ire1 (3FBV (*62*)). (Top) Structures superimposed with tRNA Asp (6UGG (*61*)) best fitted to HEPN domains colored in purple and green. Heterodimeric HEPN RNases bind substrate in only one orientation (pink) while homodimeric ones may accommodate rotationally symmetric orientations (pink and white). (Bottom) Substrates are conformationally induced to splay bases in HEPN RNases via non-specific stacking interactions (black circle) and specific nucleotide recognition (yellow lines). Experimentally determined nucleotide preference for each enzyme listed above each HEPN zoom. Predicted substrate-bound state for RNase L produced from AlphaFold3 is also grafted on Ire1 with a single G→C mutation.

Our findings describe how HEPN domain asymmetry, evolutionary decoupling, and accessory element elaboration (*16, 17*) contribute to the diversification of Cas13 enzyme specificity and efficiency. LbuCas13a HEPN accessory elements are highly positively charged to drive substrate recruitment and rapid turnover (*33*). Consistent with this, supercharging RfxCas13d (**Fig. 2**) or inserting RNA-binding domains into *Leptotrichia wadei* (Lwa) Cas13a enhances catalysis (*39*). The asymmetry of the HEPN enforces a preferred substrate tRNA orientation to deliver the anticodon for base specific readout and anticodon unstacking, not unlike tRNA modification enzymes (*40*). In contrast to LbuCas13a (NꜜU), the distinct preferences of LbaCas13a (AꜜN) and RfxCas13d (UꜜU) suggest alternative modes of substrate engagement with readout occurring on the opposite or both sides of the HEPN RNase. While most RNases specifically engage nucleotides to distort the backbone for in-line attack (*36*), stricter specificity increases the threshold for substrate engagement, an efficiency-specificity trade off that could explain RfxCas13d catalytic inefficiency and resistance to simple HEPN reprogramming. Our data show that Cas13 HEPN RNases are under strong diversifying selection with variable nucleotide preferences, mechanisms of substrate engagement, and varied site-specific cleavage of discrete tRNA.

While our study focused on asymmetric Cas13 HEPN RNases, the majority of characterized HEPN RNases are symmetric intermolecular homodimers like RNase L and Ire1 (**Fig 5C-D**). RNase L is both a general RNase and a site-specific human tRNA^Pro(UGG)^ ACNase (*27*) that has a known UNꜜN specificity. Given RNase L showcases a binding mode that does not involve either nucleotide adjacent to the scissile phosphate (*29*), these non-specific interactions appear to be mediated by base stacking, while a specific U readout must occur downstream (**Fig. 5C**). Ire1 homologs are also reported to have ACNase activity for tRNA^Gly(GCC)^ where they stringently read UGꜜC (*34, 41*). Alignment of this with our findings suggest that a specific U readout occurs while fully engaged in the active site, but non-specific base stacking occurs with the readout G and C nucleotides. This demonstrates the variety of nucleotide readouts and the highly plastic nature of substrate engagement but also suggests that base readout may occur in multiple steps for HEPN RNases that recognize multiple bases (**Fig. 5D**). This aligns with the RfxCas13d model (**Fig. 5B)** that also shows a single U readout in the active site, but a secondary recognized base that is involved in non-specific base stacking when engaged in catalysis. Further, Csm6, another symmetrical HEPN RNase, has been resolved binding an RNA that relaxes to a conformation with flipped bases (*42, 43*). While highly plastic, HEPN RNase nucleotide readout is a universal feature that functions not just for substrate selection, but also to achieve an appropriately ordered pre-catalytic conformation through base slipping - a conserved mechanism among all metal-independent RNases including RNase A and T1 (*44–46*).

In consideration of the biological implications of our findings, targeted cleavage of tRNA would reduce the pool of translation capable tRNA leading to ribosome stalling (*25*). Our finding that diverse Cas13 enzymes specifically cleave tRNA is consistent with previous findings for LshCas13a (*25*) and underscores a general mode of tRNA targeting across Type VI CRISPR-Cas13. Cas13-mediated ACNase activity aligns well with HEPN RNases such as HEPN-MNT toxin-antitoxin system, PrrC, and RloC, all of which cleave tRNA (*2, 8, 47, 48*). Similarly, RNase L and Ire1 generate tRNA fragments (halves) that serve as secondary signaling molecules (*27, 41*). Given their role in microbial adaptive immunity and toxin-antitoxin ancestry (*17*), it seems likely that diverse Cas13 enzymes cooperate with accessory proteins to trigger innate immune responses, either through tRNA fragments or indirectly through perturbations to translation (*49*). Cleavage of tRNA could act as a thresholder response (*50*) and presents a clear pathway to recovery from infection or phage escape, for example, microbes encode diverse RNA repair machinery, including anticodon tRNA ligases and CCA tailing enzymes (*51*). In response, phages are known to encode diverse tRNA, RNA repair machinery, or complex tRNA modifications that could counter Cas13 (*52–54*). Alternatively, some tRNA modifications can promote cleavage by anticodon nucleases (*55*), exposing another layer of unexplored regulation. These observations point towards a breadth of diverged HEPN RNase binding and cleavage mechanisms with tRNA at the nexus (*56*).

HEPN-mediated cleavage of tRNA has broad implications for the application of CRISPR-Cas13 in biotechnology. In molecular diagnostics (*23, 24*), understanding HEPN RNase substrate preference provides avenues for engineering superior reporter substrates for detection (*57*). In phage engineering, many bacteriophages may prove to be resistant to Cas13 activity due to phage-encoded tRNA, or modification and repair machinery (*52, 54*). In mammalian cell RNA knockdown applications (*22*), our understanding of the Cas13 collateral effect may need to be carefully reassessed, given that tRNA damage may be the indirect driver of collateral activity through mammalian responses to perturbed translation (*58–60*). Taken together, our data demonstrate that while the core HEPN active site may lack specificity, its structural environment can define strict substrate preferences and underscores a critical need to consider HEPN RNase applications in the context of HEPN diversity.

## Acknowledgements

We thank the Monash Molecular Crystallisation Platform for performing the nanoDSF experiments, and Ashleigh Kropp for preparation and analysis of mass spectrometry peptide fingerprinting. We acknowledge the use of instruments and assistance at the Monash Ramaciotti Centre for Cryo-Electron Microscopy, a Node of Microscopy Australia. ARC LIEF grants (LE200100045, LE120100090) for the Titan Krios Gatan K3 Camera and for the Titan Krios. We thank Azenta Life Sciences for carrying out the library preparation and data collection of RNA sequencing. We thank the MonARCH HPC Cluster and the MASSIVE HPC facility for providing computational resources and platforms used for the RNA sequencing data analysis. We acknowledge the Monash RNA Mass Spectrometry Platform for experimentation. Molecular graphics and analyses performed with UCSF ChimeraX (*63*). This study was funded by the following sources: National Health and Medical Research Council Investigator Grant APP1175568 (GJK), Australian Government Research and Training Programme (RWC) and Snow Medical Research Foundation SMRF2021-276 (GJK). Snow Medical were not involved in the design of the study, data collection, analysis, interpretation of the data, the writing, or the decision to submit the article for publication.

## Author contributions

GJK conceived the project. RWC and BKH prepared RNA-sequencing samples with analysis carried out by YP, RWC, JR, and CG. BKH, HXC, RWC, and JDS expressed and purified proteins. RWC and BKH carried out RNA gels. RWC, CT, and HV prepared EM grids, screened, and collected data. RWC and CT processed EM data. RWC and BKH built and refined models. BKH and HXC cloned, expressed, purified, and assayed protein mutants. CH, RWC, and CD carried out mass spectrometry. RWC, BKH, RSB and GJK analyzed all data and wrote the manuscript with input from all the authors.

## Data and materials availability

Sequencing data were deposited at the NCBI Sequence Read Archive (BioProject ID PRJNA1282368). The custom scripts can also be found at GitHub (https://github.com/YongyiPeng/Cas13_RNA_cleavage). The mass spectrometry proteomics data have been deposited to the ProteomeXchange Consortium via the PRIDE (*64*) partner repository with the dataset identifier PXD065184 under Username: and Password: Q0pXr4npHd6V. Atomic structures and cryo-EM maps are available on the Protein Data Bank and Electron Microscopy Data Bank: Apo-HEPN LbuCas13a:crRNA (PDB-9OX2, EMD-70952), Apo-HEPN LbuCas13a:crRNA:aRNA (PDB-9OX0, EMD-70950), tRNA^Lys(dUdUU)^ LbuCas13a:crRNA (PDB-9OX3, EMD-70953), tRNA^Lys(dUdUU)^ LbuCas13a:crRNA:aRNA (PDB-9OX4, EMD-70954), tRNA^Asp(fGUC)^ LbuCas13a:crRNA:aRNA (State 1 PDB-9OX1, State 2 PDB-9OX5, EMD-70951). Structures not from this study include tRNA^Asp^ (**Fig. 1D, 5A-D**) (PDB-6UGG (*61*)) and Ire1 (**Fig. 5D**) (PDB-3FBV (*62*)).

## List of Supplementary Materials

Supplementary Materials and Methods

Figs. S1 to S45

Tables S1 to S4

References (65-78)

Movie S1 and S2

