## Supplementary file for "Structures of tRNA-bound CRISPR-Cas13 reveal universal HEPN RNase mechanisms"

### Materials and Methods

**Molecular biology.** All strains and plasmids are listed in Table S2 and all RNA are summarised in Table S3.

**Recombinant protein purification.**

LbuCas13a was expressed and purified as previously described (*65*). LbaCas13a and RfxCas13d were also expressed and purified using this method. Briefly, pGJK191 (LbuCas13a), pGJK189 (LbaCas13a) or pBKH011 (RfxCas13d) was transformed into *E. coli* BL21 (DE3) with antibiotic selection (ampicillin and kanamycin respectively). An overnight was prepared with a single colony in LB media. Expression was conducted using 800 mL of terrific broth (TB) in 2.5 L flasks under antibiotic selection. Cells were resuspended in lysis buffer (see above publication for all buffer compositions), before being lysed by sonication (30s on, 30s off for 4 - 6 cycles, amplitude 10kHz), and lysates were collected via high-speed centrifugation (32,000 x g for 45 min at 4°C). This lysate was applied to pre-equilibrated loose nickel affinity resin (Qiagen), washed with 10 CV of wash buffer (10 mM imidazole) and eluted fractions (300 mM imidazole) were dialysed overnight. Dialysis included TEV protease for LbuCas13a to cleave off the MBP and His-tag. Dialysed proteins were applied to an SP ion exchange column (Cytiva) and eluted with an increasing KCl gradient (0.2 - 1 M). Fractions of interest were concentrated using an Amicon concentrator (30K MWCO, Merck) and polished by size exclusion using a S200 16/600 or S200 10/300 Increase column on an AKTA pure (Cytiva). Protein was aliquoted, snap frozen with $LN_2$ and stored at -80°C until use.

**Cloning of mutants.**

All LbuCas13a and RfxCas13d mutant primers were designed using NEBuilder (NEB). Where multiple mutations were required, mutations were introduced using a 5′ overhang mismatch on the forward primer. PCR was performed using Q5 DNA polymerase (NEB) using pGJK191 or pBKH011 (for LbuCas13a and RfxCas13d respectively) as template. KLD enzyme mix was used as per protocol to generate circularized plasmids (NEB). All mutant plasmids were transformed into DH5α *E. coli* (for both), NEB turbo *E. coli* (some LbuCas13a mutants) or NEB 10-beta *E. coli* (some RfxCas13d mutants) onto LB agar plates under antibiotic selection (ampicillin and kanamycin respectively). The toxicity of some mutants required LB agar plates with 0.4% (v/v) D-glucose for repression. Mutants were confirmed using Nanopore whole plasmid sequencing (Plasmidsaurus).

**Expression and purification of mutants.**

For expression, all mutants were transformed into BL21(DE3) *E. coli* under appropriate antibiotic selection. A single colony was picked from each plate and used to inoculate 10 mL Overnight Express Instant TB autoinduction media (Merck) in 50 mL flasks. Cells were grown at 37°C, 180 rpm shaking for 8 hours before the temperature was dropped to 16°C for expression overnight (~16 hours). Cells were harvested by centrifugation (5000 x g, 4°C for 10 min). Pellets were frozen at -80°C until use, or snap frozen using dry ice before being immediately thawed for purification.

To lyse cells, B-PER complete (ThermoFisher Scientific) supplemented with 0.5 M NaCl, 10 mM imidazole and a protease inhibitor tablet (Roche, 1 per 50 mL of lysis buffer) was added to cell pellets (5 mL per 1 g of cell mass). Resuspended cells were incubated at room temperature for 30 min with gentle rocking. Clarified lysate was obtained by centrifugation at 21,000 × g, 4°C for 30 min. Lysate was added to ~200 µL of loose nickel affinity resin (Qiagen) pre-equilibrated with wash buffer (50 mM Tris pH 7, 0.5 M NaCl, 5% glycerol, 20 mM imidazole, 1mM TCEP) and incubated at 4°C for 1 hour with gentle agitation. Resin was washed with 20CV of wash buffer, twice, then eluted with 3CV of wash buffer containing 300 mM imidazole. Protein concentration was determined using Bradford assay, purity was assessed using SDS-PAGE and protein was snap frozen using liquid nitrogen and stored at -80°C until use. Mutant folding was assessed using Prometheus nanoDSF (NanoTemper) with a single replicate at 0.4 mg/mL.

**Isolation of total *E. coli* RNA.**

Competent BL21(DE3) *E. coli* (NEB) were streaked out on LB agar and grown overnight at 37°C. A single colony was used to inoculate a 25 mL starter culture of LB media and grown overnight at 37°C with 200 rpm shaking. The starter culture was added 1:100 (250 µL to 25 mL) to LB media and incubated at 37°C with 200 rpm shaking until an $OD_{600}$ of 1.0. Resulting cells (8 x $10^8$) were pelleted, and resuspended in 250 µL of lysozyme solution (10 mg/mL lysozyme, 10 mM Tris pH 7, 50 mM NaCl) and incubated at room temperature for 5 min. RNA lysis buffer was added 2:1 and centrifuged for 2 min at 16000 × g. Total *E. coli* RNA was purified using Monarch Total RNA Miniprep kit (NEB) as per manufacturer's instructions. RNA concentration and purity were assessed using $A_{260}/A_{280}$ and protein was snap frozen using $LN_2$ and stored at -80°C until use.

**RNA gel electrophoresis.**

RNA purity and Cas13 cleavage products were assessed using TBE-Urea-PAGE (16.5 cm x 28 cm). Either 12.5% (v/v) or 15% (v/v) polyacrylamide gels (29:1) were prepared with 8M Urea and 0.5X TBE, before being heated at 30 mA for 40 min. Samples were then resolved at 30 mA for ~50 min. Unlabeled RNA (10-50 ng) was stained for 10 min with SYBR Gold (ThermoFisher Scientific) before being imaged, while fluorescently labelled RNA (0.3-0.5 pmol) was immediately imaged. Both were imaged using Cy2 fluorescence (excitation 489 nm, emission 506 nm). All gels were imaged using a Typhoon laser scanner at 400 PMT with a pixel size of 100 µm (Cytiva).

**RNA sequencing.**

Preparation of the RNA library for sequencing was conducted similar to previously described (*25*). Prior to RNA sequencing, optimal aRNA concentrations were determined using TBE-Urea-PAGE (**Fig S2**). The concentration of aRNA was deemed optimal when minor depletion of rRNA was observed after 1 min, approximately half was depleted at 5 min and almost full depletion had occurred after 30 min. To prepare the cleaved samples, Cas13a:crRNA were combined in assay buffer (20 mM Tris pH 7, 30 mM KCl, 5 mM MgCl2, 2% v/v glycerol) with 100 nM of each (final concentration for the total assay) and left to incubate for 30 min at 37°C. Total *E. coli* RNA extract (10 µg per replicate, 3 replicates per time point) was added to the Cas13a:crRNA complex, and left to incubate at 37°C for 5 min. Either aRNA (10 nM for RfxCas13d and LbaCas13a, or 100 pM for LbuCas13a) or water (negative control) were added to start the reaction, with time points taken at 1, 5 and 30 min where the reaction was quenched with TRI-reagent (n=2 technical replicates for each enzyme with and without aRNA). RNA clean-up was conducted using the Direct-zol RNA microprep kit (Zymo Research: all following RNA clean-up steps were done using this kit) as per manufacturer's instructions. rRNA was depleted by oligo hybridisation with magnetic beads and ethanol precipitation using MICROBExpress Bacterial mRNA Enrichment Kit (Invitrogen). Resuspended RNA in water was fragmented with RNA fragmentation reagents (Invitrogen) at 70°C for 1 min before a second RNA clean-up was conducted. T4 PNK (NEB) treatment was conducted at 37 °C for 1 hour before a final RNA clean-up. All samples were sent to Azenta Life Sciences for library preparation including adapter ligation and cDNA synthesis (Vazyme Small RNA Preparation Kit), size selection (<200 nt) and Illumina PE150 sequencing (2x150bp single-read (SR) configuration with ~20.0 Mb reads PF data per sample) (LbuCas13a samples had PE100 configuration).

**RNA sequencing data analysis.**

RNA-seq reads were quality-filtered and trimmed for adapters using Trimmomatic v0.39. The trimmed reads for each library were aligned to the reference genome (RefSeq: NZ_CP053601.1) using Bowtie2 with unaligned reads excluded. The resulting alignments were sorted and indexed with SAMtools. Per-base read coverage was calculated with SAMtools depth. To investigate the differential coverage of all genes, the number of aligned reads per transcript (including rRNA, mRNA and tRNA) were quantified using Salmon from BAM files generated by Bowtie2. DESeq2 was then used to calculate fold changes and pairwise q-values. Fold change was defined as the coverage level in the Cas13 active samples divided by that in the inactive samples. Transcripts with an adjusted P-value $\leq 0.05$ and log2(Fold Change) $\geq 2$ were considered significant. The results were visualized as volcano plots displaying log2(Fold Change) versus log10(adjusted P-value). The precise as genomic positions of 5′ ends of aligned reads were extracted using BEDTools and deemed to be cut sites created by activated Cas13. To precisely locate the cutting sites, the 5′ end was defined as the alignment end for both positive and negative strand reads, instead of using the default alignment start for positive strands. The 3′ end was defined as the alignment start for both positive and negative strand reads. Per-position coverage for 5′ or 3′ ends were calculated using BEDTools genomecov. Differential coverage analysis was performed using DESeq2. Raw count data were normalized by estimating size factors based on the total number of aligned reads to account for differences in library size. Fold changes at individual sites were calculated as the ratio of coverage levels in Cas13 active vs inactive samples for each sample condition.

Signature Cas13 cleavage motifs were determined by extracting the local sequences of all cut sites and tallying (+1) the occurrence of nucleotides at each position weighted to the associated cut site fold change (+1x$\log_2$FC). tRNA cut sites were determined by aligning each gene according to consensus tRNA data (tRNAViz (*66*)) to account for sequence differences interfering with alignment of cut sites at analogous positions.

**Assays to determine cleavage of rRNA bound to ribosomes.**

To determine if rRNA could be cleaved when associated with ribosomal proteins (**Fig. S3**), we complexed Cas13:crRNA in assay buffer (final concentration in assay 25 mM KCl, 2.5 mM MgCl2, 2% (v/v) glycerol, 10 mM Tris-HCl, pH 7.0) for 15 mins at 37°C. The substrate (ribosomal bound rRNA: NEB P0763S) was then combined with RNase-free water or aRNA, before the Cas13:crRNA complex was added at a ratio of 1:3 substrate to complex. The reaction was incubated at 37°C for 60 min for the negative control (no aRNA) and 1, 3, 10, 30, 60, 120 and 240 mins for experimental conditions. In all cases the reaction was quenched with 2X RNA loading dye (30 mM EDTA, 0.2% (w/v) SDS, 200 µg/mL heparin, 0.5% (w/v) bromophenol blue in formamide). Samples were analysed using RNA gel electrophoresis with SYBR Gold (ThermoFisher Scientific) as above.

**RNA cleavage or competition assays.**

The activity of Cas13 was determined by TBE-Urea-PAGE (using unlabeled or 5′ 6-FAM labelled RNA) or quantified by 384-well plate fluorescence cleavage assays (for 5′ 6-FAM labelled RNA only). Prior to all assays, Cas13 was complexed with crRNA (1:1) in assay buffer (20 mM Tris pH 7, 30 mM KCl, 5 mM MgCl2, 2% v/v glycerol) and incubated for 30 min at 37°C. The final concentration of Cas13 and crRNA in all assays was 100 nM unless stated otherwise.

***TBE-Urea-PAGE cleavage assays.***

For the TBE-Urea-PAGE, the reaction was started by the addition of aRNA and 5′ 6-FAM labelled substrate RNA 1:3 to the RNAP complex at 37°C. Time points were taken at 1, 3, 10, 30, and 60 min where the reaction was quenched with 2X RNA loading dye (30 mM EDTA, 0.2% (w/v) SDS, 200 µg/mL heparin, 0.5% (w/v) bromophenol blue in formamide). Samples labelled with 0 min were incubated with Cas13a and crRNA for 60 mins but were not incubated with aRNA (hence 0 min). Gels were then run as described above with 0.3 pmol of fluorophore for labelled RNA (0.5 pmol for rRWC009 and rRWC043 (**Fig. S9**) or ~10 ng for unlabeled RNA loaded per lane. Resulting gels were analyzed by densitometry analysis using ImageQuant (Cytiva). The percentage cleaved (%cleavage) of substrates was calculated by the intensity of full-length substrates relative to the entire lane. Mean % cleaved at time point 0 min was used to define 0% for normalization (where 100%=100) per substrate. Normalized data had fitted non-linear regression curves with plateaus set at >80% with associated K (rates) per replicate (n=4) to determine rate of substrate cleavage.

***Plate based fluorescence cleavage assays.***

The 384-well plate assays were conducted as described previously (*65*). Briefly, after complexing Cas13:crRNA, the aRNA and 5′ 6-FAM labelled substrate RNA were combined in assay buffer and 5 µL was added to a 384-well plate. The aRNA concentrations were variable (see **Table S4**), but the substrate RNA had a final concentration of 400 nM. To start the reaction, 15 µL of the Cas13:crRNA complex was added for a final concentration of 100 nM of each. Cleaved fluorescent RNA products were measured on a ClarioSTAR plus plate reader (BMG Labtech) for 30 - 60 min at 37°C, with readings taken at most every 30 s. All assay data analysis was performed using GraphPad Prism (10.4.2). Observed velocity was calculated by measuring the initial linear phase of the assay.

***Plate based fluorescence competition assays.***

After complexing Cas13:crRNA, a dilution series of unlabeled RNA competitors (see **Table S3**) were combined with solutions of aRNA and 5′ 6-FAM labelled substrate (rGJK123) and 5 µL was added to a 384-well plate. The aRNA concentration was 2 pM (unless otherwise specified), the substrate RNA had a final concentration of 500 nM, and the competitor spanned concentrations from 10 µM to ~2 nM. Cleaved fluorescent RNA products were measured on a ClarioSTAR plus plate reader (BMG Labtech) for 30 min at 37°C, with readings taken every 30s. All assay data analysis was performed using GraphPad Prism (10.4.2). Observed velocity was calculated by measuring the initial linear phase of the assay and IC50 values calculated per replicate by Inhibitor Vs Response curves (bottom = 27, determined from -aRNA samples) across n=3 replicates unless otherwise stated.

**Cryo-electron microscopy.**

***Sample preparation***

LbuCas13a and crRNA (rGJK119) were combined at a 1:1.2 molar ratio, and incubated at RT for 12 min. Activator RNA (rGJK120) was then added at a 1.4 molar ratio (relative to LbuCas13a) and incubated for a further 3 min. The complex was developed over a Superdex S200 3.2/300 column (Cytiva) equilibrated in 20 mM Tris pH 7, 50 mM KCl, 5 mM MgCl2 and 2% (v/v) glycerol (AKTAmicro, Cytiva). The peak containing the LbuCas13a ternary complex was used for grid preparation at ~8 µM (as determined by $A_{280}$). For addition of the tRNA ASL substrate RNA (rRWC039: $tRNA^{Lys(dUdUU)}$ and rRWC60: $tRNA^{Asp(fGUC)}$), 0.25 µL (1 mM) was combined with 3.3 µL of the LbuCas13a ternary complex and incubated for 1 min at RT. A volume of 3.5 µL was applied to a freshly positive glow discharged (30s at 30 mA with amylamine) UltrAUfoil R 1.2/1.3 Au 300 grid (Quantifoil GmbH). Grids were blotted (-3 blot force, 3s blot time) with a Vitrobot Mark IV (ThermoFisher Scientific) set at 4°C, 100% humidity before plunging into liquid ethane.

***Data collection.***

Initial grid screening was performed using a FEI Talos Arctica with 200 kV accelerated voltage equipped with a Falcon 3EC direct electron detector (ThermoFisher Scientific). Data collection was performed using a G1 Titan Krios (ThermoFisher Scientific) equipped with a Schottky field emission gun (S-FEG) operated at an accelerating voltage of 300 kV. A C2 condenser aperture of 50 µm was used and images were captured using a K3 direct electron detector (GATAN) positioned post a BioQuantum energy filter (GATAN). Automated data collection was performed using EPU

(ThermoFisher Scientific) and a nominal magnification of 105kX in EFTEM mode resulting in a pixel size of 0.82 Å/pixel. Zero loss filtering was done using an energy filter slit width of 10eV. Movies were fractionated into 60 frames with a total dose of 60 e/Å2. The defocus range was set between -0.5 and -1.6 µm.

***Data processing.***

All datasets were subjected to Patch Motion Correction disregarding the initial frame and Patch CTF Estimation in cryoSPARC 4.5.3 (*66*) except for LbuCas13a:crRNA or LbuCas13a:crRNA:aRNA data (**Fig. S21**), which were subjected to MotionCor2 (*67*) and CTFFIND-4.1 (*68*) in RELION 4.0.0 (*69*). Ideal particles (around 200-1000) were manually picked from a micrograph subset and used to train an initial Topaz model (*70*). Poor-quality Topaz picks were filtered out using 2D classification with cryoSPARC.

For LbuCas13a:crRNA and LbuCas13a:crRNA:aRNA data without substrate, filtered particle picks were reconstructed into an initial ab-initio then consensus high-resolution refined reconstruction using Non-Uniform Refinement (NU-Refine (*71*)) (cryoSPARC). LbuCas13a:crRNA:aRNA complexes were a sub-population compared to LbuCas13a:crRNA and were separated by 3D classification (5 classes, 6 Å target resolution). After Bayesian polishing (*72*), (reimported to RELION) and local CTF refinement, final LbuCas13a:crRNA and LbuCas13a:crRNA:aRNA NU-refined reconstructions were obtained for model building. After gaussian smoothing (2σ) (ChimeraX (*63*)), these reconstructions were also used as templates for heterogeneous refinement in following datasets (**Fig. S22-24**).

For remaining datasets, the filtered particle picks were used to train an improved Topaz model. After micrograph curation, particle picks were subjected to heterogeneous refinement (cryoSPARC) using both +/-aRNA templates mentioned previously and noise templates. Once particle stacks produced resolved protein density from a separate NU-refinement, a mask surrounding the HEPN active site was used for focused 3D classification to separate particles with clear substrate density (3-5 classes, 10 Å target resolution). Final reconstructions were acquired after local CTF refinement and reference-based motion correction.

For data involving $tRNA^{Lys(dUdUU)}$ substrate (rRWC039), the final minus (-) aRNA reconstruction were from particle stacks of a single dataset whereas the final plus (+) aRNA reconstruction combined particle stacks from separate datasets (**Fig. S22-23**). For data involving $tRNA^{Asp(fGUC)}$ substrate (rRWC060), particles were imported to RELION (5.0.0) for focused 3D classification to resolve density surrounding the scissile phosphate before reimporting back to cryoSPARC for final reconstruction (**Fig. S24**). Tau values for classification were selected to remove particles lacking density in specified areas (very large ~10000) or to resolve structural differences (middling ~3500).

Focused 3D Variability Analyses (*73*) were carried out for all particles in the final LbuCas13a:crRNA:aRNA:$tRNA^{Asp}$ (**Movie S1**) and LbuCas13a:crRNA:$tRNA^{Lys}$ (**Movie S2**) reconstructions. Whole particle masks generated from NU-refinement were trimmed to include the HEPN active site, HEPN accessory elements, and the substrate entirety. Custom parameters used were to solve a single (1) mode at 5 Å resolution filter.

***Model building.***

For the apo-HEPN structures (LbuCas13a:crRNA and LbuCas13a:crRNA:aRNA), PDB 5XWY and 5XWP were used as initial input (**Fig. S25A,B**) (*18*). For the binary $tRNA^{Lys}$ structure (LbuCas13a:crRNA:$tRNA^{Lys}$), PDB 5XWY and 6UGG (unmodified $tRNA^{Asp}$) were used as initial input (**Fig. S25C**) (*18, 61*). For the ternary $tRNA^{Asp}$ structures (LbuCas13a:crRNA:aRNA:$tRNA^{Asp}$), the LbuCas13a wild-type ternary structure was used as initial input (**Fig. S25E**). The ternary $tRNA^{Asp}$ structure (state 1) was then used as input for the ternary $tRNA^{Lys}$ structure (LbuCas13a:crRNA:aRNA:$tRNA^{Lys}$) (**Fig. S25D**). All models were placed in the cryoEM density using ChimeraX (*63*). All structures were refined using Phenix (1.21.2) (*74*) real space refinement, with manual corrections using Coot (0.9.8.93) against EMReady (*75*) post-processed maps for ambiguous density. Molprobity (*76*) was used for model inspection, and all model figures were produced using ChimeraX (1.9) (*63*).

**Mass spectrometry for LbuCas13a product identification.**

LbuCas13a was complexed with crRNA (1:1 ratio) in Cas13 assay buffer (20 mM Tris pH 7, 30 mM KCl, 5 mM MgCl2, and 2% v/v glycerol), and incubated at room temperature for 10 min. The substrate was added (rBKH020) at the same time as aRNA, for a final concentration of LbuCas13a of 100 nM, crRNA 100 nM, 1 nM aRNA and 4 µM substrate. For RNase T1, 0.4 mU/uL (ThermoFisher Scientific) of enzyme was added to 4 µM amount of substrate in Cas13 assay buffer. Both reactions were incubated for 10 min before being quenched with equal volume 100% acetonitrile (to 50%). Additionally, substrate and product (rBKH021) standards were made in water to 4 µM and combined with equal volume 100% acetonitrile (to 50%).

For LC-MS/MS sample preparation, the sample was dried down in a SpeedVac concentrator (Labconco) and resuspended in 10 µL of 80% acetonitrile/20% RNase-free water. 1 µL was injected for LC-MS/MS system analysis. Chromatographic separation of the sample was performed on a Vanquish Flex UHPLC system (ThermoFisher Scientific), on a GTxResolve Premier BEH Amide column (2.1 × 100 mm, 1.7µm, Waters) at 80°C utilizing a 8 min 90–30% gradient of solvent A (25mM ammonium acetate, 10% Acetonitrile) and solvent B (25mM ammonium acetate, 80% Acetonitrile) at a flow rate 300 µl/min. Mass spectrometry analysis was performed on an Orbitrap Eclipse Tribrid mass spectrometer (ThermoFisher Scientific) in negative electrospray ionization mode with the default source settings. Full mass data was acquired at a resolution of 120000 (FWHM) at m/z 200. Data was reviewed in FreeStyle 1.8 (ThermoFisher Scientific) and deconvoluted utilizing Xtract (ThermoFisher Scientific), with the following parameters: Charge Range: 2-10; Analyzer Type: OT; Relative Abundance Threshold: 0%; Isotope Table: Nucleotide; Negative Charge; Minimum Number Detected Charge: 3. Data are available via ProteomeXchange with identifier PXD065184 (*64*).

**HEPN RNase AlphaFold modelling**

For analysis of potential accessory elements and residues that may determine substrate preference for RfxCas13d, a structural model was produced using the AlphaFold3 server (*77*). To obtain the ternary state, the full-length protein (WP_075424065.1), crRNA (rBKH022), and aRNA (rGJK074) were used as input. Five models were produced, and the top model (based on pTM score) was used for analysis (**Fig. S28**). The top model had a reported pTM was 0.85 and iPTM 0.49. To generate hypothetical substrate-bound states of a non-Cas13 HEPN, Human RNase L was folded as a dimer with 2-5A ligand and a single tRNA Proline ACN stem loop using the AlphaFold3 server (*77*). The top candidate with splayed bases around conserved catalytic residues was used for visualization in **Figure 5C** and mutated in **Figure 5D** with HEPN-aligned Ire1 (ChimeraX).

### Supplementary Figures

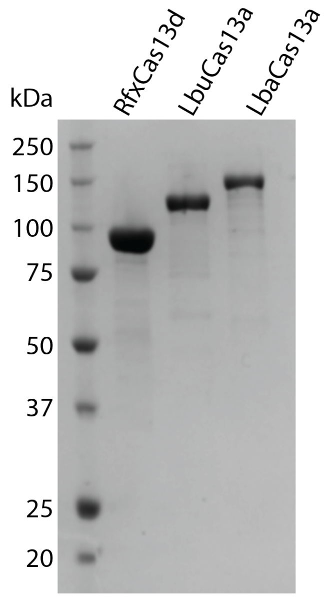


**Figure S1. Purified recombinant Cas13 enzymes.** From left to right, *Ruminococcus flavefaciens* (Rfx) Cas13d (102 kDa), *Leptotrichia buccalis* (Lbu) Cas13a (140 kDa) and *Lachnospiraceae bacterium* (Lba) Cas13a (170 kDa).

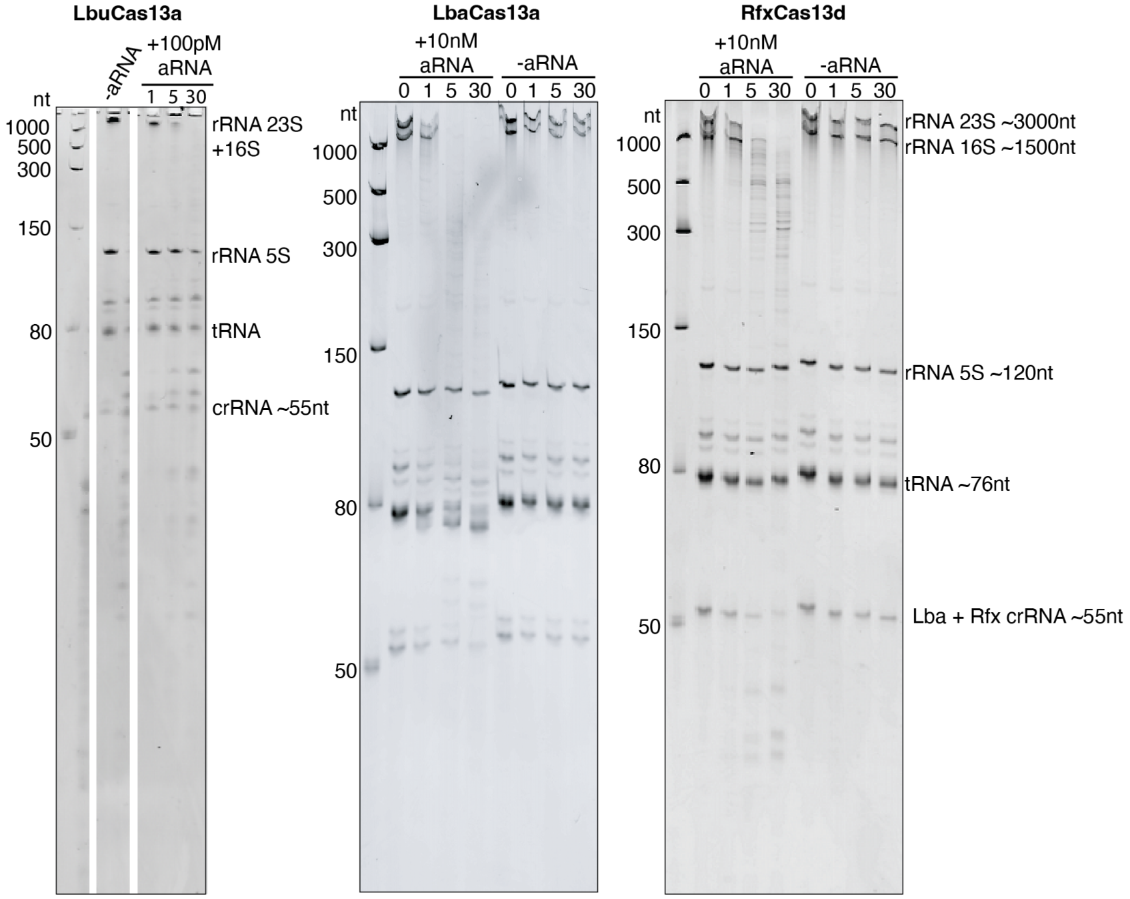


**Figure S2. TBE-UREA polyacrylamide gels showing degradation of isolated *E. coli* RNA by various Cas13 enzymes.** All numbers above the gels refer to mins. From left to right, total *E. coli* RNA degradation by LbuCas13a, LbaCas13a and RfxCas13d. For all panels a ssRNA ladder is on the left of the gel, with sizes indicated in nt. For LbuCas13a, a -aRNA control is included, and +100pM aRNA samples are taken at 1, 5 and 30 mins. For LbaCas13a and RfxCas13d, samples are taken + and - 10 nM aRNA at 0, 1, 5 and 30 mins.To the right of the gels, the rRNA and tRNA bands are labelled, as well as the LbaCas13a and RfxCas13b crRNA as there are visible in the gel at these high concentrations.

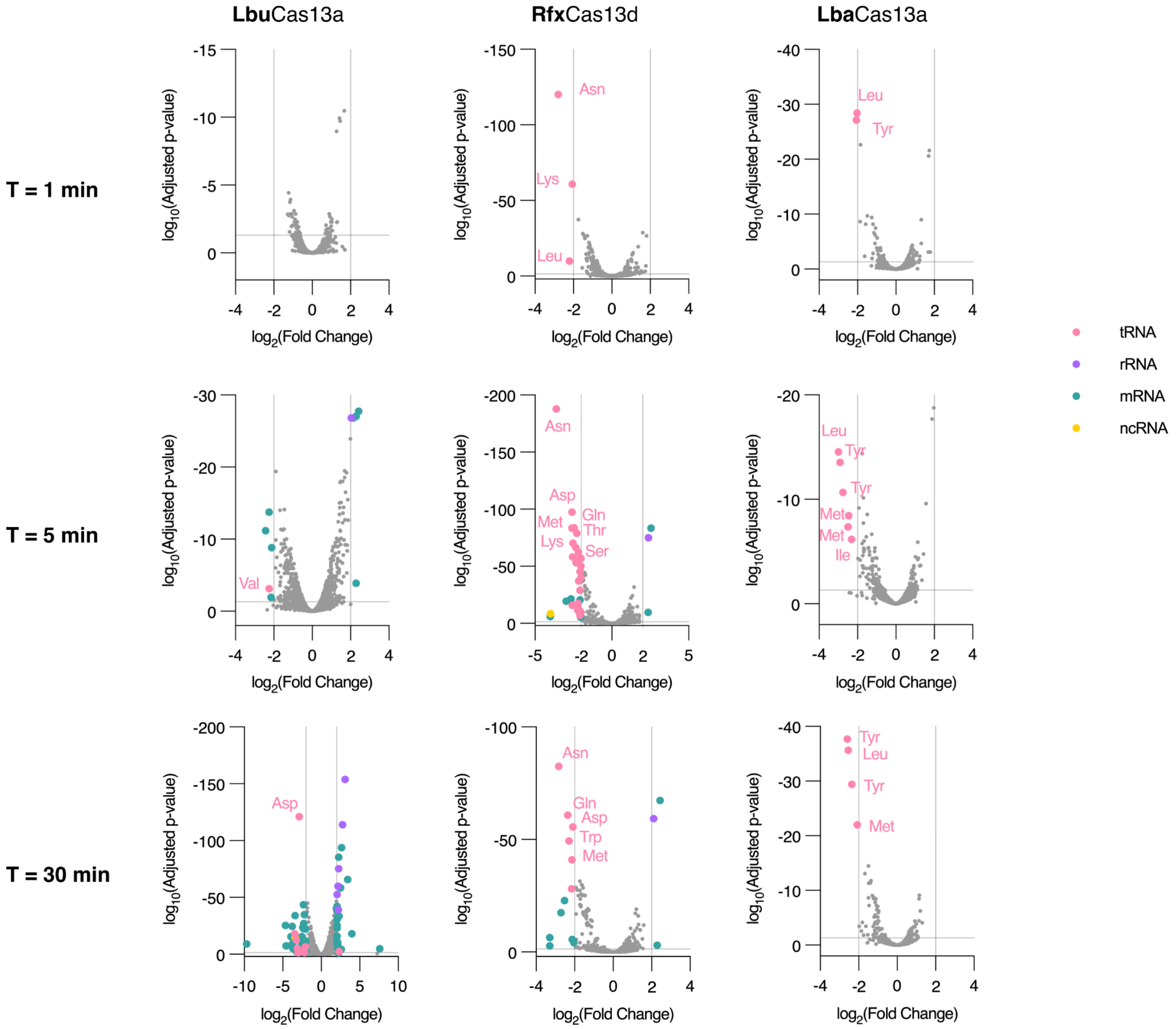


**Figure S3. Differential transcriptome coverage of *E. coli* RNA after cleavage by Cas13.** RNA Volcano plots show differential transcriptome coverage of total *E. coli* RNA by LbuCas13a (left), RfxCas13d (middle) and LbaCas13a (right) activity after 1 (top), 5 (middle), and 30 min (bottom) of reaction time (n=2). Thresholds: adj. P = 0.05 and $|log_2(FC)| = 2$. tRNA genes surpassing these thresholds are marked in pink and labelled by their associated amino acid.

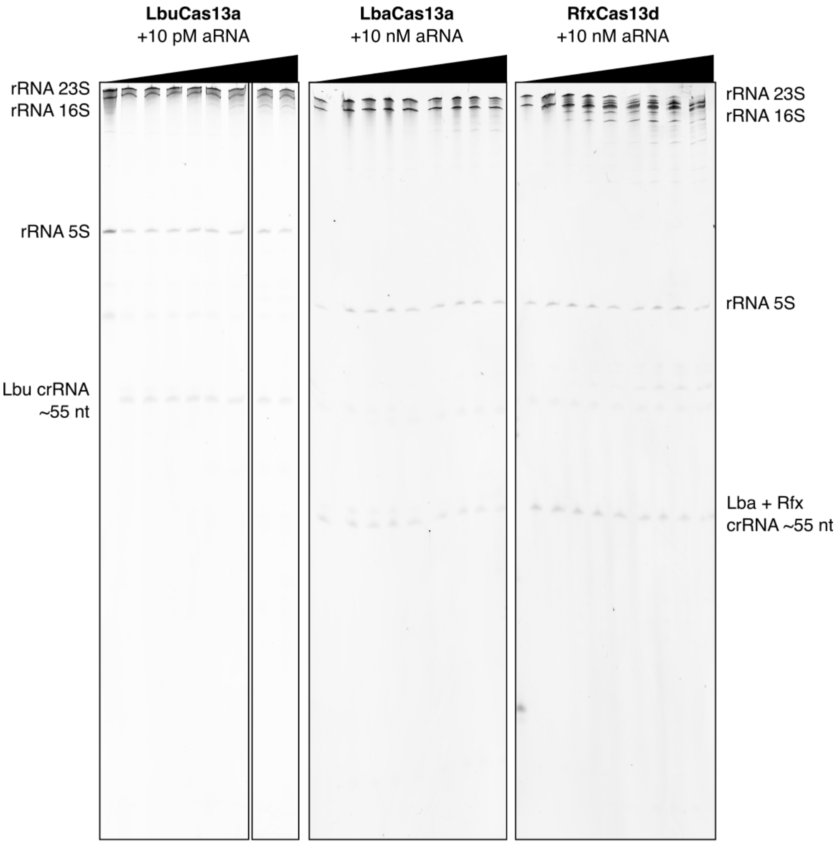


**Figure S4. rRNA within complete ribosomal proteins is protected from degradation by various Cas13 enzymes.** SYBR-gold-stained gel images of ribosome samples incubated with various activated Cas13 enzymes (n=1). aRNA concentrations used for each enzyme labelled above. Triangles above gels indicate a full reaction time course (0, 1, 3, 10, 30, 60, 120, and 240 minutes).

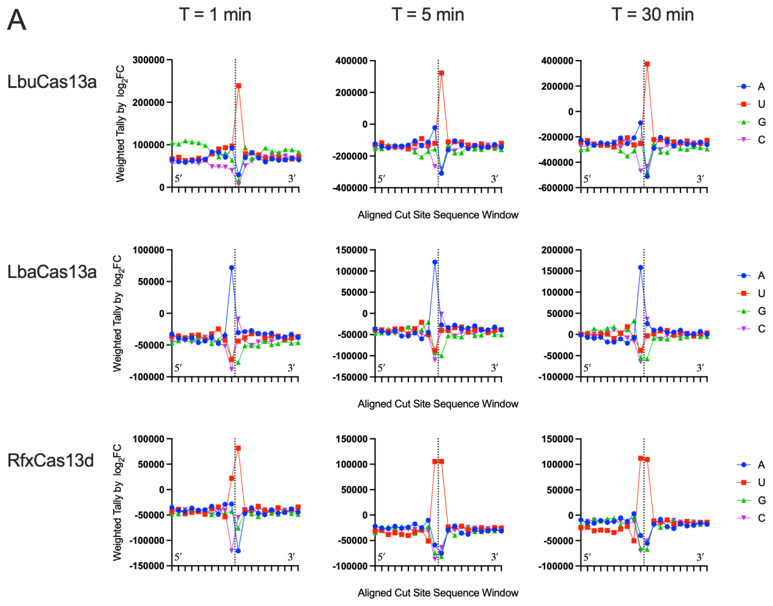


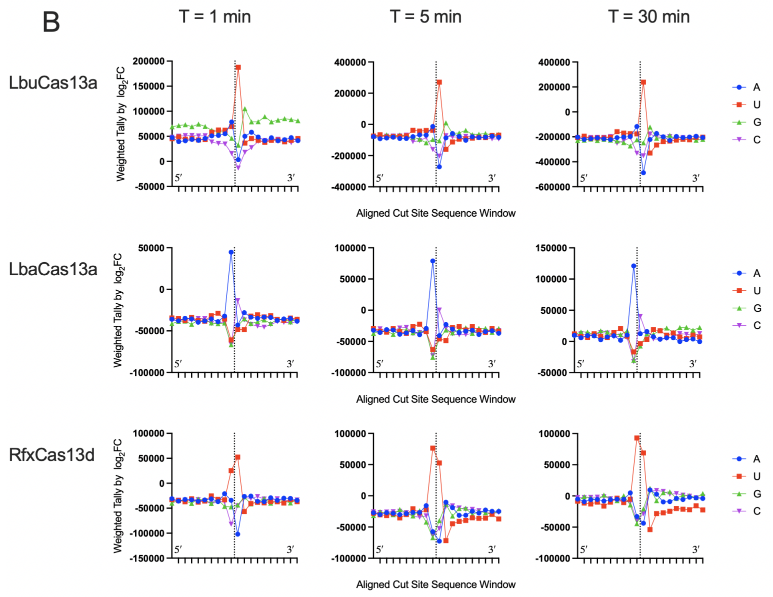


**Figure S5. Cas13 cleavage signatures from total *E. coli* RNA cut sites.** $log_2$FC-weighted tally of nucleotide type (+1 x $log_2$FC) along local cut site sequence demonstrate Cas13 sequence preferences with enriched nucleotide type (A, U, G, or C) at certain positions. Local 9 nt either side from scissile phosphate (dotted line) extracted. Cut sites detected by enrichment of 5′ ends (**A**) or 3′ ends (**B**). Within each panel, LbuCas13a (top) cuts N$^{\downarrow}$U, LbaCas13a (right) cuts A$^{\downarrow}$N, and RfxCas13d (middle) cuts U$^{\downarrow}$U across all the time points (left-right: 1, 5, 30 min).

348

LbuCas13a. Analysis of 5' ends in tRNA

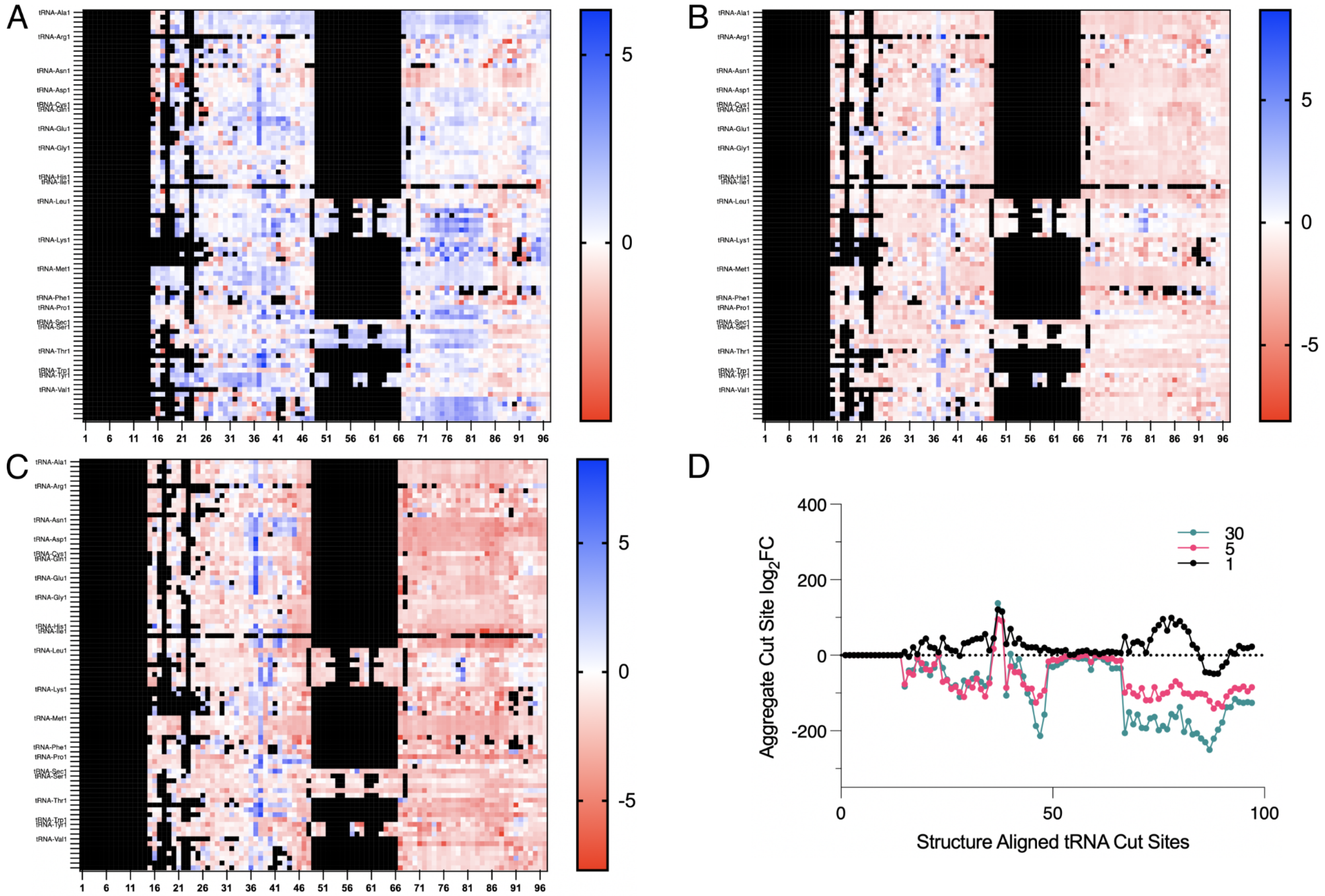

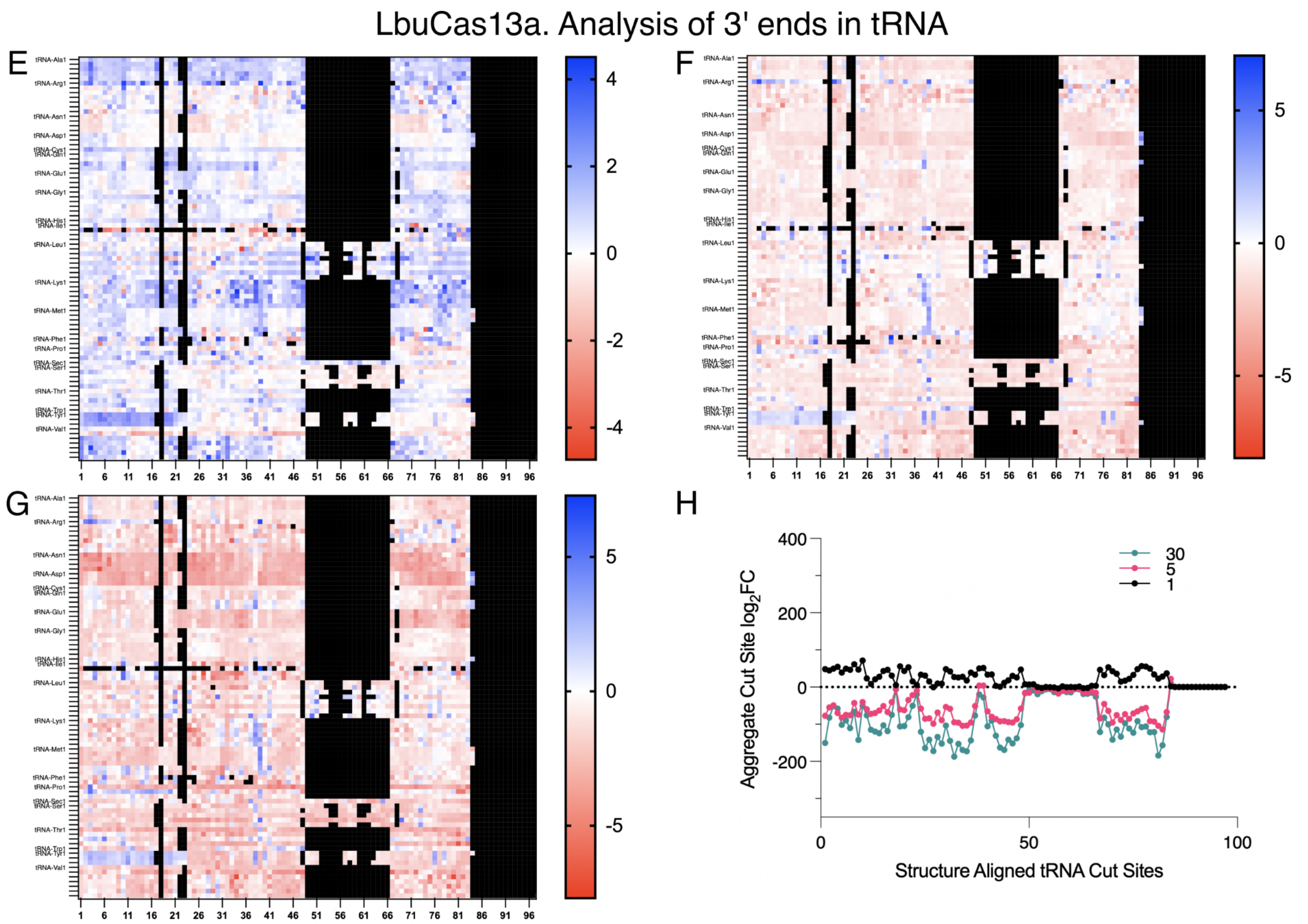


**Figure S6. tRNA cut sites by LbuCas13a occur at the anticodon.** Analyses of differential 5′ (**A-D**) and 3′ (**E-H**) ends in tRNA. Heatmaps of aligned tRNA sites (x-axis) for each tRNA gene (y-axis) (if unlabeled, same type as above) colored by $log_2FC$ at T=1 (A, E), 5 (B, F), and 30 minutes (**C, G**). Larger fold change indicates an enrichment of 5′ or 3′ ends from RNase activity. (**D, H**) Aggregate FC of aligned tRNA cut sites by structure (T=1 black, T=5 teal, T=30 pink) reveal anticodon cut sites are enriched across the tRNA cohort.

RfxCas13d. Analysis of 5' ends in tRNA

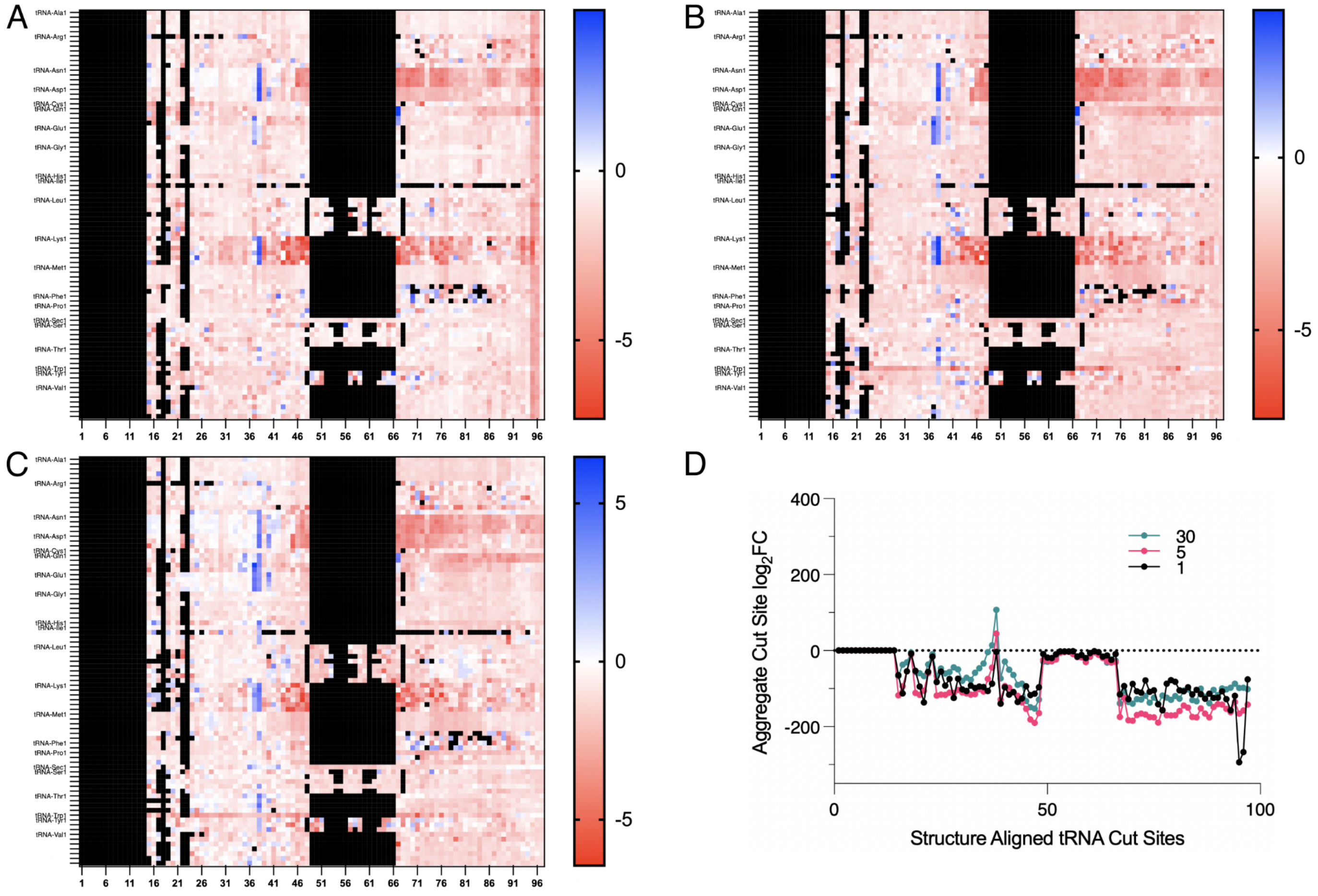

354

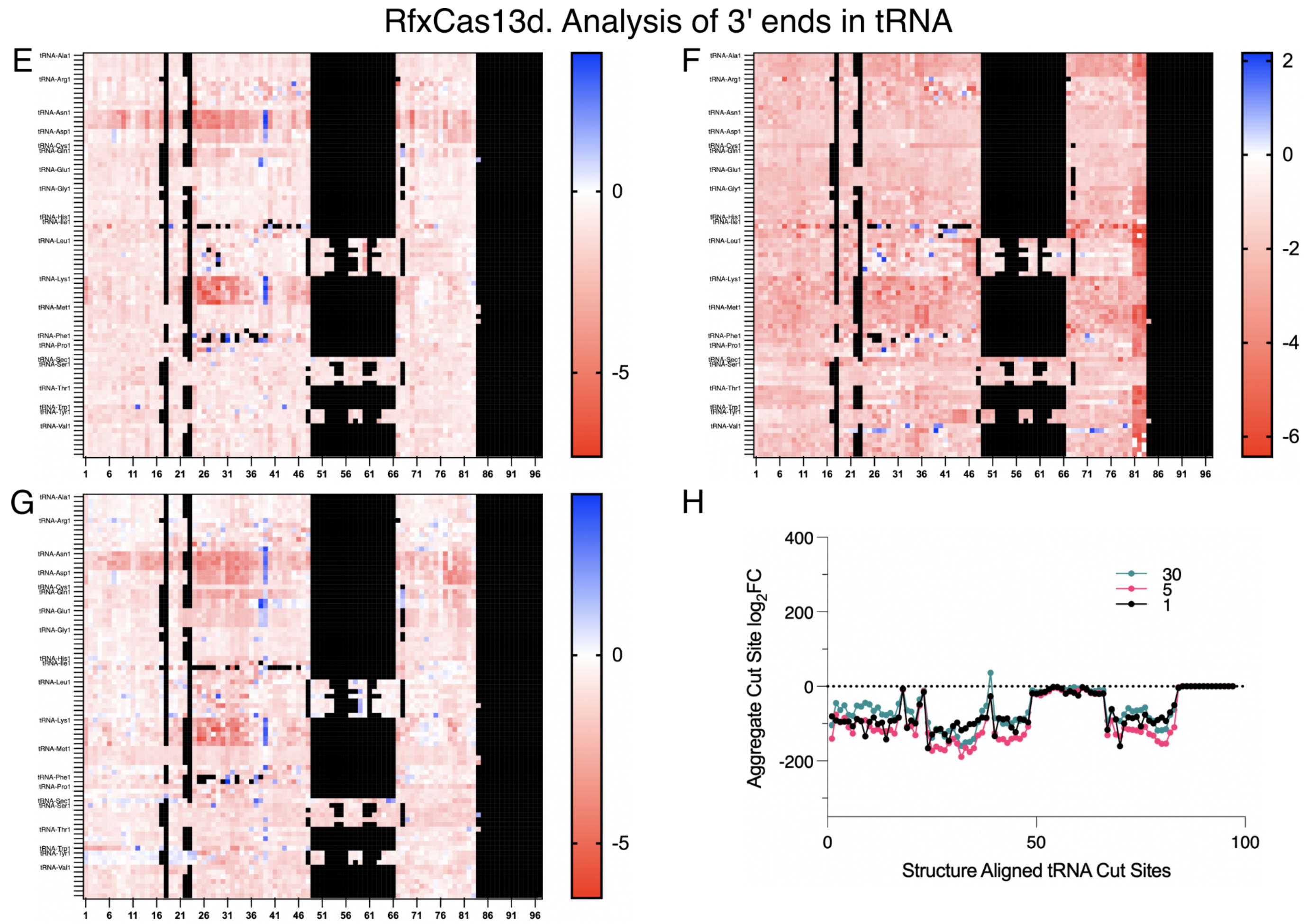


**Figure S7. tRNA cut sites by RfxCas13d occur at the anticodon.** Analyses of differential 5′ (**A-D**) and 3′ (**E-H**) ends in tRNA. 5′ ends between 1-14 and 3′ ends between 84-97 are undetectable. Heatmaps of aligned tRNA sites (x-axis) for each tRNA gene (y-axis) (if unlabeled, same type as above) colored by $log_2$FC at T=1 (A, E), 5 (B, F), and 30 minutes (**C, G**). Larger fold change indicates an enrichment of 5′ or 3′ ends from RNase activity. (**D, H**) Aggregate FC of aligned tRNA cut sites by structure (T=1 black, T=5 teal, T=30 pink) reveal anticodon cut sites are enriched across the tRNA cohort.

LbaCas13a. Analysis of 5' ends in tRNA

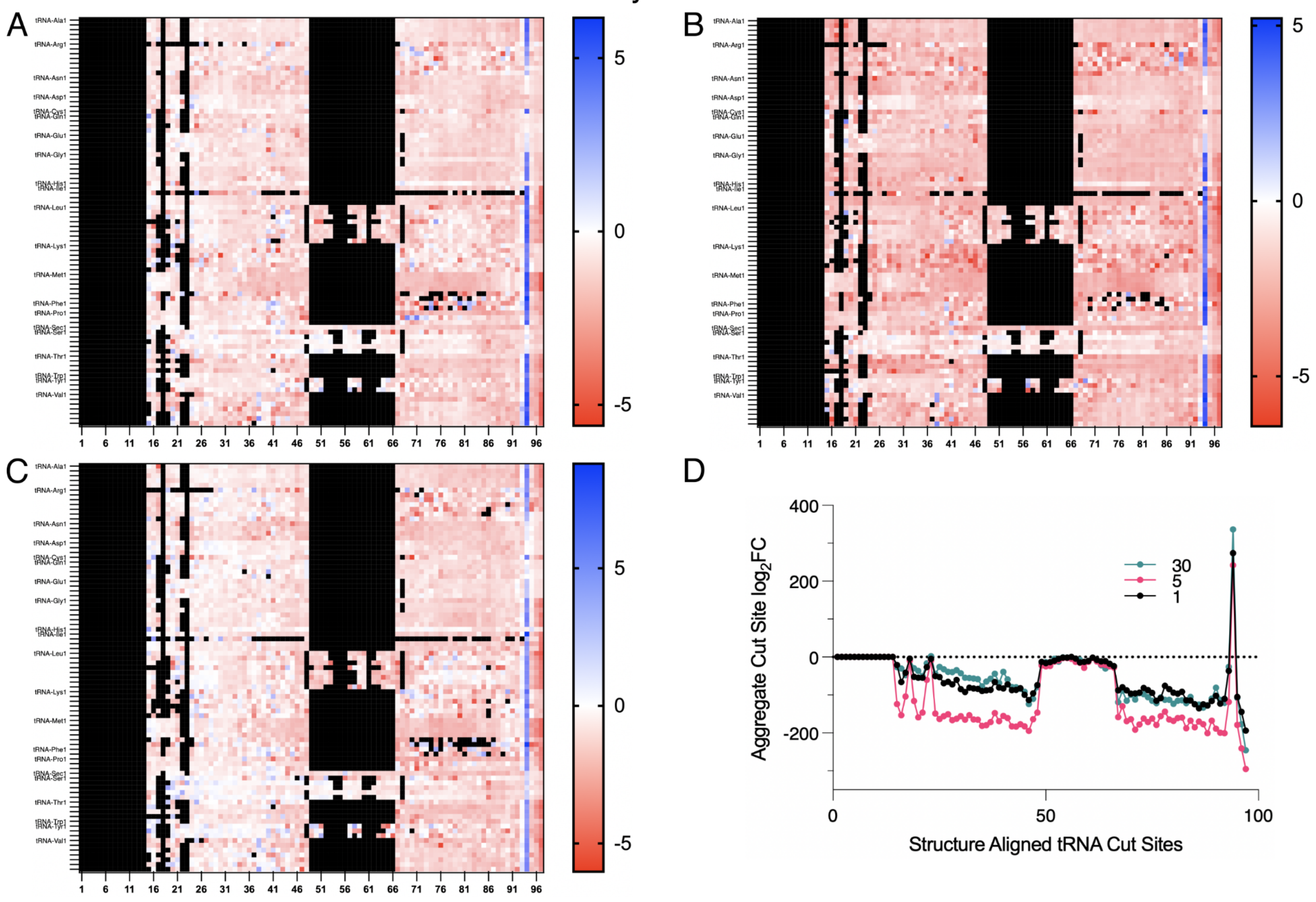


360

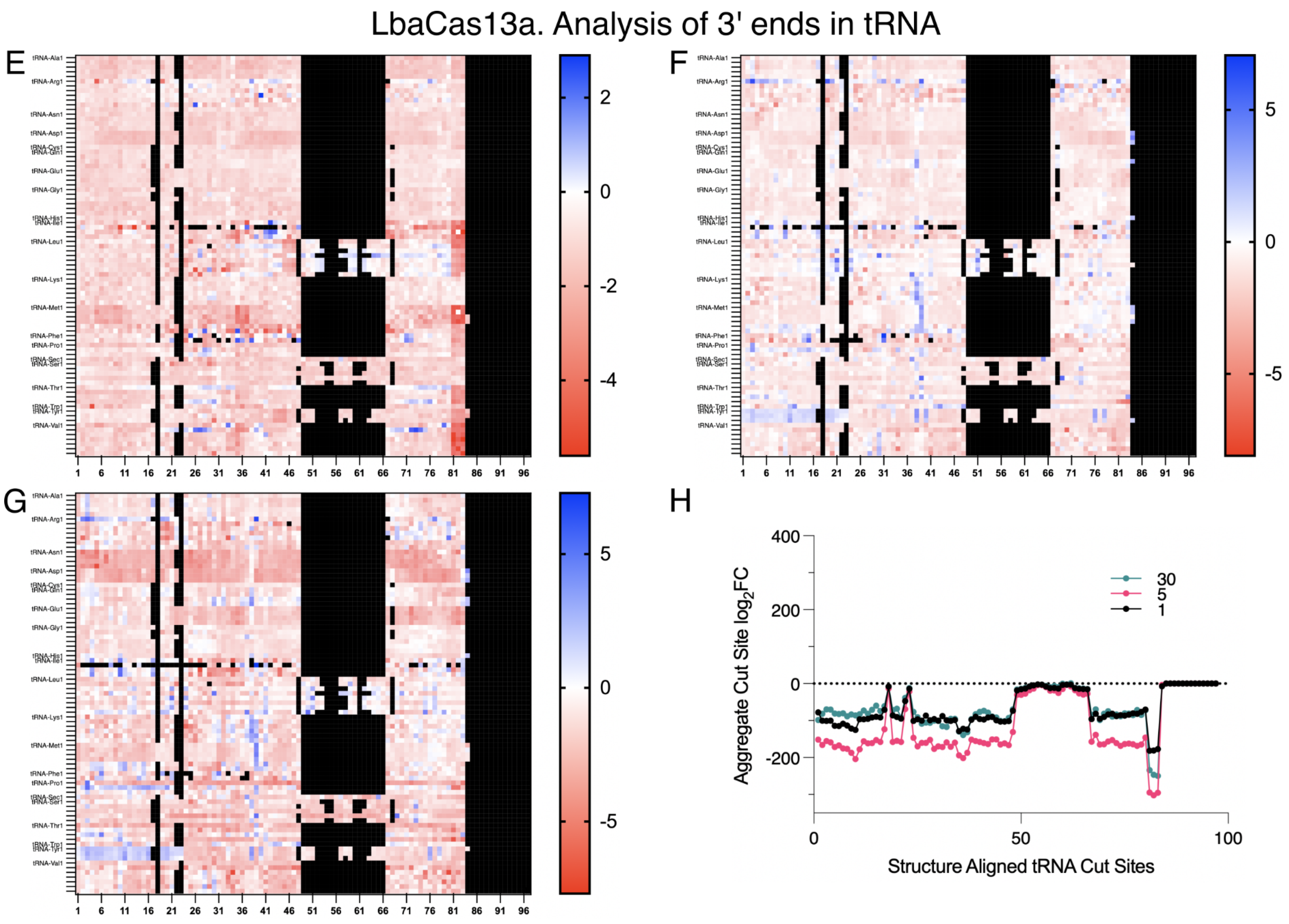


**Figure S8. tRNA cut sites by LbaCas13a occur at the CCA-tail.** Analyses of differential 5′ (**A-D**) and 3′ (**E-H**) ends in tRNA. Heatmaps of aligned tRNA sites (x-axis) for each tRNA gene (y-axis) (if unlabeled, same type as above) colored by $log_2FC$ at T=1 (A, E), 5 (**B, F**), and 30 minutes (**C, G**). Larger fold change indicates an enrichment of 5′ or 3′ ends from RNase activity. (**D, H**) Aggregate FC of aligned tRNA cut sites by structure (T=1 black, T=5 teal, T=30 pink) reveal acceptor stem A↓CCA cut sites are enriched across the tRNA cohort.

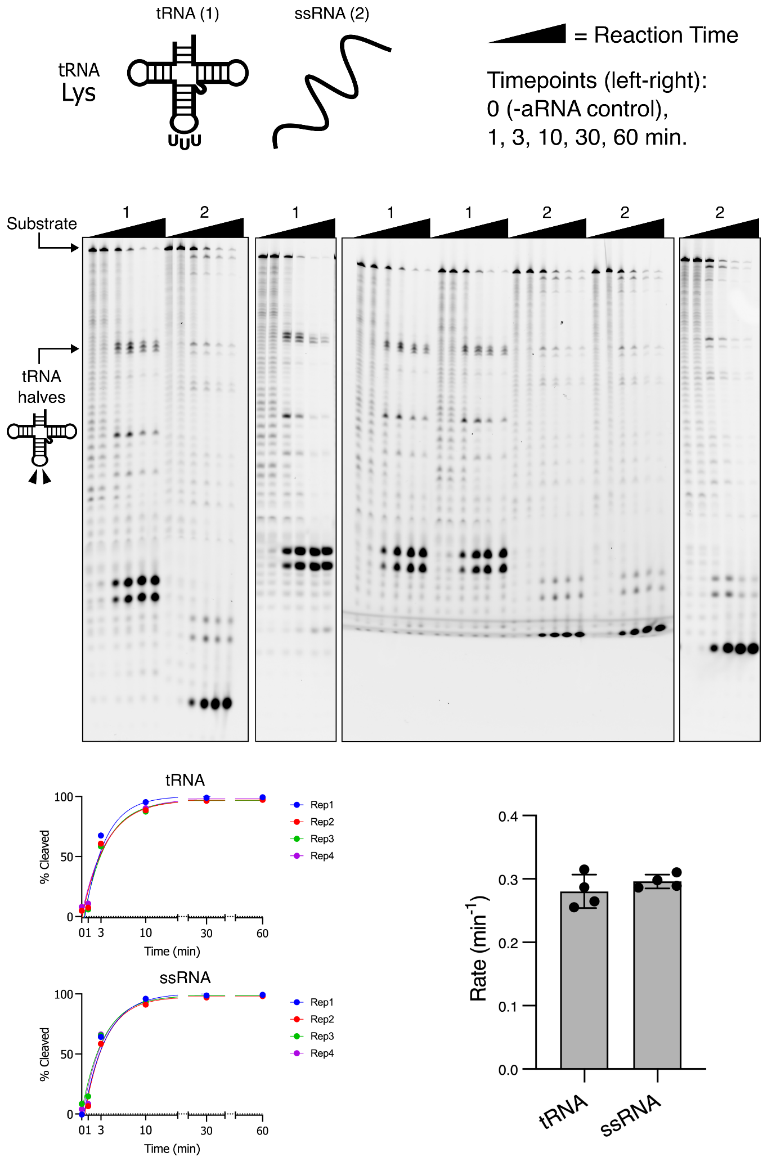


**Figure S9. Cleavage profiles of chemically synthesized *Leptotrichia buccalis* tRNA$^{Lys(UUU)}$ and a nucleotide-scrambled ssRNA derivative. (Top)** Gel images showing cleavage by LbuCas13a of 5′ 6-FAM tRNA$^{Lys}$ without modification (1, rRWC009) and 5′ 6-FAM ssRNA with identical nucleotide content but different sequence (2, rRWC043). Triangles above gels indicate a full reaction time course with a number to indicate the substrate. 5′ tRNA halves are indicated on the left. **(Bottom left)** Plots of %Cleaved relative to full-length substrate for both tRNA and ssRNA substrates and fitted non-linear regression curves per replicate (n=4 technical replicates). **(Bottom right)** Associated rate constants (K) of curve fits represented as mean +/- SD.

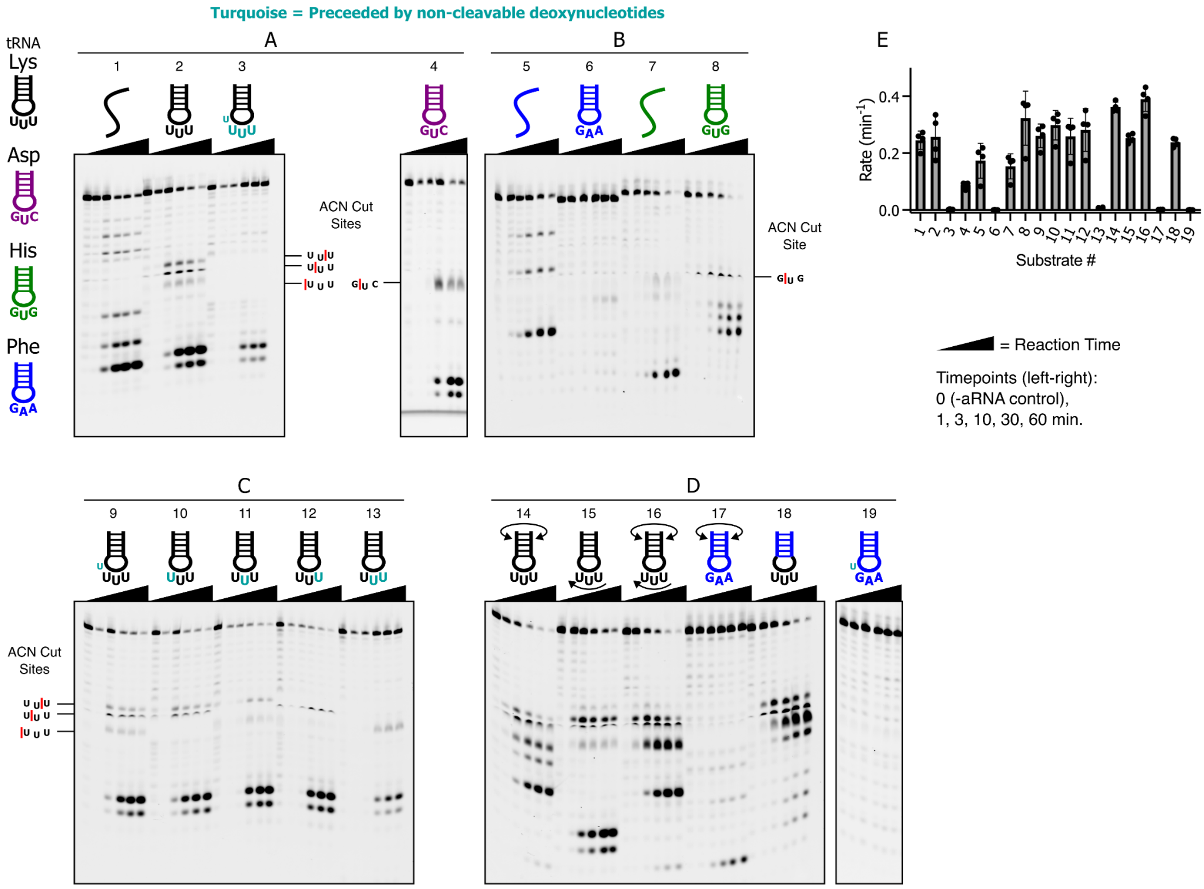


**Figure S10. Cleavage profiles of chemically synthesised tRNA-derived small substrates.** Representative gel images showing cleavage by LbuCas13a of 5′ 6-FAM RNA substrates. Triangles above gels indicate a full reaction time course with a number to indicate the 5′ 6-FAM substrate. Substrates are derived from tRNA ACN stem loops (ASL) from *Leptotrichia buccalis*. Substrates are numbered in order: (**A**) scrambled ASL$^{Lys(UUU)}$ (1, rRWC024), ASL$^{Lys(UUU)}$ (2, rRWC023), ASL$^{Lys(dUdUU)}$ with 4 deoxy-blocked cut sites (3, rRWC033), and ASL$^{Asp(GUC)}$ (4, rRWC063): (**B**) scrambled ASL$^{Phe(GAA)}$ (5, rRWC027), ASL$^{Phe(GAA)}$ (6, rRWC026), scrambled ASL$^{His(GUG)}$ (7, rRWC048), ASL$^{His(GUG)}$ (8, rRWC047): (**C**) a series of ASL$^{Lys(UUU)}$ substrates with single deoxy-blocked cut sites (9, rRWC029) (10, rRWC030) (11, rRWC031) (12, rRWC032) or 2 (13, rRWC035): (**D**) ASL$^{Lys(UUU)}$ with swapped stems (14, rRWC025), ASL$^{Lys(UUU)}$ with reversed loop (15, rRWC036), ASL$^{Lys(UUU)}$ with swapped stems and reversed loop (16, rRWC037), ASL$^{Phe(GAA)}$ with swapped stems (17, rRWC038), chimeric ASL$^{Lys(UUU)}$ loop and ASL$^{Phe(GAA)}$ stems (18, rRWC038), and ASL$^{Phe(GAA)}$ with single deoxy-blocked cut site (19, rRWC034). (**E**) Associated rate constants (K) of curve fits from **Fig. S12** represented as mean +/- SD (n=4 technical replicates).

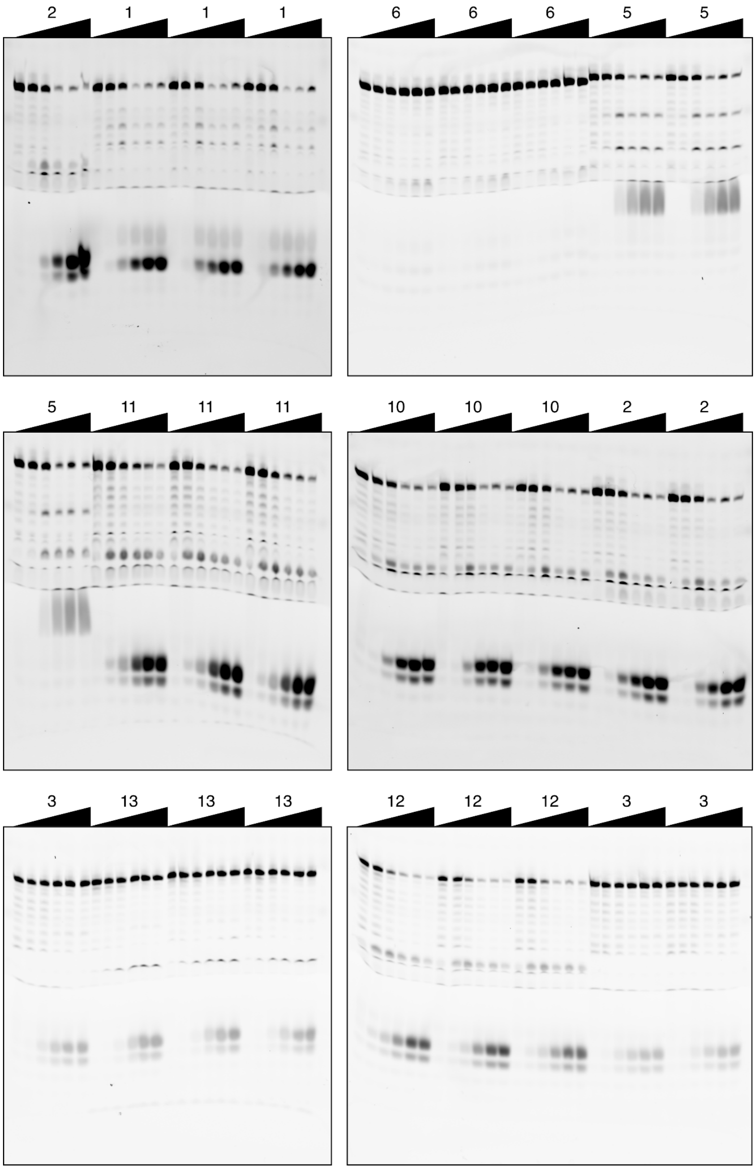
2 1 1 1
6 6 6 5 5
5 11 11 11
10 10 10 2 2
3 13 13 13
12 12 12 3 3

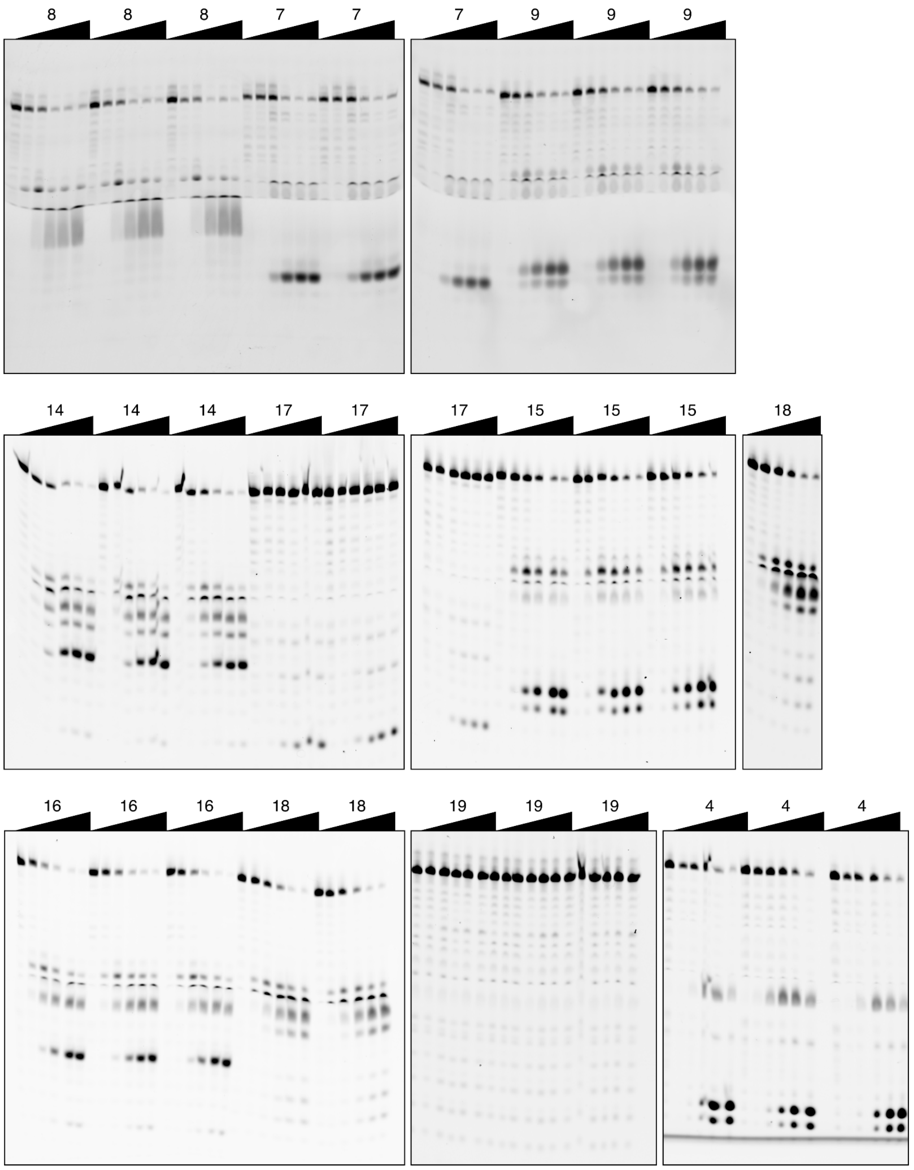


**Figure S11. Replicate gel images showing cleavage by LbuCas13a of 5′ 6-FAM RNA substrates.** Triangles above gels indicate a full reaction time course with a number to indicate the 5′ 6-FAM substrate as previously described (**Fig. S10**).

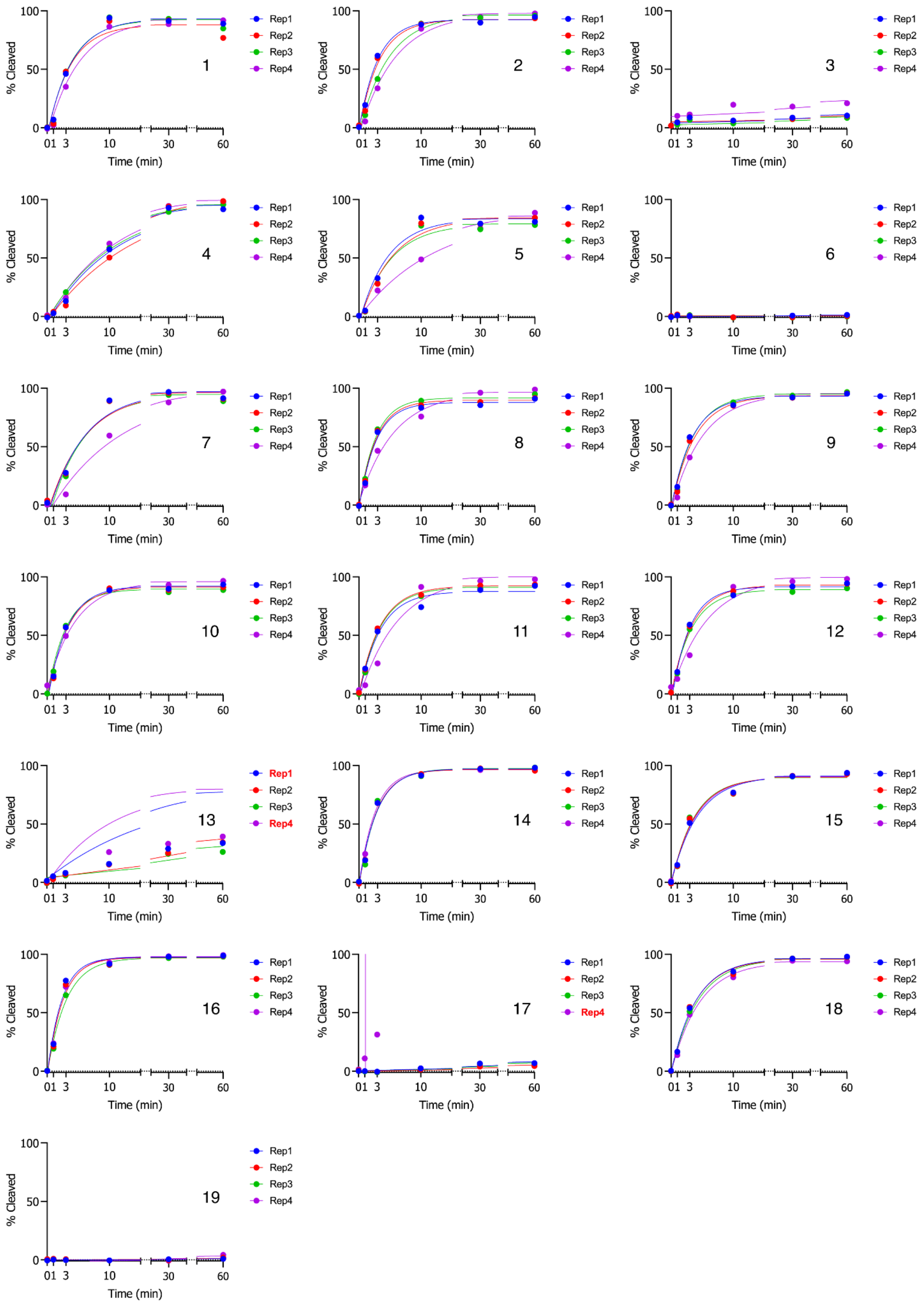


**Figure S12. Cleavage plots of fluorescent RNA substrates by LbuCas13a.** Plots of %Cleaved relative to full-length substrate and fitted non-linear regression curves per replicate (n=4 technical replicates) for all substrates listed in **Fig. S10E**. Replicates with bad fits are marked with red text and were excluded in further analysis.

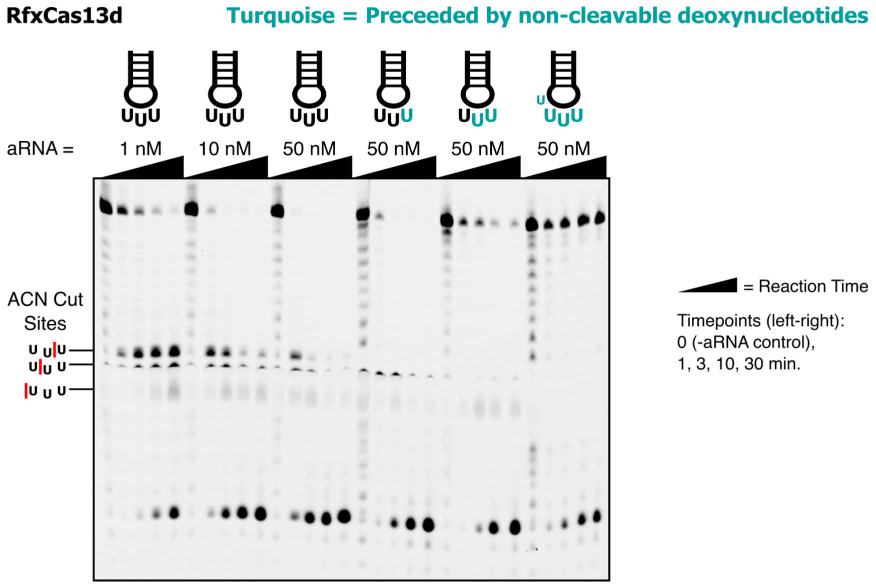


**Figure S13. RfxCas13d cleavage of tRNA$^{Lys}$ ACN Stem Loops.** Gel images showing cleavage by RfxCas13d of 5′ 6-FAM RNA substrates (n=1). Triangles above gels indicate a full reaction time course with the concentration of aRNA labelled above. Substrates are derived from tRNA ACN stem loops (ASL) from *Leptotrichia buccalis*. Starting from the left, substrates used are: ASL$^{Lys(UUU)}$ (Reactions 1-3, rRWC023), ASL$^{Lys(UUU)}$ with a single deoxy-blocked cut site (Reaction 4, rRWC032), or 2 (Reaction 5, rRWC035), or 4 (Reaction 6, rRWC033).

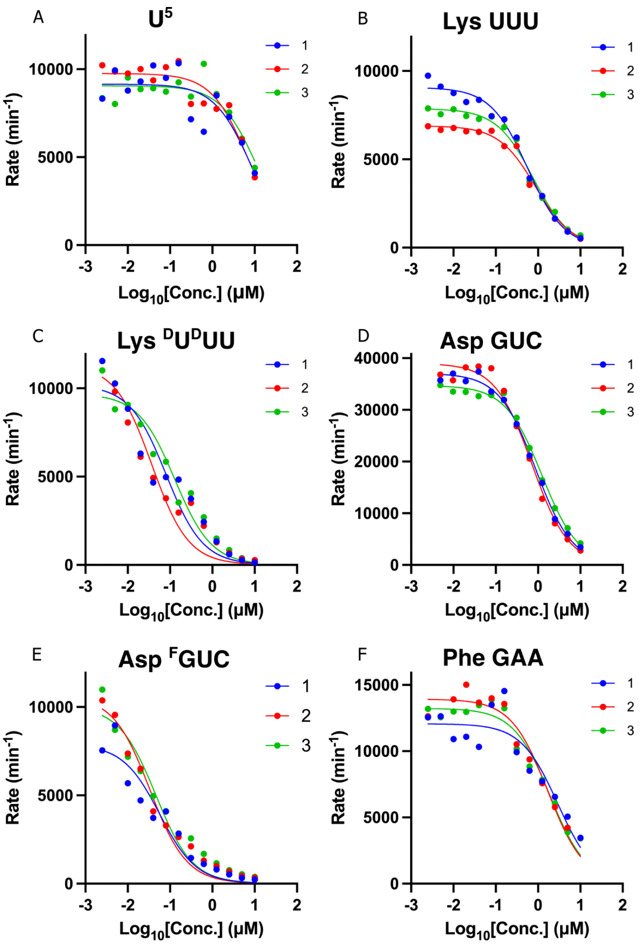


**Figure S14. Competition assay inhibition curves.** Linear slope values of data from **Fig. S15-20** plotted as a function of competitor concentration. Inhibitor Vs Response curves used to calculate IC50 values per replicate (n=3 technical replicates). Each competitor listed as in **Fig. 1E**.

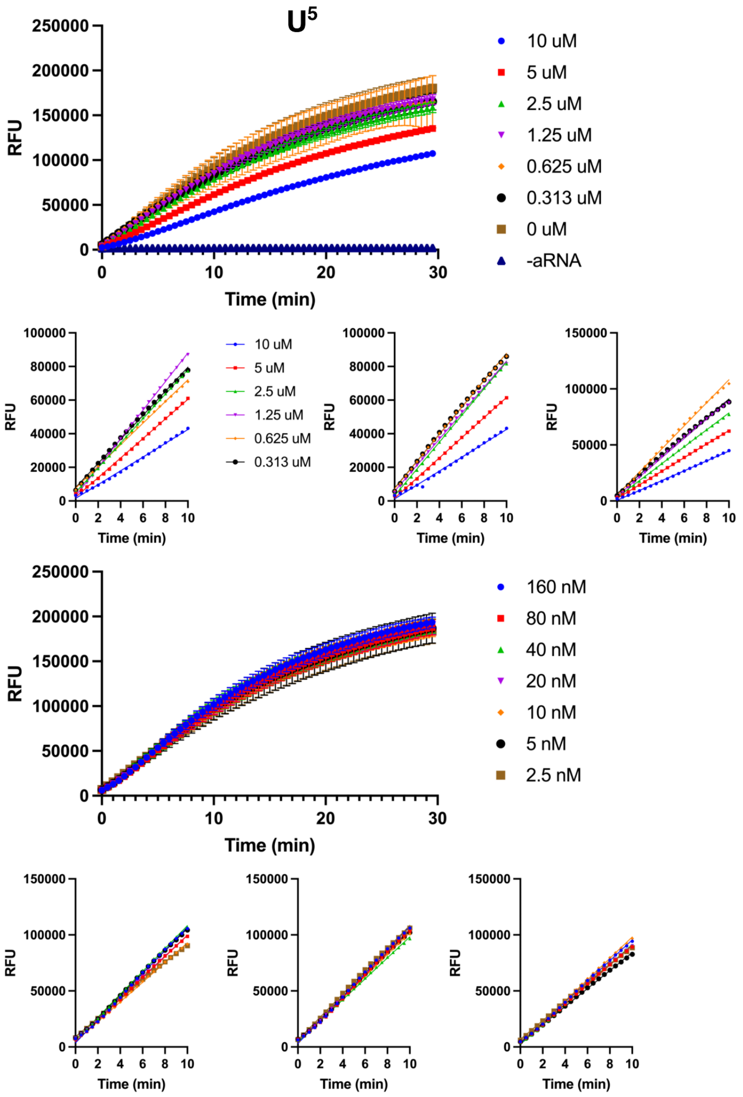


**Figure S15. Raw data of competition assay for $U^5$.** Cleavage of fluorescent reporter (rGJK123) measured in relative fluorescence units (RFU) against competitor rRWC013. Assay split into 2 dilution series. Linear range of assay determined to be <10 min post-initiation with data and linear curve fits shown in smaller graphs per replicate. The 0 µM and -aRNA curves are disregarded in further analysis.

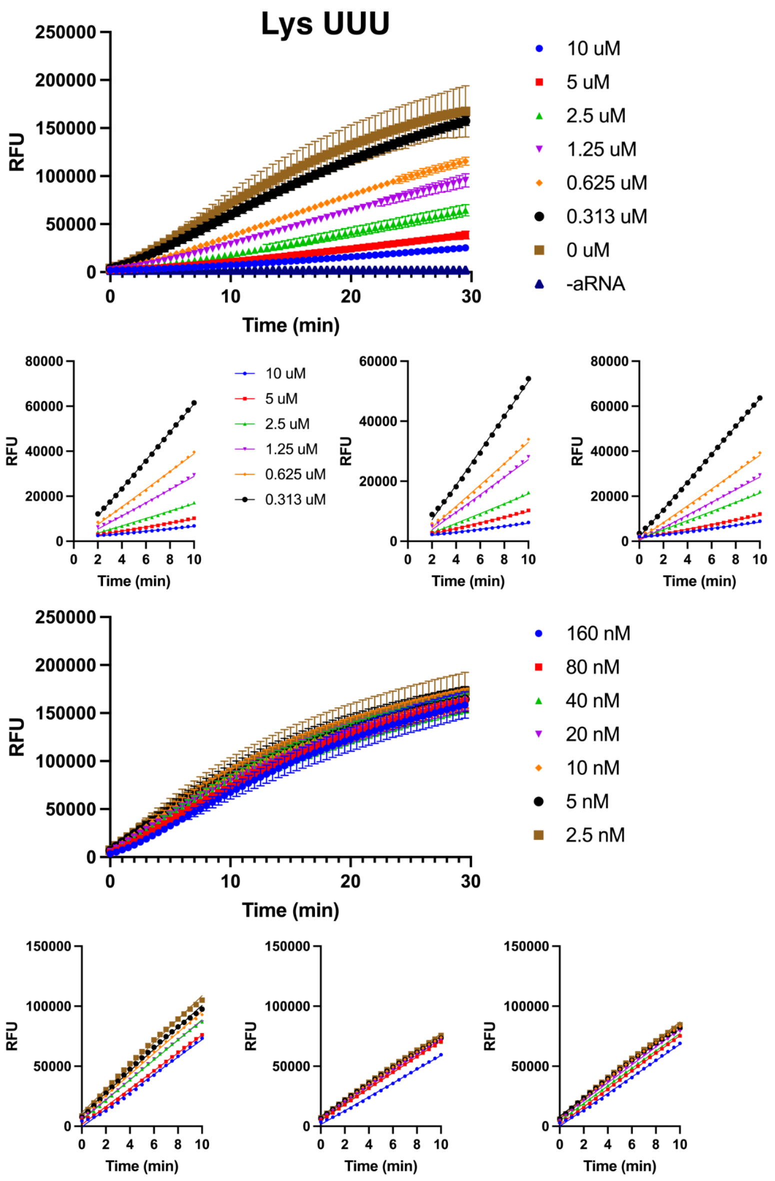


**Figure S16. Raw data of competition assay for ASL$^{Lys(UUU)}$.** Cleavage of fluorescent reporter (rGJK123) measured in relative fluorescence units (RFU) against competitor rRWC050. Assay split into 2 dilution series. Linear range of assay determined to be <10 min (or 2-10 min) post-initiation with data and linear curve fits shown in smaller graphs per replicate. The 0 µM and -aRNA curves are disregarded in further analysis.

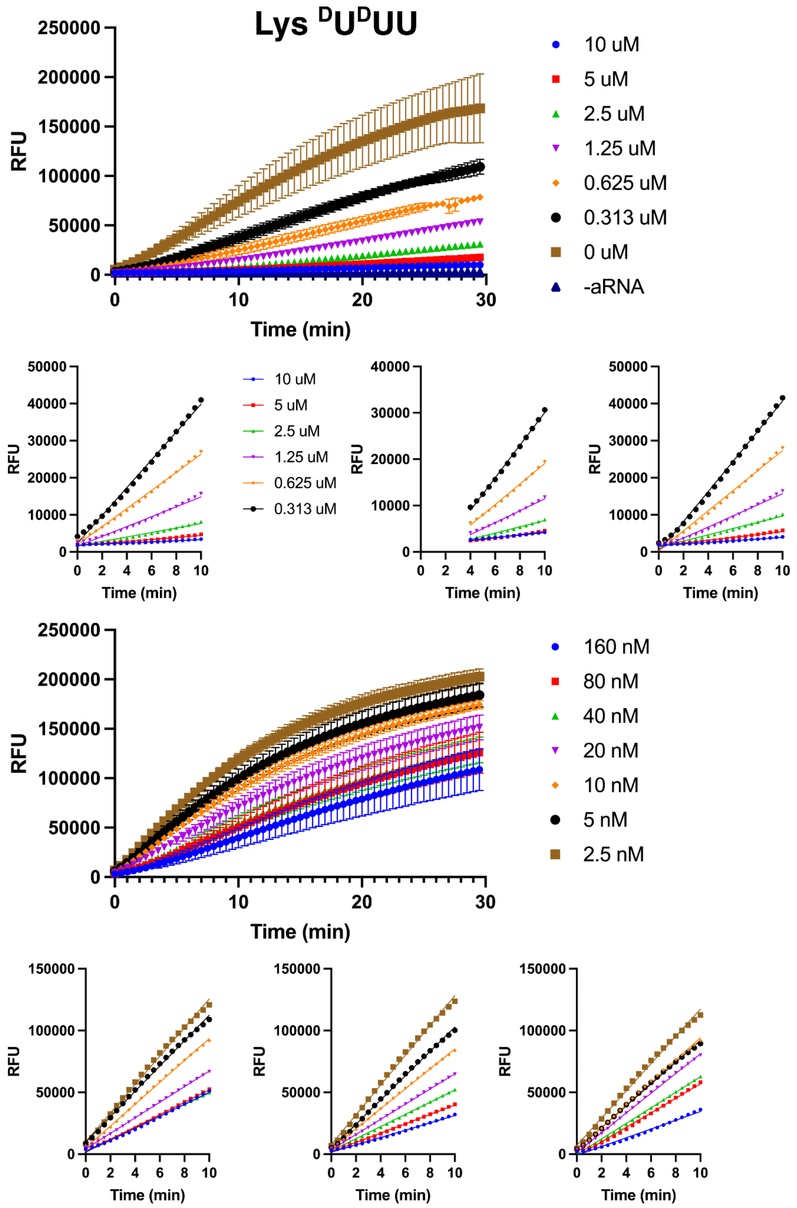


**Figure S17. Raw data of competition assay for $ASL^{Lys(dUdUU)}$.** Cleavage of fluorescent reporter (rGJK123) measured in relative fluorescence units (RFU) against competitor rRWC039. Assay split into 2 dilution series. Linear range of assay determined to be <10 min (or 4-10 min) post-initiation with data and linear curve fits shown in smaller graphs per replicate. The 0 µM and -aRNA curves are disregarded in further analysis.

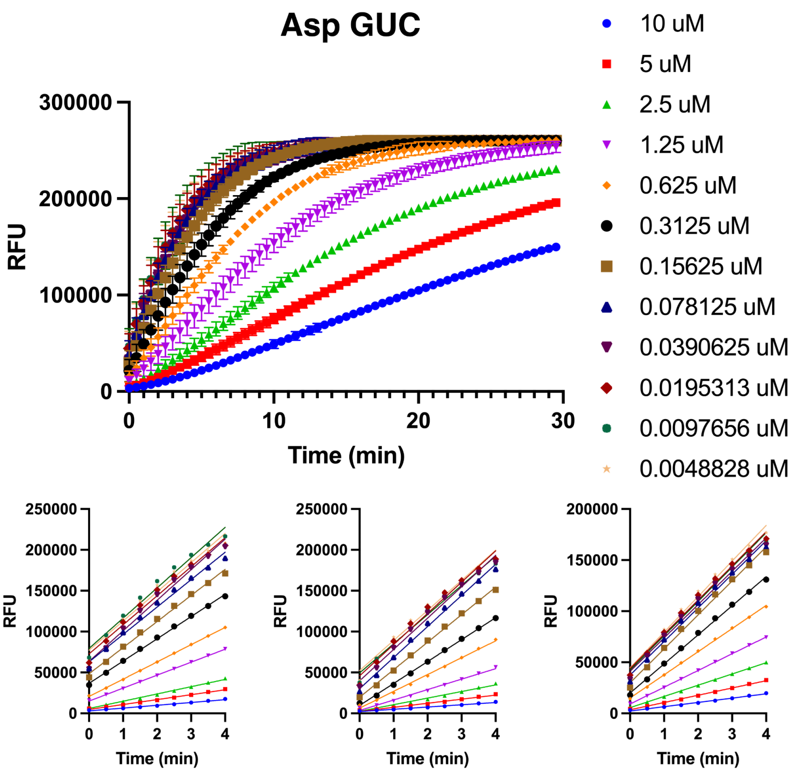


**Figure S18. Raw data of competition assay for $ASL^{Asp(GUC)}$.** Cleavage of fluorescent reporter (rGJK123) measured in relative fluorescence units (RFU) against competitor rRWC059. Linear range of assay determined to be <4 min post-initiation with data and linear curve fits shown in smaller graphs per replicate. This competition assay used 8 pM aRNA.

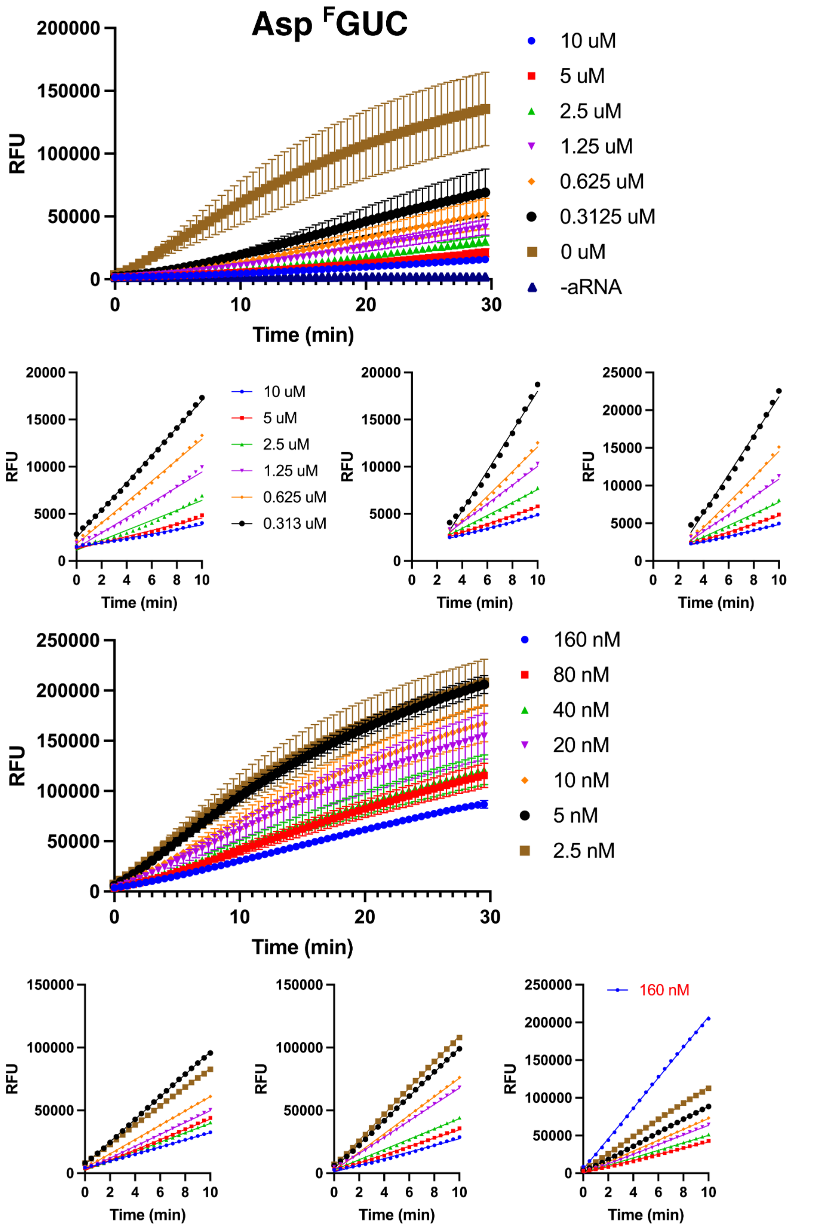


**Figure S19. Raw data of competition assay for ASL$^{Asp(fGUC)}$.** Cleavage of fluorescent reporter (rGJK123) measured in relative fluorescence units (RFU) against competitor rRWC060. Assay split into 2 dilution series. Linear range of assay determined to be <10 min (or 3-10 min) post-initiation with data and linear curve fits shown in smaller graphs per replicate. Outlier replicate conditions are marked with red text and were excluded from further analysis. The 0 µM and -aRNA curves are disregarded in further analysis.

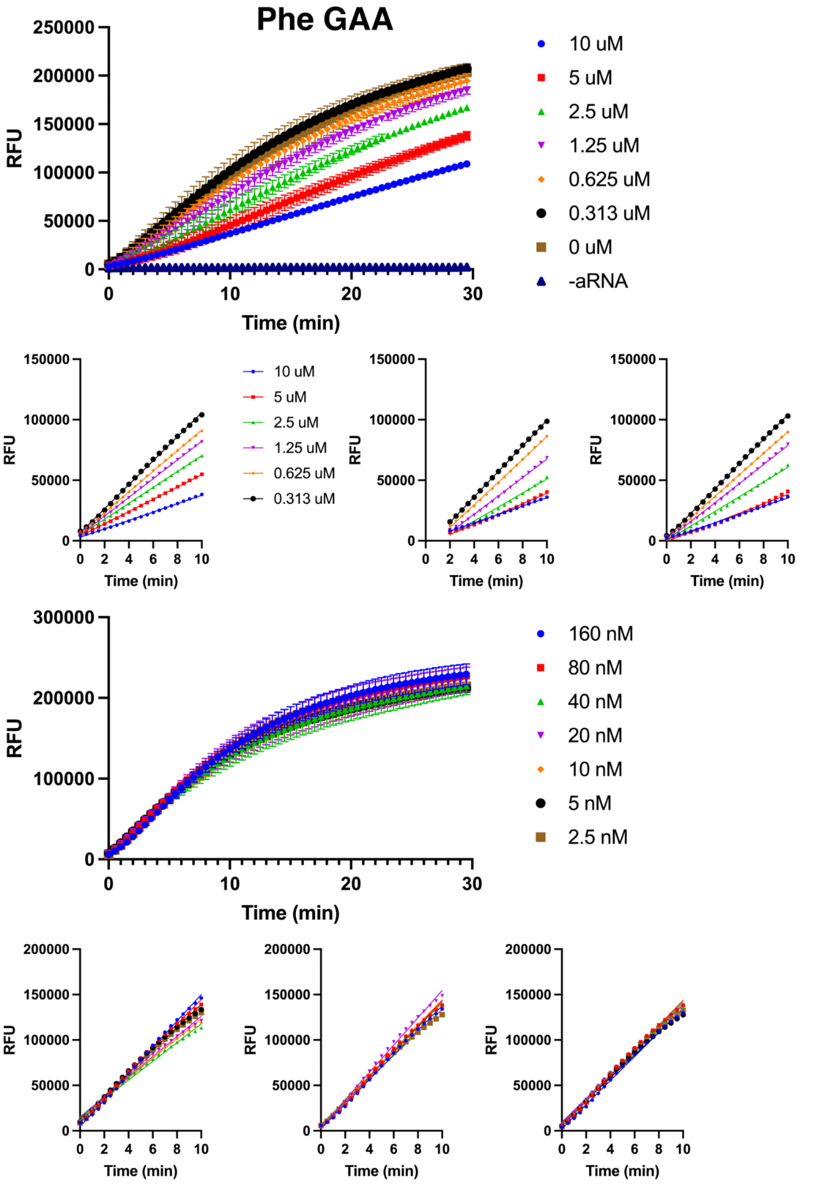


**Figure S20. Raw data of competition assay for $\text{ASL}^{\text{Phe(GAA)}}$.** Cleavage of fluorescent reporter (rGJK123) measured in relative fluorescence units (RFU) against competitor rRWC051. Assay split into 2 dilution series. Linear range of assay determined to be <10 min (or 2-10 min) post-initiation with data and linear curve fits shown in smaller graphs per replicate. The 0 µM and -aRNA curves are disregarded in further analysis.

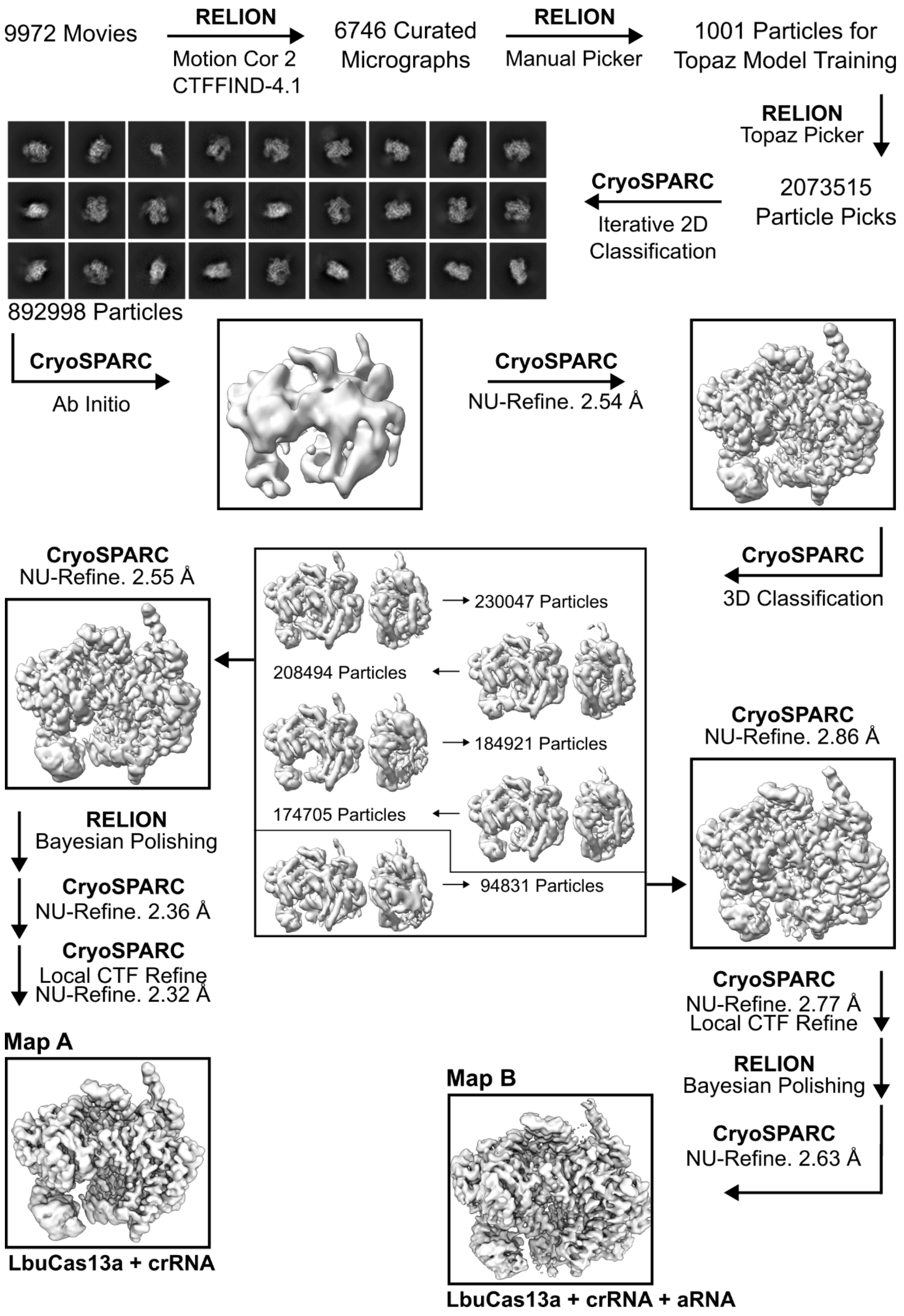


**Figure S21. CryoEM data analysis pipeline of LbuCas13a complex without substrate.** Processed using cryoSPARC (4.5.3) and RELION (4.0.0).

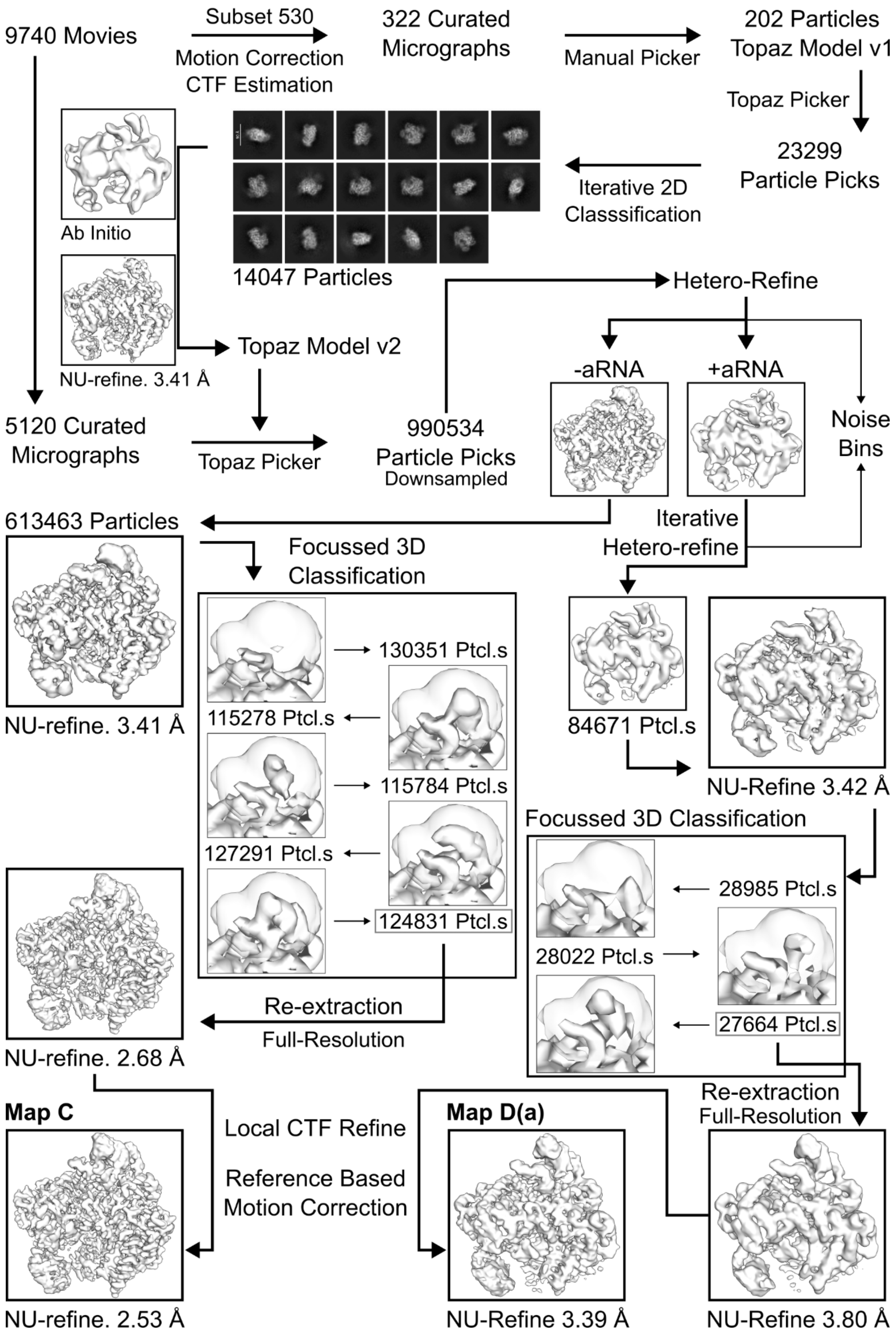


**Figure S22. CryoEM data analysis pipeline of LbuCas13a complex with ASL$^{Lys(dUdUU)}$ (Dataset 1).** Processed using cryoSPARC (4.5.3).

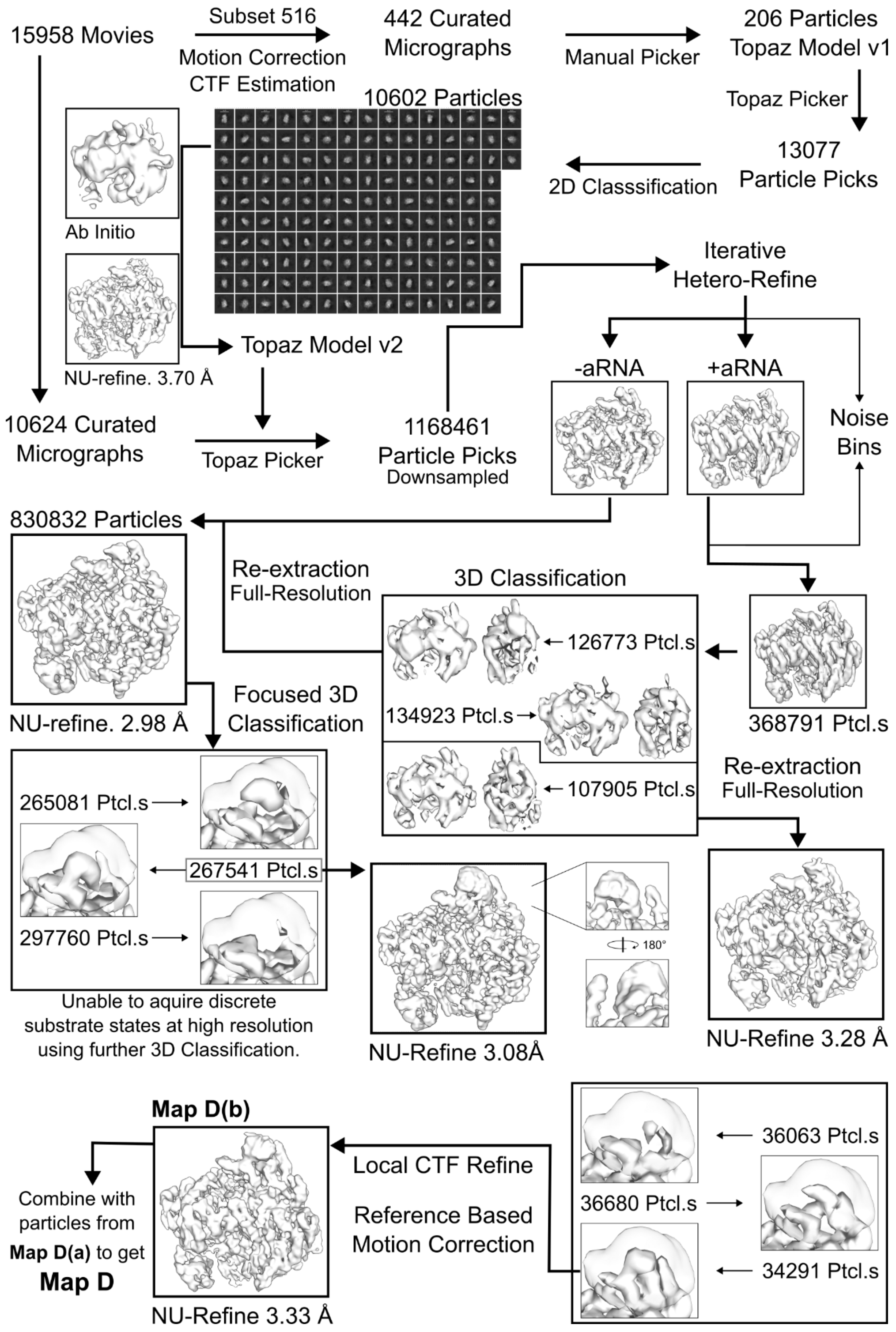


**Figure S23. CryoEM data analysis pipeline of LbuCas13a complex with ASL$^{Lys(dUdUU)}$ (Dataset 2).** Processed using cryoSPARC (4.5.3).

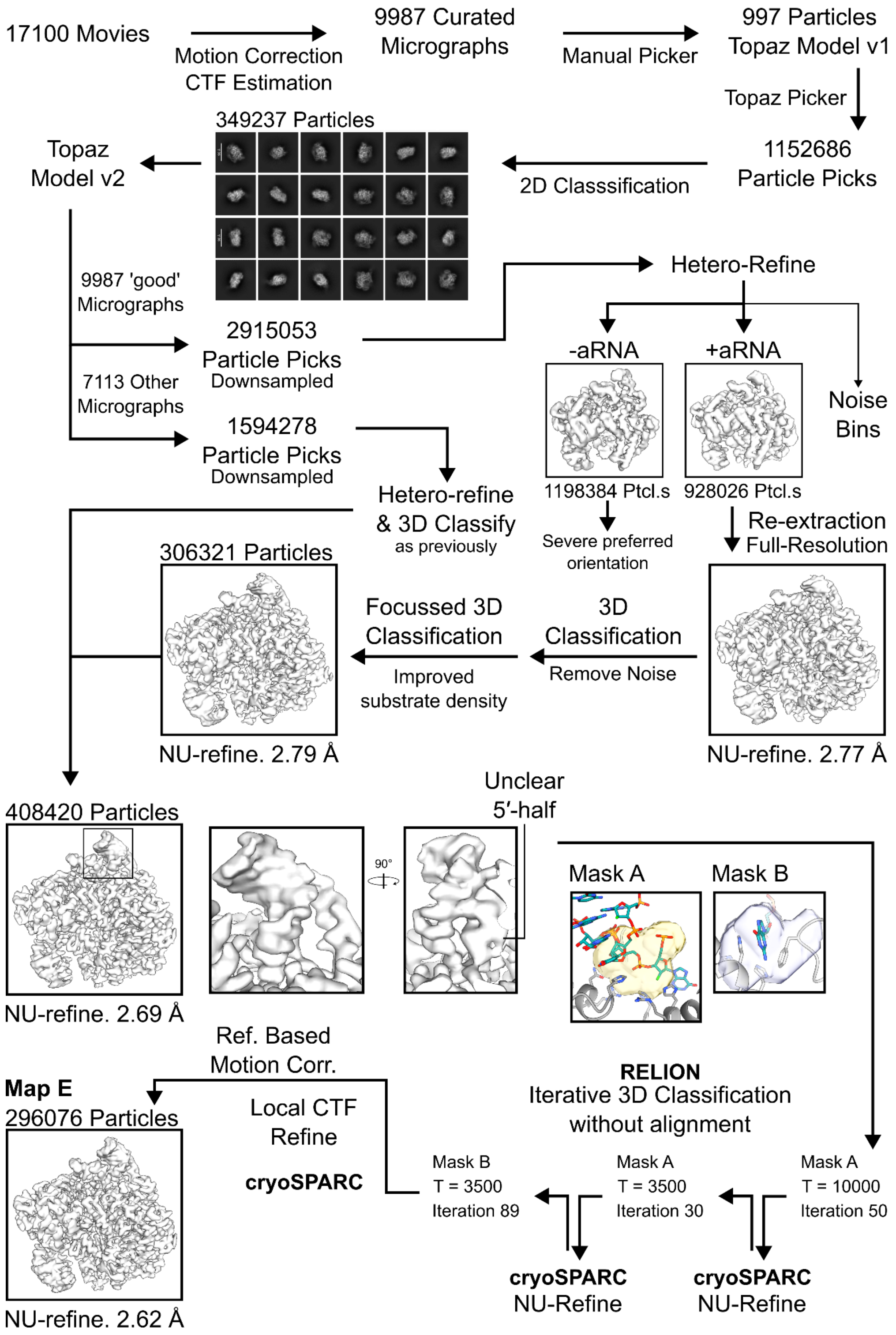


**Figure S24. CryoEM data analysis pipeline of LbuCas13a complex with ASL$^{Asp(fGUC)}$.** Processed using cryoSPARC (4.5.3) and RELION (5.0.0) for 3D classification without alignment.

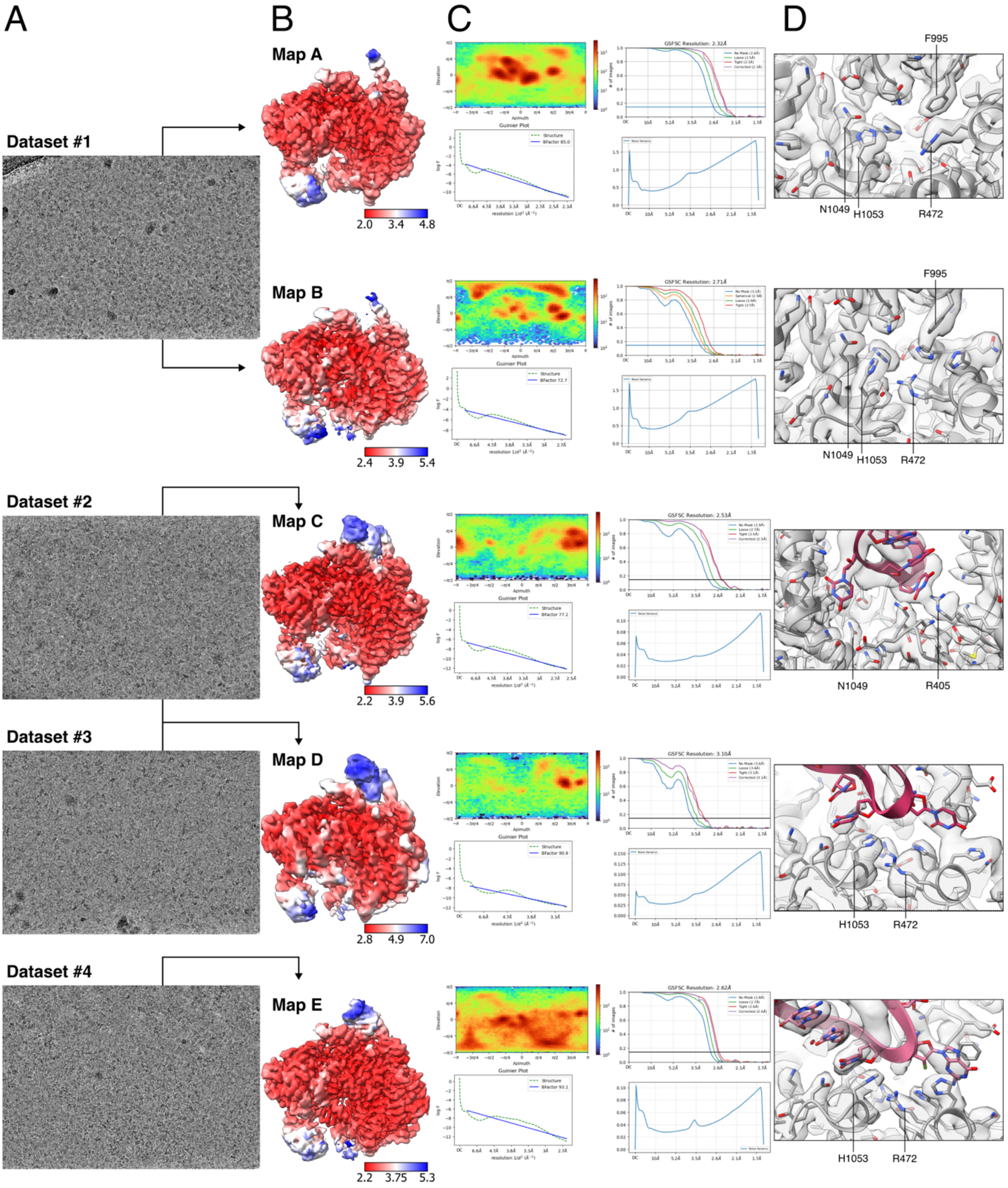


**Figure S25. Summary statistics and model fits of cryoEM maps.** Maps and models sorted as outlined in Table S1. (A) Representative micrographs of datasets to reconstruct LbuCas13a states. (B) Final maps coloured by local resolution. (C) Summary validation plots after refinement including orientation plot, FSC plot, noise model plot, and Guinier plot. (D) Representative map-model fit images focused at key HEPN regions.

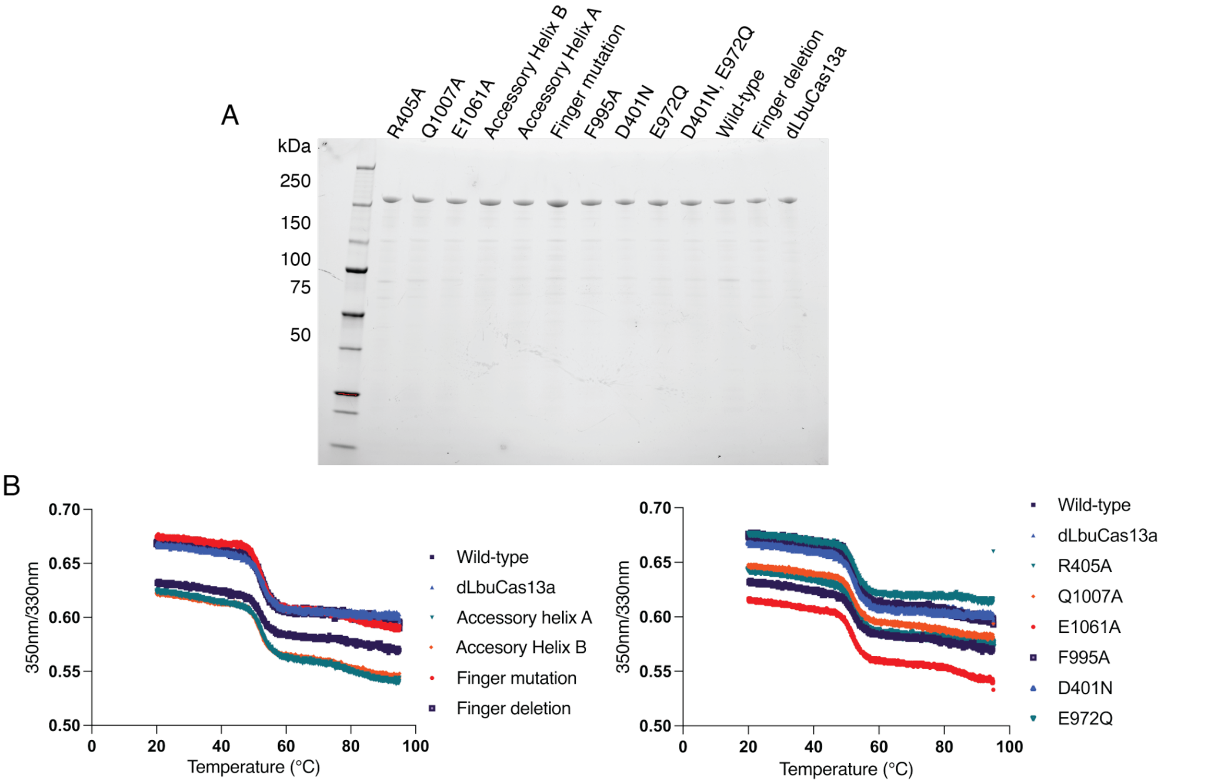


**Figure S26. Purification and folding of LbuCas13a mutants.** (**A**) SDS-PAGE showing all purified mutants. Single point mutants are as indicated, Accessory Helix B (K1036A, K1039A, Q1040A, K1042A), Accessory Helix A (K1004A, K1011A, K1014A, K1017A), Finger mutation (K409A, N414A, K415A, K419A), Finger deletion (Δ409-421) and dLbuCas13a (H477N, H1053N). (**B**) nanoDSF thermal unfolding of mutants in A) (n=1). Left shows nanoDSF for accessory elements, right shows nanoDSF for point mutants. No decrease in structural stability was observed for any mutants.

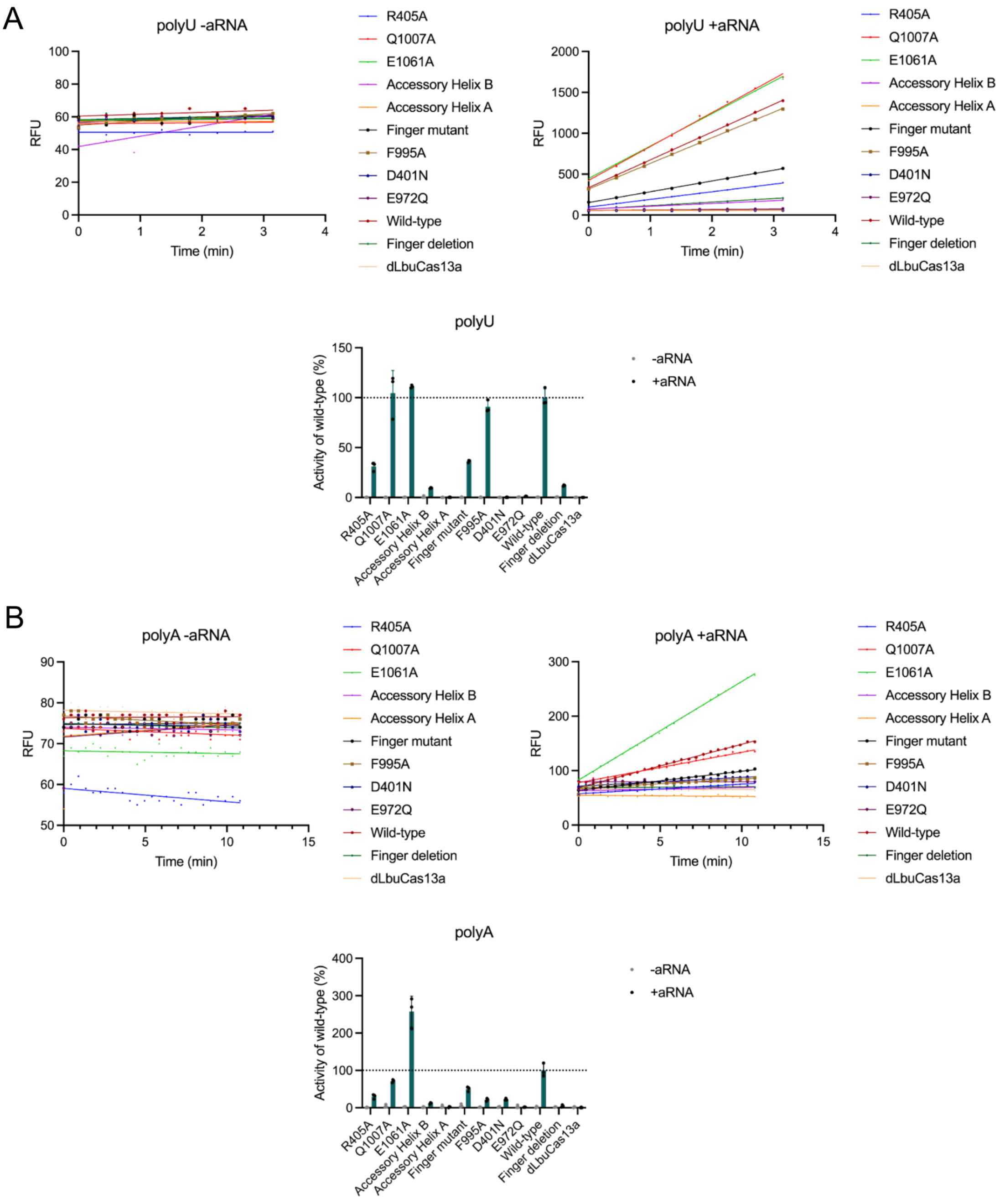
A
polyU -aRNA
RFU
Time (min)
polyU +aRNA
R405A
Q1007A
E1061A
Accessory Helix B
Accessory Helix A
Finger mutant
F995A
D401N
E972Q
Wild-type
Finger deletion
dLbuCas13a
polyU
Activity of wild-type (%)
-aRNA
+aRNA
B
polyA -aRNA
polyA +aRNA
polyA

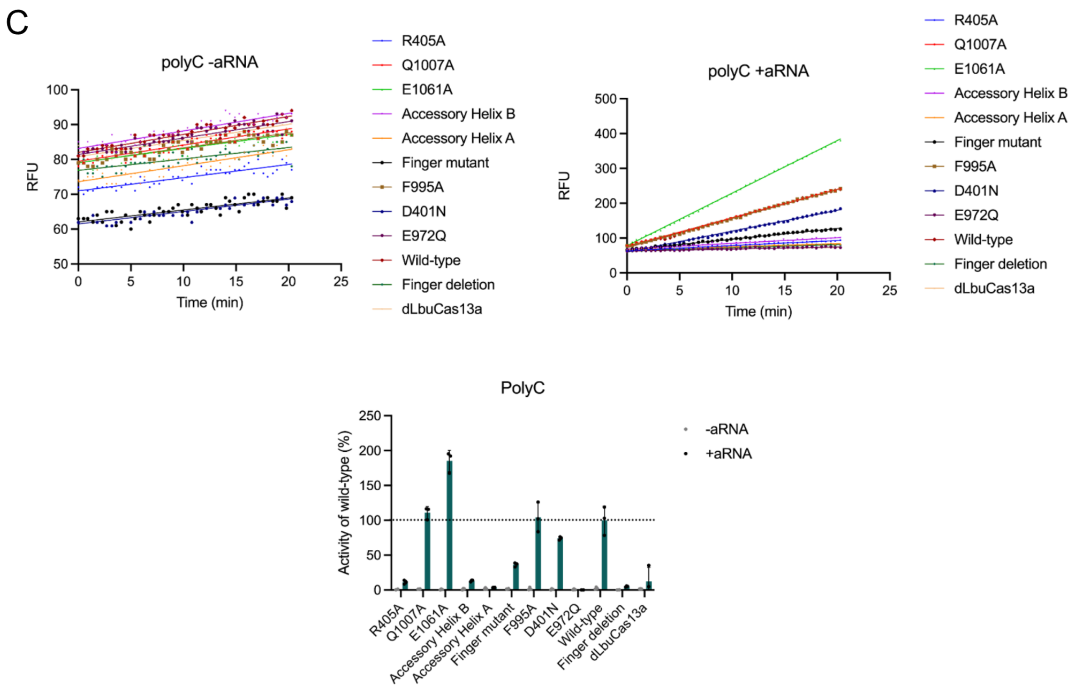
C
polyC -aRNA
RFU
Time (min)
polyC +aRNA
R405A
Q1007A
E1061A
Accessory Helix B
Accessory Helix A
Finger mutant
F995A
D401N
E972Q
Wild-type
Finger deletion
dLbuCas13a
PolyC
Activity of wild-type (%)
-aRNA
+aRNA

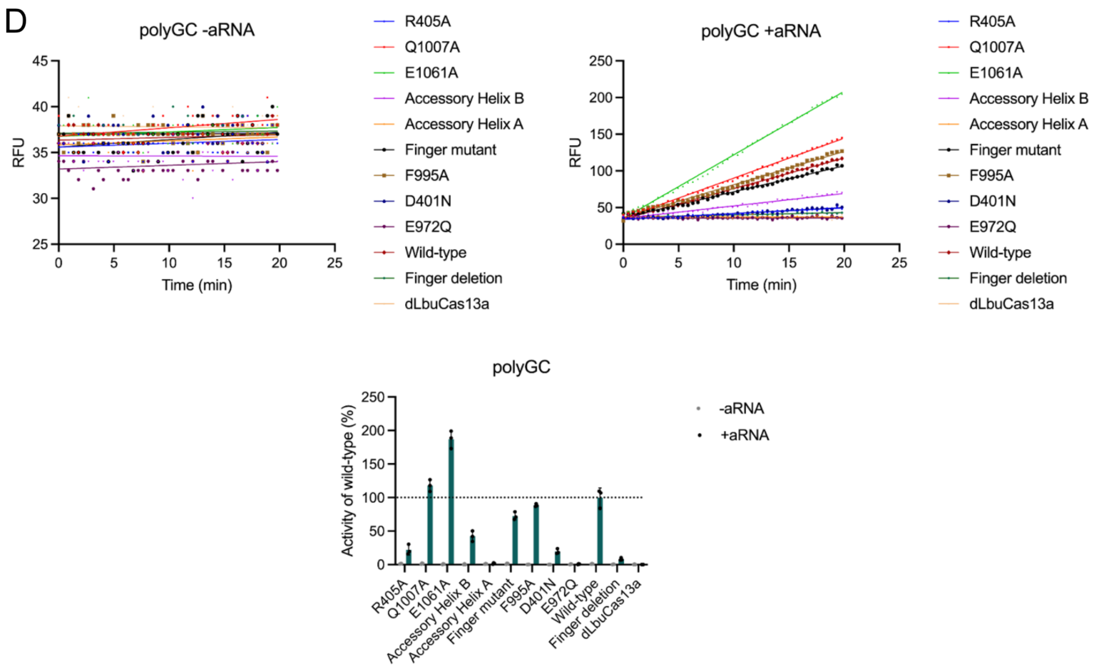
D
polyGC -aRNA
RFU
Time (min)
polyGC +aRNA
R405A
Q1007A
E1061A
Accessory Helix B
Accessory Helix A
Finger mutant
F995A
D401N
E972Q
Wild-type
Finger deletion
dLbuCas13a
polyGC
Activity of wild-type (%)
-aRNA
+aRNA

E

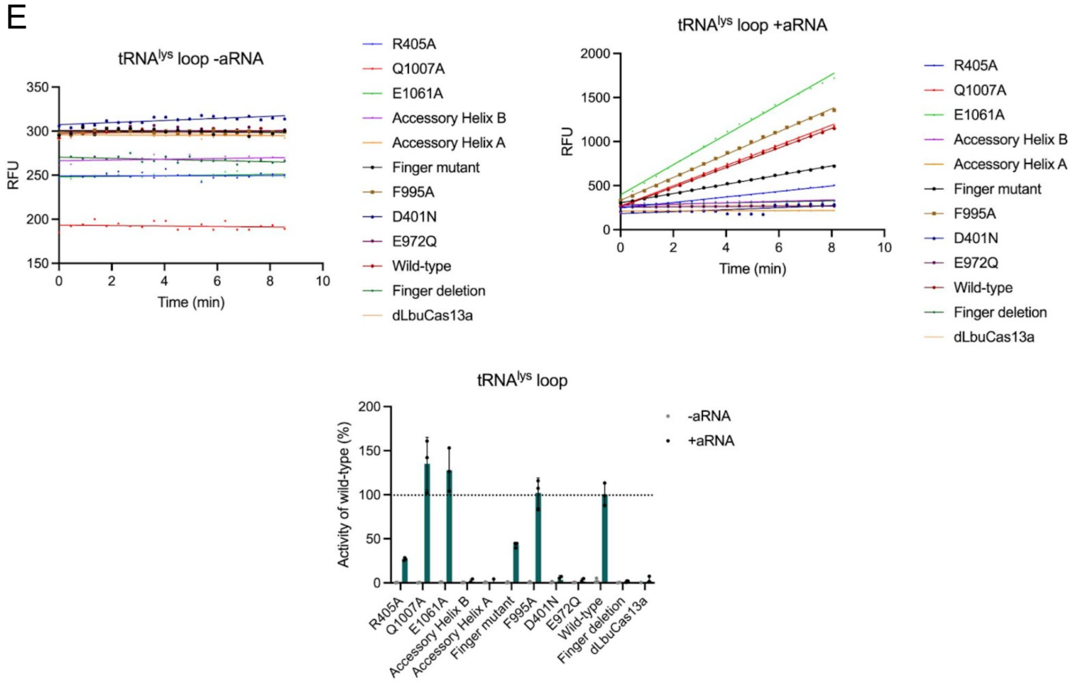


F

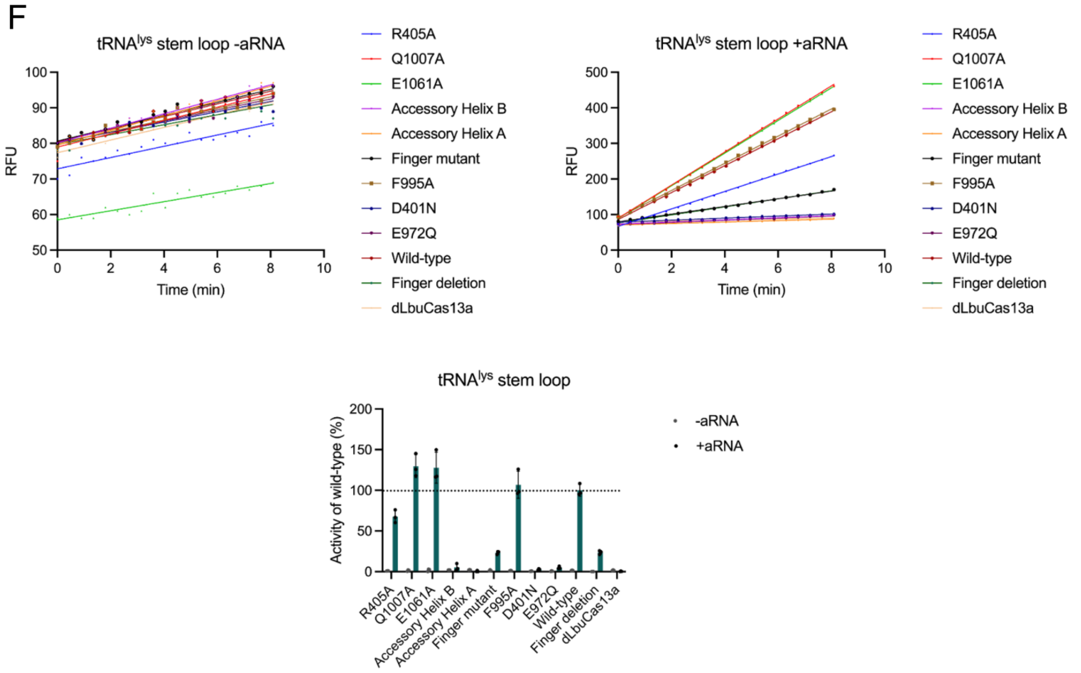


**Figure S27. Raw data for activity testing of LbuCas13a mutants.** Summary of data is presented in Fig. 2, relating to mutants from **Fig. S26**. Single point mutants are as indicated, Accessory Helix B (K1036A, K1039A, Q1040A, K1042A), Accessory Helix A (K1004A, K1011A, K1014A, K1017A), Finger mutation (K409A, N414A, K415A, K419A), Finger deletion (Δ409-421) and dLbuCas13a (H477N, H1053N). Substrates used for the assays are (**A**) $U^5$ (rGJK123), (**B**) $A^5$ (rBKH017), (**C**) $C^5$ (rBKH005), (**D**) CGCGC (rGJK400) (**E**) $tRNA^{Lys}$ loop (rBKH002) and (**F**) $tRNA^{Lys}$ structured stem loop (rRWC041). Each sample has n=1 shown, but n=3 was conducted for the assays with and without activator RNA (+aRNA and -aRNA, respectively). For each substrate, -aRNA is top left, +aRNA top right. Bottom shows activity normalised to wildtype. The -aRNA data is purple bars with individual points in grey, and +aRNA data are shown as green bars with individual points in black. For all, n=3 technical replicates with the mean shown and error bars indicating SD.

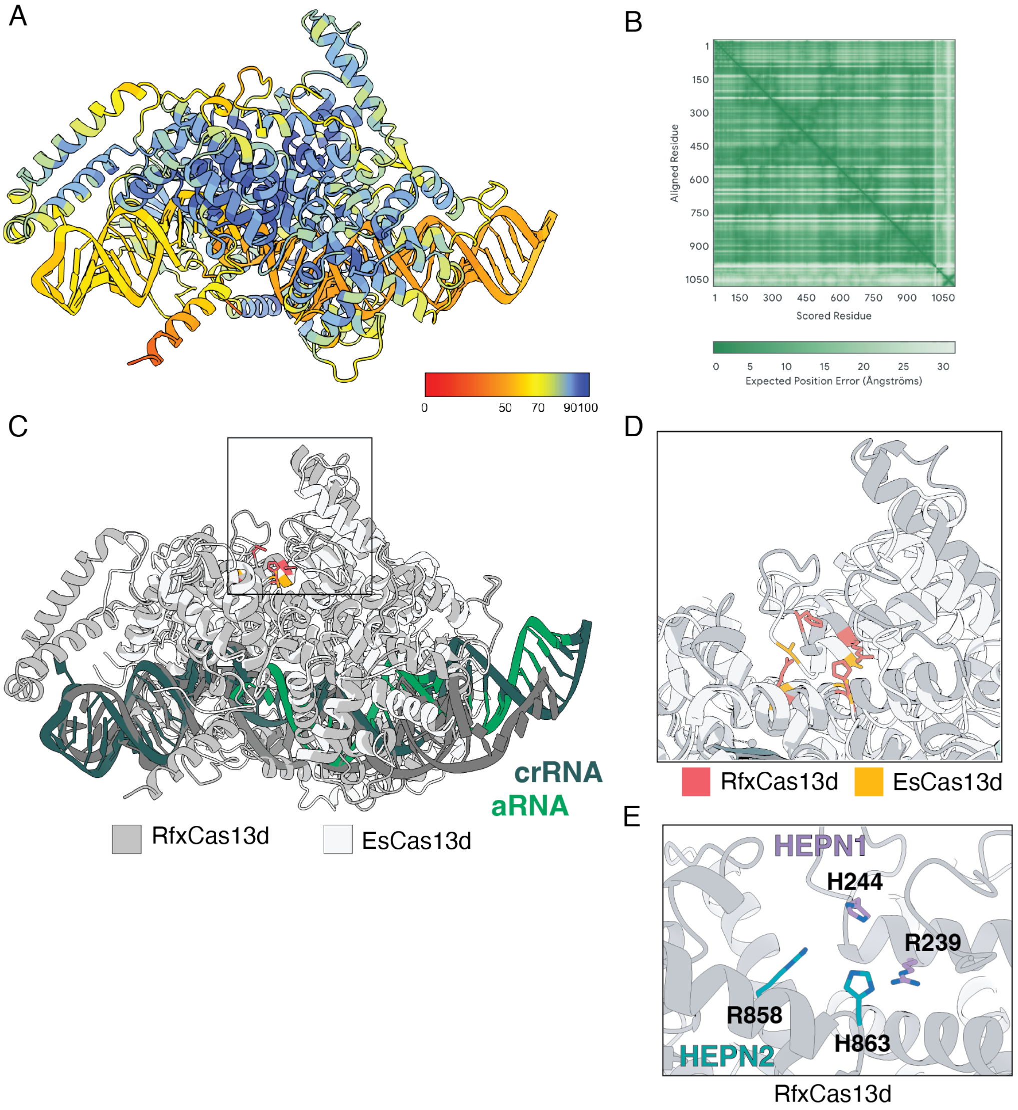


**Figure S28. AlphaFold3 RfxCas13d model.** Five models were produced and assessed for convergence. Compared to the top model, all other models had an RMSD range of 0.71 - 2.3 Å. (**A**) top model of ternary RfxCas13d (RfxCas13d:crRNA:aRNA) coloured by model confidence. (**B**) PAE plot for the top model. (**C**) Flat cartoon of RfxCas13d AlphaFold3 model aligned to EsCas13d structure (PDB 6E9F) (*28*): global RMSD 7.1Å. RfxCas13d mid grey, crRNA dark green and aRNA light green. EsCas13d light grey, crRNA and aRNA dark grey. (**D**) Comparison of the RfxCas13d and EsCas13d active site. Key HEPN residues are shown in red for RfxCas13d (R238, H244, R858, H863), and yellow for EsCas13d (these are mutated to Ala in the structure: R295A, H300A, R849A, H854A). (**E**) Zoom into the active site of ternary RfxCas13d shown as a flat cartoon in grey (region shown as a box on panel B): catalytic residues are shown as sticks and coloured by heteroelement. HEPN1 coloured purple and HEPN2 coloured green. Residue numbers are as indicated.

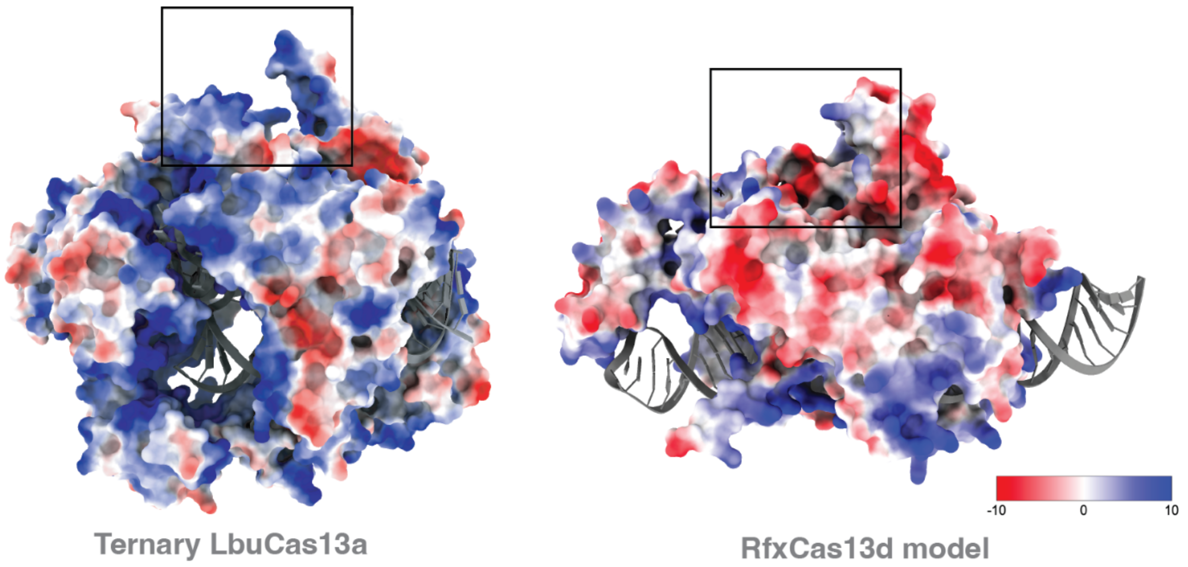


**Figure S29. Surface charge comparison for LbuCas13a cryoEM structure and RfxCas13d AlphaFold3 prediction.** Both structures are ternary state (protein:crRNA:aRNA). Left) LbuCas13a and right) RfxCas13d AlphaFold3 prediction. Boxes for both show the substrate binding pocket. Blue denotes positive charge and red denotes negative charge.

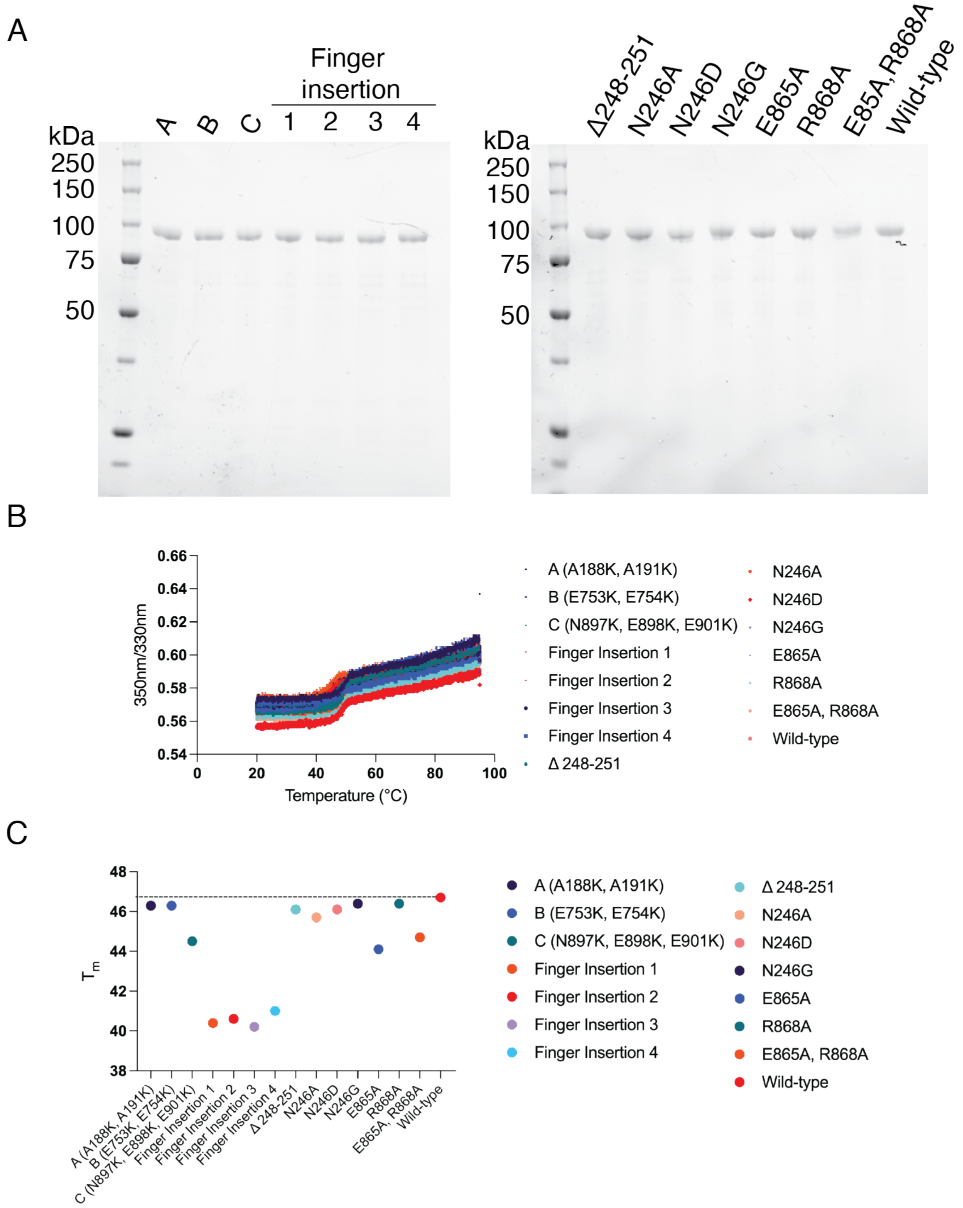


**Figure S30. Purification and folding of RfxCas13d mutants.** (**A**) SDS-PAGE showing all purified mutants. Left shows charged mutants A (A188K, A191K), B (E753K, E754K) and C (N897K, E898K, E901K) as well as finger insertion mutants between residues 185 - 185 (1), 188 - 189 (2), 189 - 190 (3) and 192 - 193 (4). Right shows specificity mutants where mutants are as indicated, (**B**) nanoDSF thermal unfolding of mutants in A) (n=1). (**C**) plotted Tm values for all mutants that show a decrease in stability for all finger insertion mutants. $T_m$ is shown on the Y axis, and mutations on the X axis.

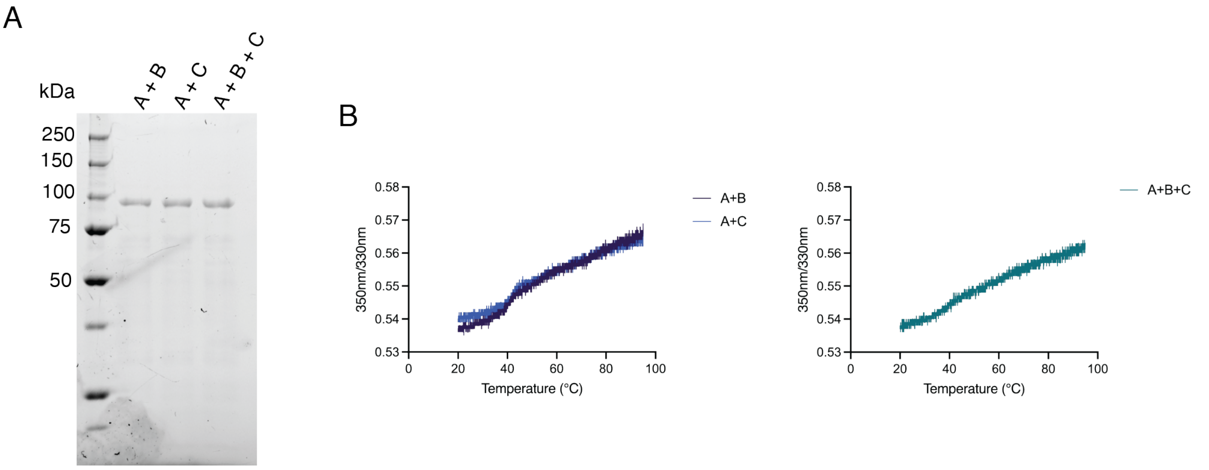


**Figure S31. Purification and folding of RfxCas13d stacked charge substitution mutants.** (**A**) SDS-PAGE showing all purified RfxCas13d stacked charge mutants. A + B includes mutations (A188K, A191K, E753K, E754K), A + C includes (A188K, A191K, N897K, E898K, E901K) and A + B + C (A188K, A191K,E753K, E754K, N897K, E898K, E901K). (**B**) thermal melt nanoDSF of mutants in panel A (n=1). Mutants are as panel A. Given the wildtype $T_m$ value of 47°C, all mutants had an observed change in structural stability.

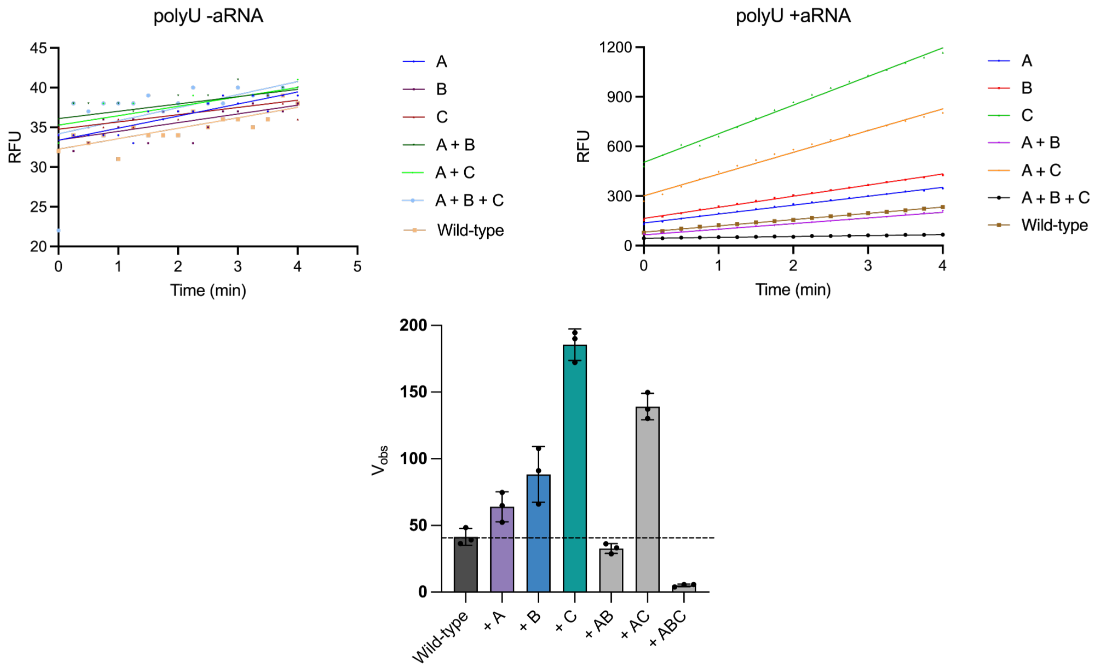


**Figure S32. Raw activity data for RfxCas13d charged mutants.** For all panels: A (A188K, A191K), B (E753K, E754K), C (N897K, E898K, E901K), A + B includes mutations (A188K, A191K, E753K, E754K), A + C includes (A188K, A191K, N897K, E898K, E901K) and A + B + C ( A188K, A191K,E753K, E754K, N897K). These data relate to mutants from **Fig. S30-31**. Substrate used for the assays is $U^5$ (rGJK123). Each mutant has n=1 shown as an example, but n=3 was conducted for the assays with and without activator RNA (+aRNA and -aRNA, respectively). For each substrate, -aRNA is shown on the top left, +aRNA top right and bottom shows the observed velocity for each mutant and wildtype. For all, n=3 technical replicates with the mean shown and error bars indicating SD of the mean.

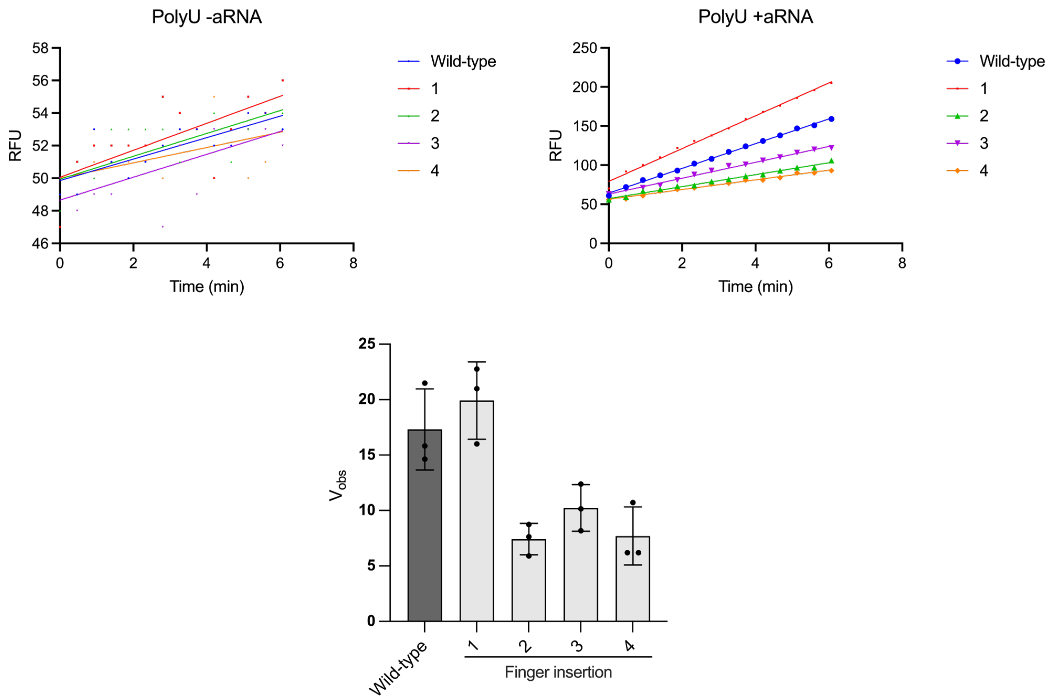


**Figure S33. Raw activity data for RfxCas13d finger insertion mutants.** For all panels: finger insertion mutants between residues 185 - 185 (1), 188 - 189 (2), 189 - 190 (3) and 192 - 193 (4). These data relate to mutants from **Fig. S30**. Substrate used for the assays is $U^5$ (rGJK123). Each mutant has n=1 shown as an example, but n=3 was conducted for the assays with and without activator RNA (+aRNA and -aRNA, respectively). For each substrate, -aRNA is shown on the top left, +aRNA top right and bottom shows the observed velocity for each mutant and wildtype. For all, n=3 technical replicates with the mean shown and error bars indicating SD of the mean.

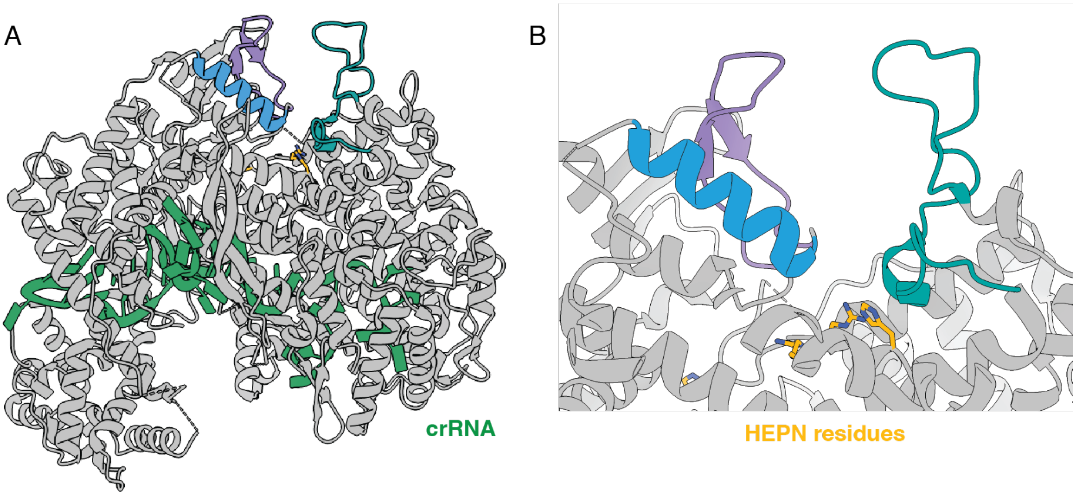


**Figure S34. Accessory elements for LbaCas13a.** Shown is the crystal structure of binary LbaCas13a (PDB ID: 5W1I (*78*)) as a flat cartoon. (**A**) depicts the global structure of LbaCas13a (grey) with crRNA bound (green). Proposed accessory elements are coloured purple, blue and cyan from left to right. Active site HEPN residues are shown as sticks, and coloured yellow and by heteroelement. (**B**) into LbaCas13a active site as shown in A.

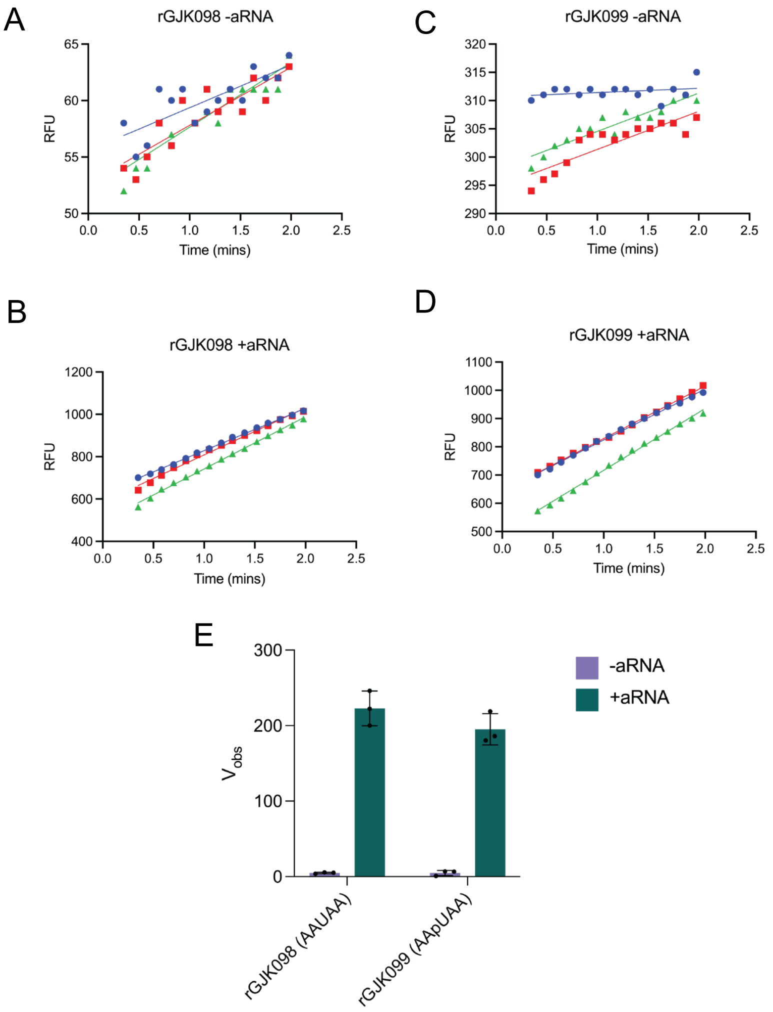


**Figure S35. Comparison of LbuCas13a activity against a uridine and pseudo-uridine containing substrates.** For panels A, B, C and D each colour (blue, green and red) indicates a replicate. Panels (**A**) and (**B**) the substrate used contains a standard U (rGJK098: AAUAA) and for panels (**C**) and (**D**) the substrate used contains a pseudo-U (rGJK099: AApUAA). Panels A) and C) are minus activator RNA (-aRNA, binary complex) and B) and D) are plus activator (+aRNA, ternary complex). Panel (**E**) shows the observed velocity of the linear phase of the reaction in the top panels (A, B, C and D). The -aRNA data is shown as purple bars and +aRNA data are shown as green bars. For both datasets, individual points in black with the bar showing the mean (n=3 technical replicate) and error bars indicating SD of the mean.

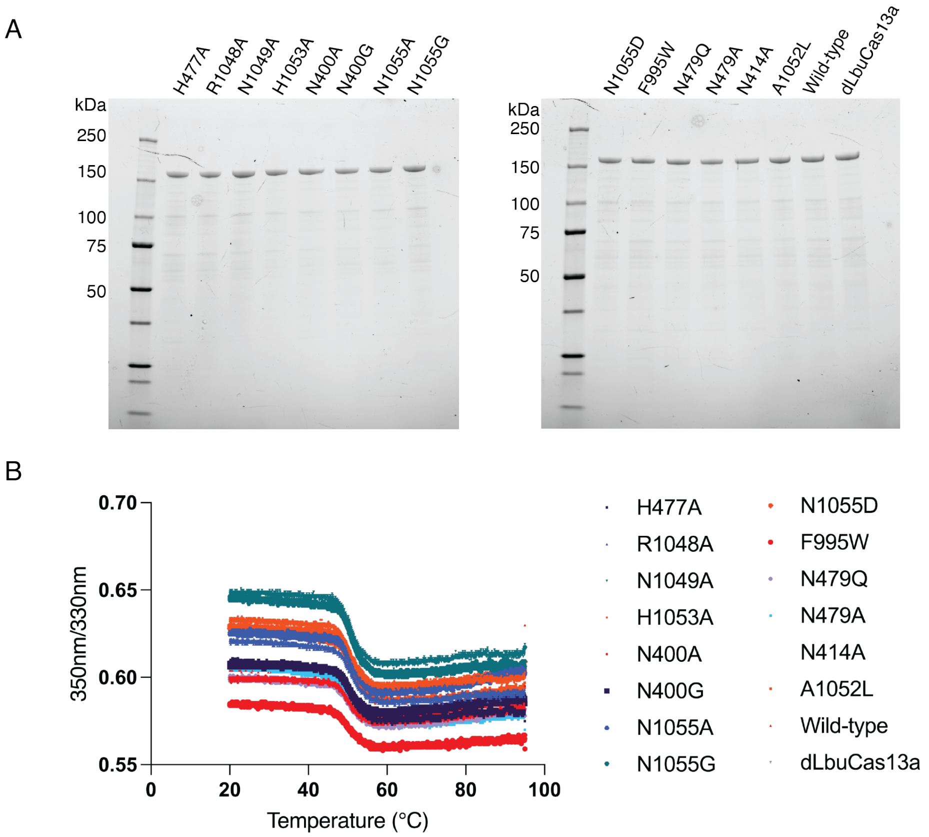


**Figure S36. Purification and folding of LbuCas13a specificity and HEPN mutants. (A)** SDS-PAGE showing all purified mutants. Single point mutants are as indicated, while dLbuCas13a denotes H477N, H1053N. (**B**) nanoDSF thermal unfolding of mutants in A) (n=1). No decrease in structural stability was observed for any mutants.

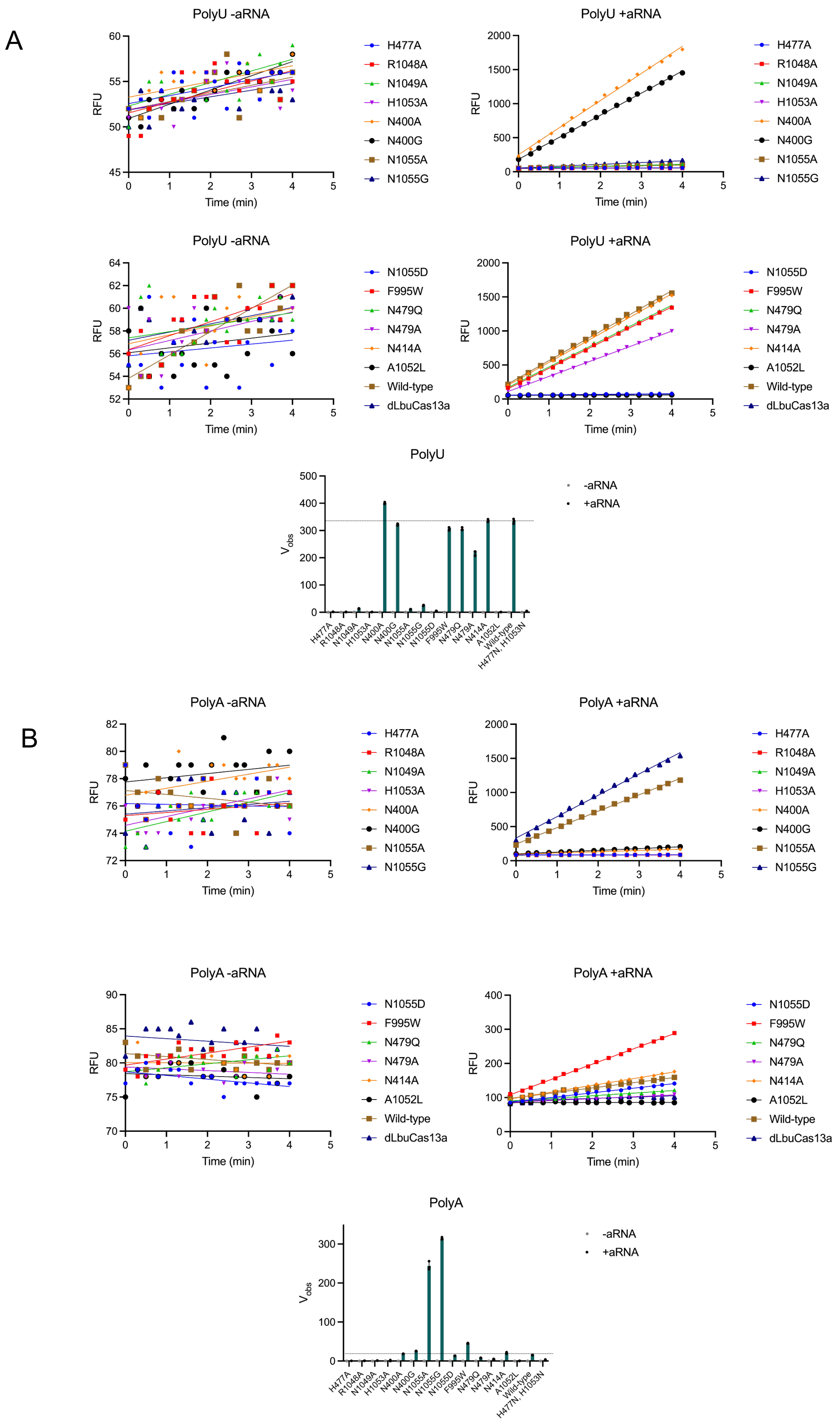
A
PolyU -aRNA
RFU
Time (min)
H477A
R1048A
N1049A
H1053A
N400A
N400G
N1055A
N1055G
PolyU +aRNA
RFU
Time (min)
H477A
R1048A
N1049A
H1053A
N400A
N400G
N1055A
N1055G
PolyU -aRNA
RFU
Time (min)
N1055D
F995W
N479Q
N479A
N414A
A1052L
Wild-type
dLbuCas13a
PolyU +aRNA
RFU
Time (min)
N1055D
F995W
N479Q
N479A
N414A
A1052L
Wild-type
dLbuCas13a
PolyU
Vobs
-aRNA
+aRNA
H477A
R1048A
N1049A
H1053A
N400A
N400G
N1055A
N1055G
N1055D
F995W
N479Q
N479A
N414A
A1052L
Wild-type
H477N, H1053N
B
PolyA -aRNA
RFU
Time (min)
H477A
R1048A
N1049A
H1053A
N400A
N400G
N1055A
N1055G
PolyA +aRNA
RFU
Time (min)
H477A
R1048A
N1049A
H1053A
N400A
N400G
N1055A
N1055G
PolyA -aRNA
RFU
Time (min)
N1055D
F995W
N479Q
N479A
N414A
A1052L
Wild-type
dLbuCas13a
PolyA +aRNA
RFU
Time (min)
N1055D
F995W
N479Q
N479A
N414A
A1052L
Wild-type
dLbuCas13a
PolyA
Vobs
-aRNA
+aRNA
H477A
R1048A
N1049A
H1053A
N400A
N400G
N1055A
N1055G
N1055D
F995W
N479Q
N479A
N414A
A1052L
Wild-type
H477N, H1053N

C

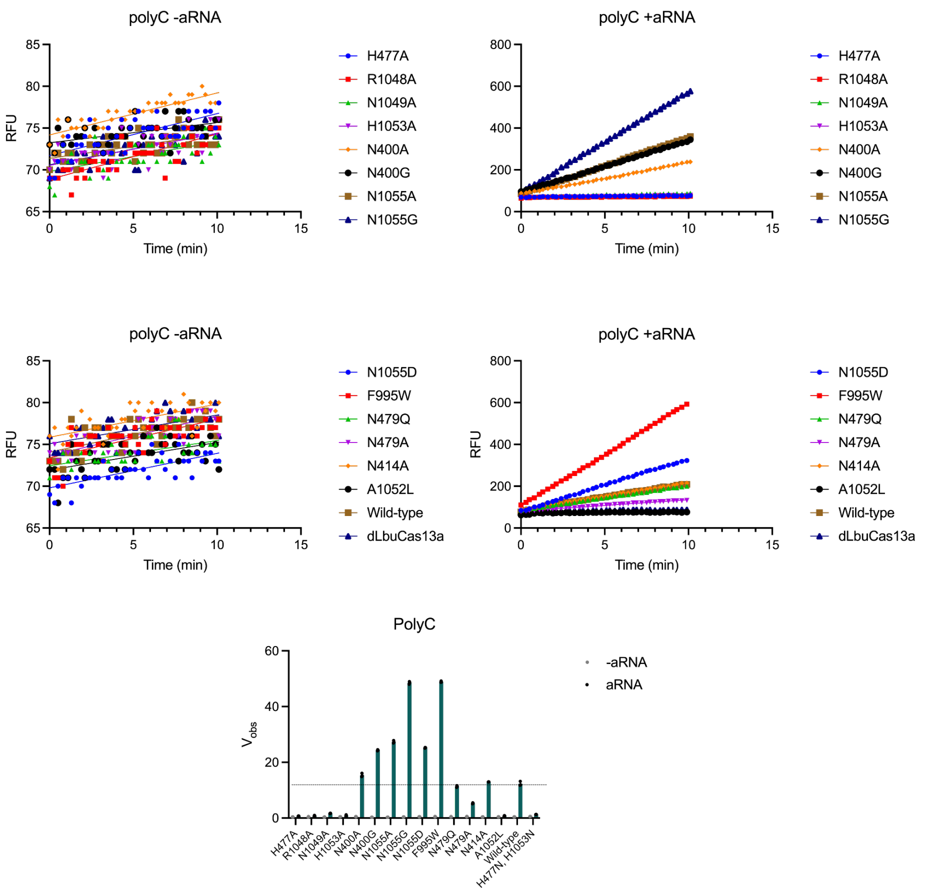
polyC -aRNA
polyC +aRNA
RFU
Time (min)
H477A
R1048A
N1049A
H1053A
N400A
N400G
N1055A
N1055G
N1055D
F995W
N479Q
N479A
N414A
A1052L
Wild-type
dLbuCas13a
PolyC
$V_{obs}$
-aRNA
aRNA
H477N, H1053N

D

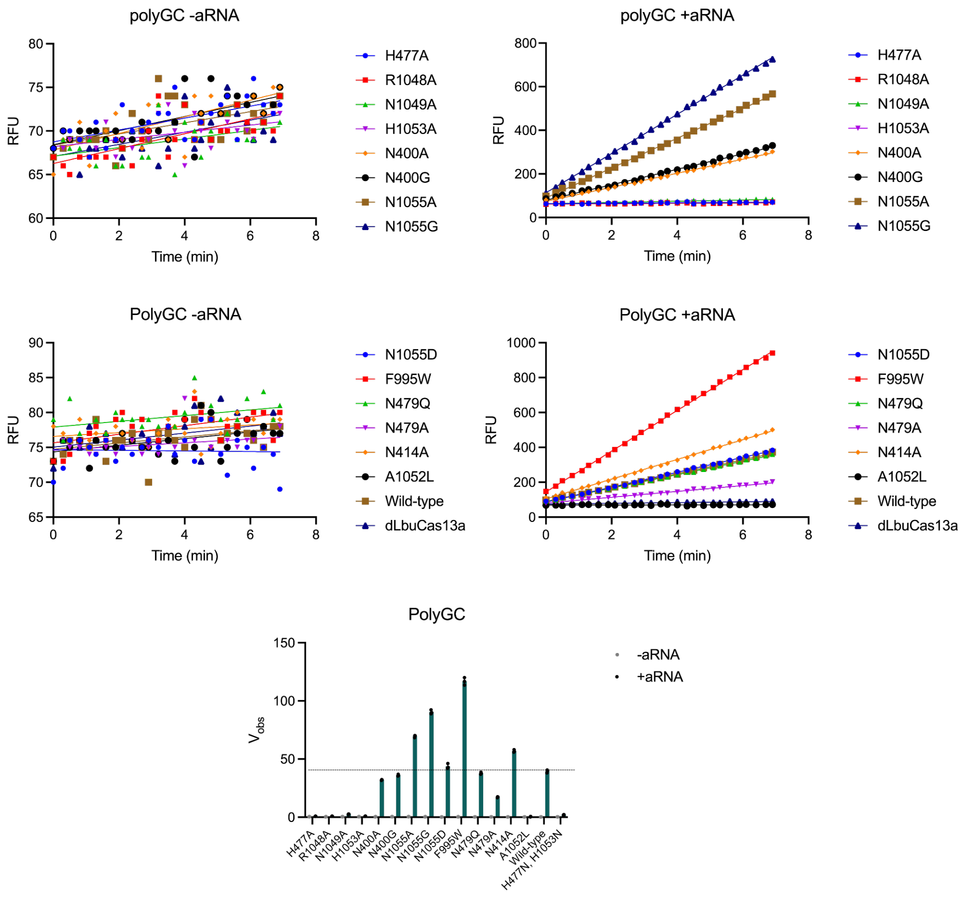
polyGC -aRNA
polyGC +aRNA
PolyGC -aRNA
PolyGC +aRNA
RFU
Time (min)
H477A
R1048A
N1049A
H1053A
N400A
N400G
N1055A
N1055G
N1055D
F995W
N479Q
N479A
N414A
A1052L
Wild-type
dLbuCas13a
PolyGC
$V_{obs}$
-aRNA
+aRNA
H477N, H1053N

**Figure S37. Raw data for activity testing of LbuCas13a specificity and HEPN mutants.** Summary of the data is presented in Figure 3, relating to mutants from **Fig. S36**. Single point mutants are as indicated, while dLbuCas13a denotes H477N, H1053N. Substrates used for the assays are (**A**) $U^5$ (rGJK123), (**B**) $A^5$ (rBKH017), (**C**) $C^5$ (rBKH005), and (**D**) CGCGC (rGJK400). Each sample has n=1 shown as an example, but n=3 was conducted for the assays with and without activator RNA (+aRNA and -aRNA, respectively). For each substrate, -aRNA is shown on the top left, +aRNA top right and bottom shows observed velocity for each of the mutants during the linear phase of the reaction. The -aRNA data is shown as purple bars with individual points as grey dots, and +aRNA data are shown as green bars with individual points in black. For all, n=3 technical replicates with the mean shown and error bars indicating SD of the mean.

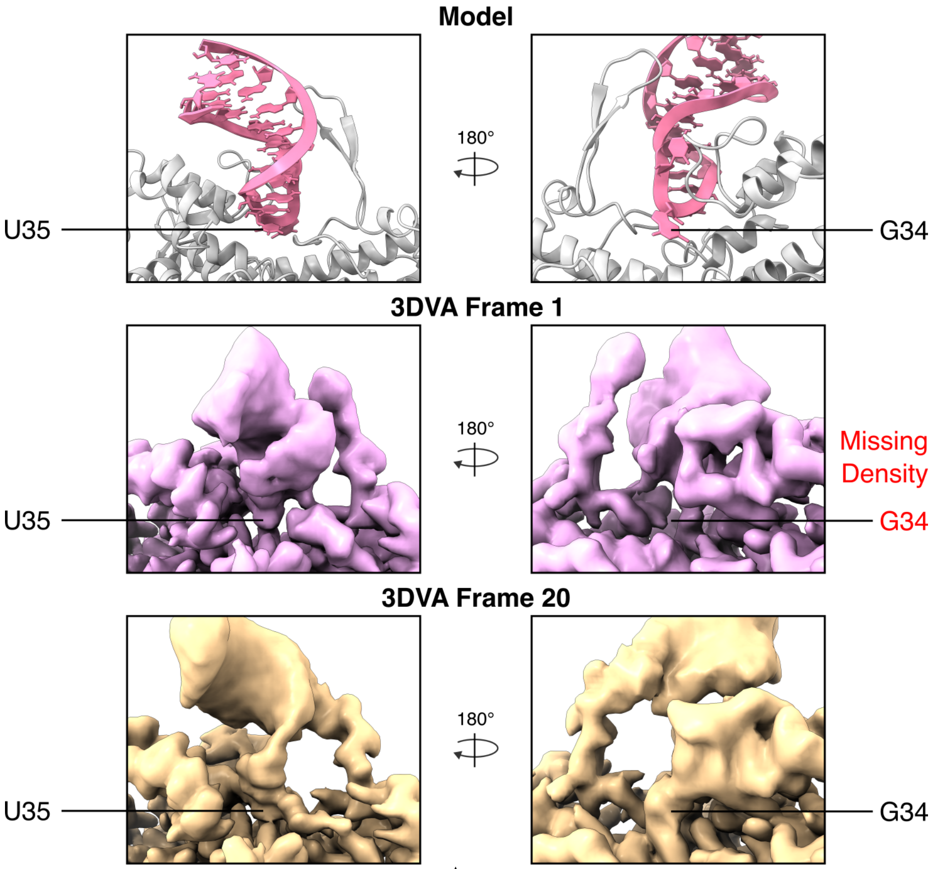


**Figure S38. Continuous ensemble of tRNA$^{Asp}$ substrate binding states showcased by 3D Variability Analysis.** Final particle stack used to produce Map E (296076 particles - FigS25) (**Table S1**) analysed with focussed 3D variability analysis (target resolution 5 Å) with large mask surrounding HEPN active site and bound substrate (See **Movie S1**). (Top) Model showing recognised U35 and flipped G34. Between frames 1 (middle) and 20 (bottom) of the 3D variability analysis, density for U35 is minimally changed but G34 is highly variable.

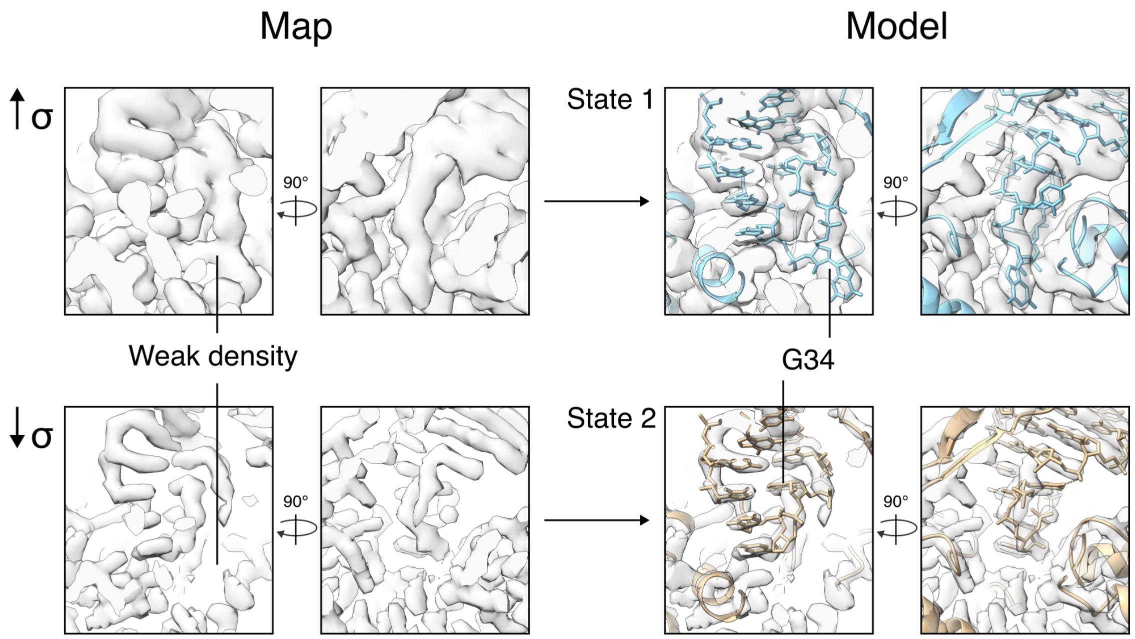


**Figure S39. Different orientation of G34 in states of tRNA$^{Asp}$ bound to LbuCas13a.** EMReady post-processed map shows flipped-G34 density in State 1 is weak, reflecting disorder exhibited in **Fig. S38**. State 2 fits G34 with the strongest density away.

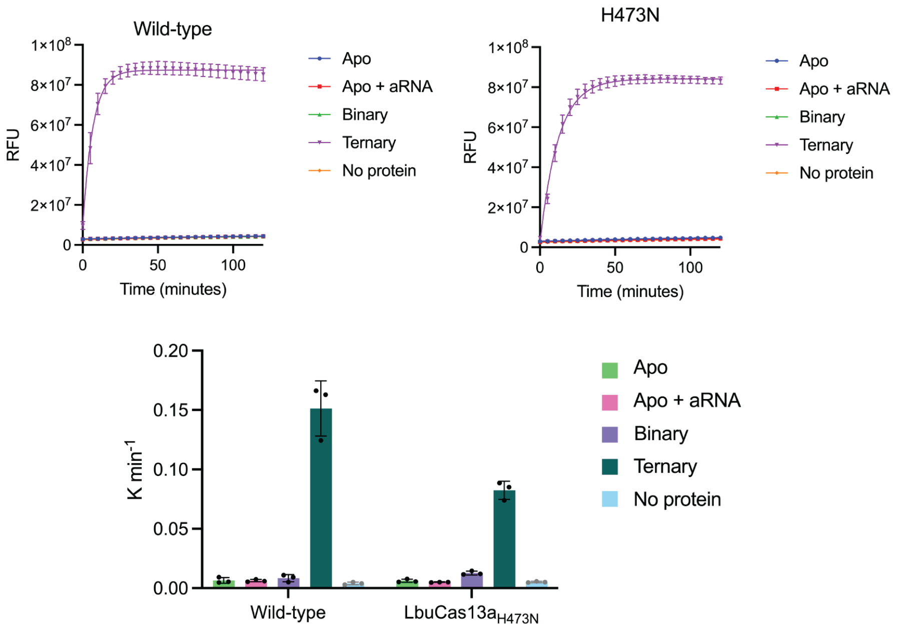


**Figure S40. Activity data for LbuCas13a$_{H473N}$ vs wildtype.** The activity of LbuCas13a$_{H473N}$ in different states (apo, apo + aRNA, binary (+crRNA) and ternary (+crRNA, +aRNA)) was compared to wild-type LbuCas13a in the equivalent states. Assays were conducted with $U^5$ (rGJK123). Top panels show the raw data with n=3 technical replicates for all states, the mean is shown as a line and error bars indicate SD. Bottom shows the rate of the reactions in the top panels which each individual rate shown, the mean rate as a bar and error bars are standard deviation of the mean.

A

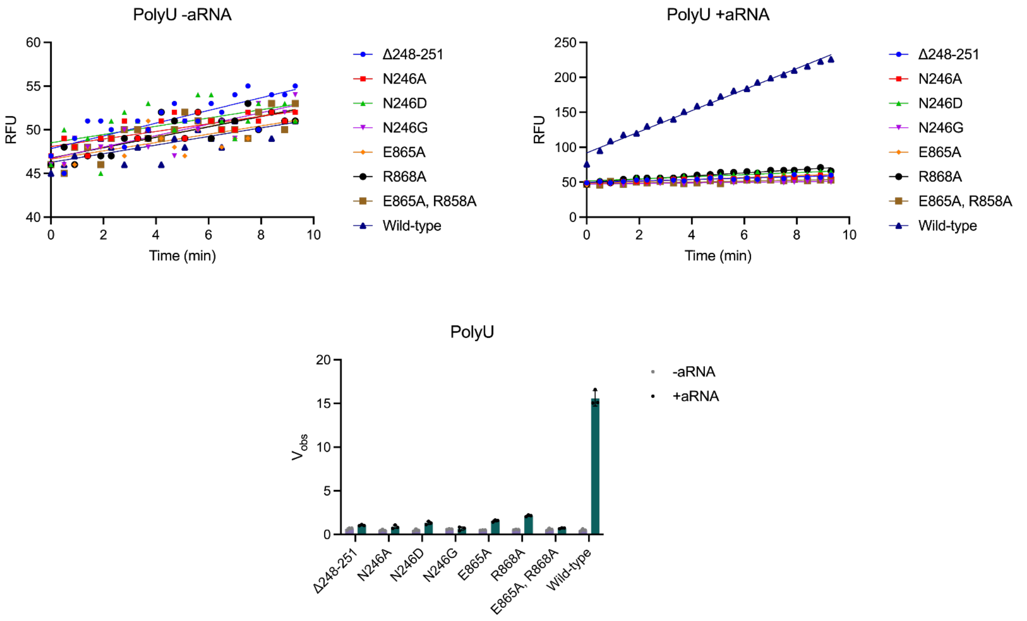
PolyU -aRNA
PolyU +aRNA
RFU
Time (min)
Δ248-251
N246A
N246D
N246G
E865A
R868A
E865A, R858A
Wild-type
PolyU
-aRNA
+aRNA

B

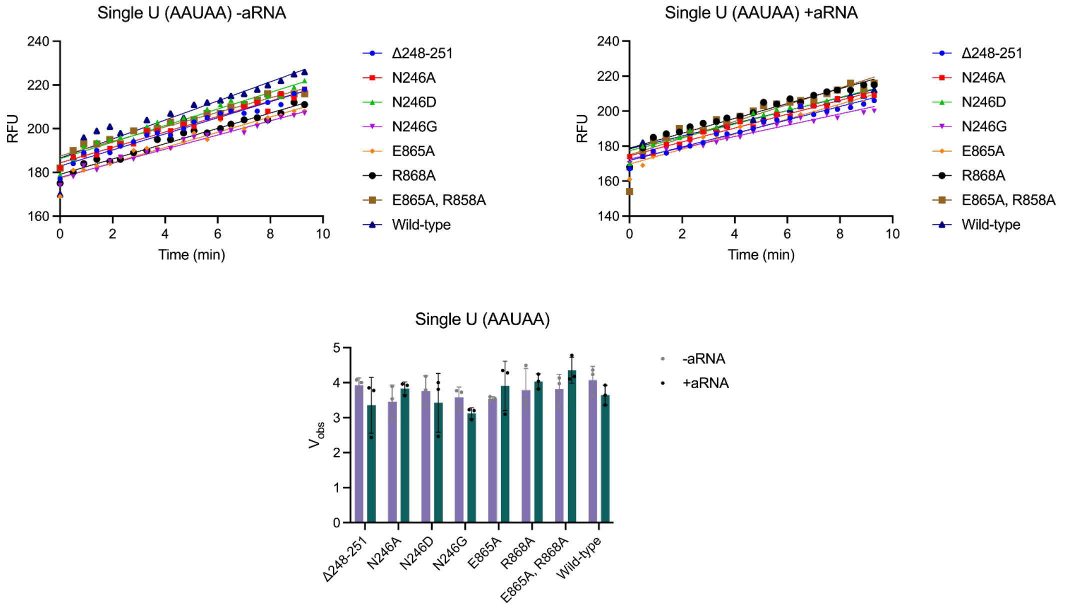
Single U (AAUAA) -aRNA
Single U (AAUAA) +aRNA
RFU
Time (min)
Δ248-251
N246A
N246D
N246G
E865A
R868A
E865A, R858A
Wild-type
Single U (AAUAA)
-aRNA
+aRNA

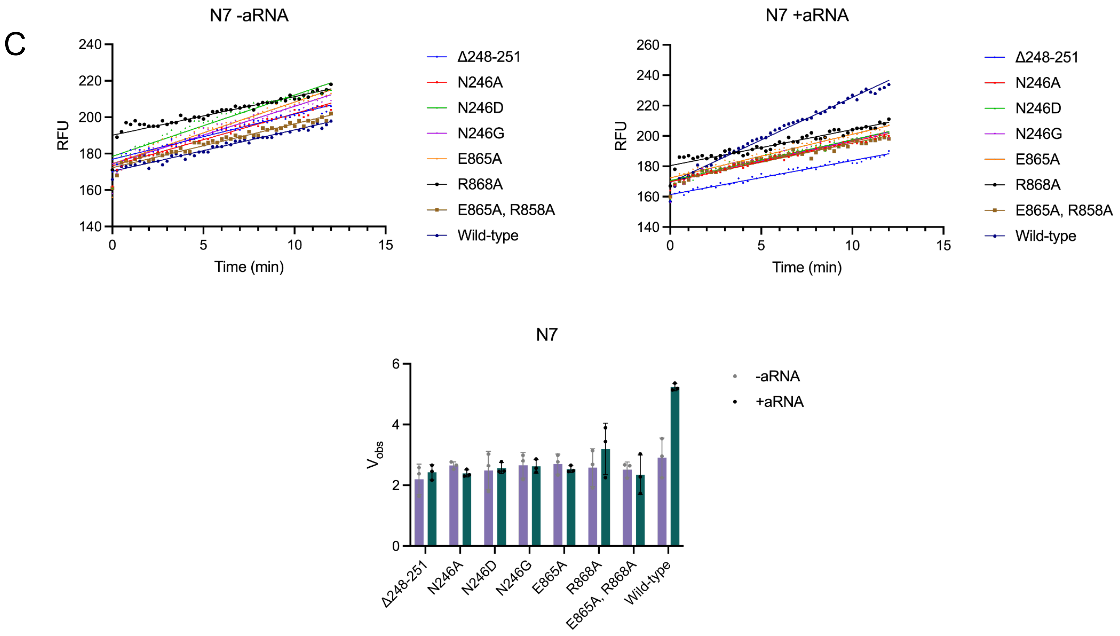


**Figure S41. Raw data for activity testing of RfxCas13d specificity mutants.** Summary of the data is presented in Figure 3, relating to mutants from **Figure S30**. Mutants are as indicated. Substrates used for the assays are (**A**) $U^5$ (rGJK123), (**B**) AAUAA (rGJK098) and (**C**) $N^7$ that includes every combination of nucleotides possible (rLVA001). Each sample has n=1 shown as an example, but n=3 was conducted for the assays with and without activator RNA (+aRNA and -aRNA, respectively). For each substrate, -aRNA is shown on the top left, +aRNA top right and bottom shows observed velocity for each of the mutants during the linear phase of the reaction. The -aRNA data is shown as purple bars with individual points as grey dots, and +aRNA data are shown as green bars with individual points in black. For all, n=3 technical replicates, with the mean shown and error bars indicating SD of the mean.

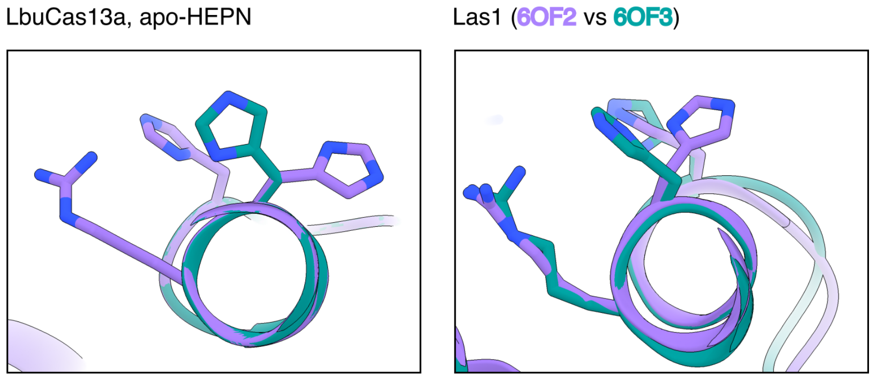


**Figure S42. Dual-occupancy of H473 resembles Las1 conformational switching.** (Left) Display of apo-HEPN LbuCas13a ternary (protein:crRNA:aRNA) H473 states facing towards (green) and away (purple) the catalytic core. (Right) H142 in Las1 (equivalent to H473 in LbuCas13a) faces towards (green, 6OF2 (*3*)) or away (purple, 6OF3 (*3*)) from the catalytic core.

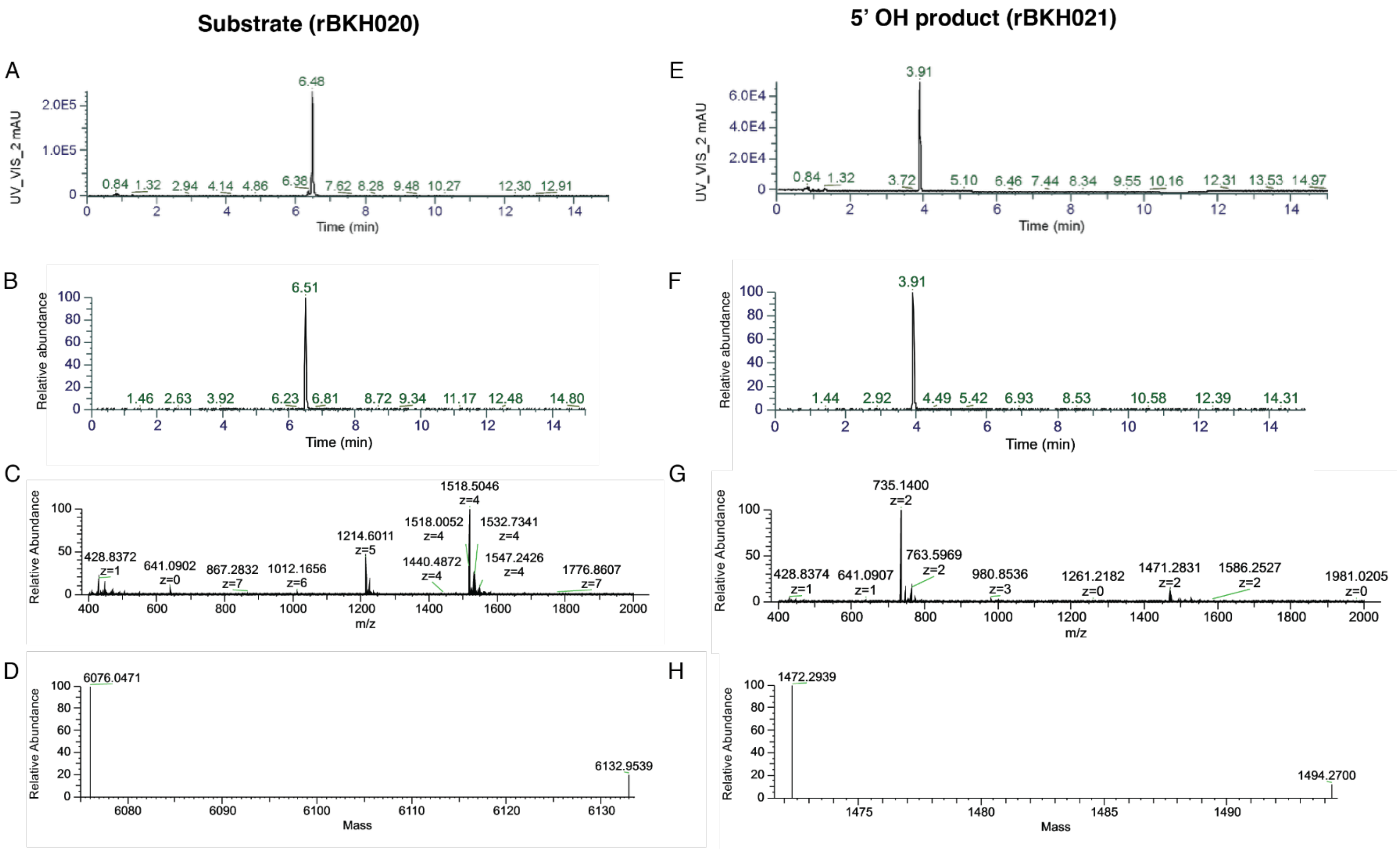


**Figure S43. LC-UV-MS data for substrate and 5′-OH product controls.** Panels A, B, C and D show the data for the substrate (rBKH020) and panels E, F, G and H show the data for the 5′-OH product (rBKH021). (**A**) and (**E**) show the LC-UV chromatogram, (**B**) and (**F**) Extracted Ion Chromatogram, (**C**) and (**G**) Full MS spectra and (**D**) and (**H**) show the deconvoluted intact masses from (C) and G).

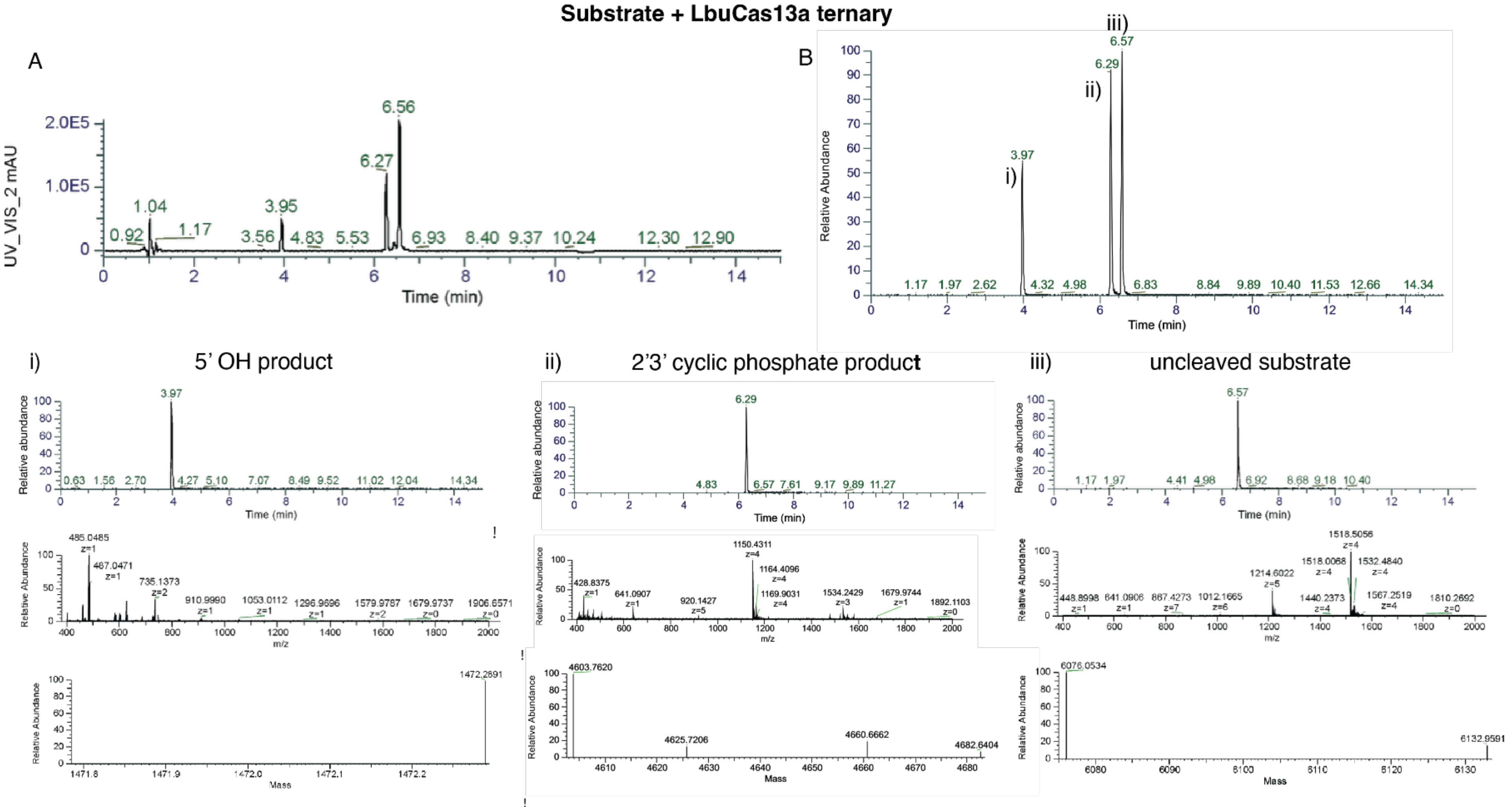


**Figure S44. LC-UV-MS data of substrate cleavage by LbuCas13a.** Substrate refers to rBKH020. (**A**) LC-UV chromatogram and (**B**) Extracted Ion Chromatogram with the target massesFurther analyses are then shown for each of the Extracted Ion Chromatogram: 5′-OH (i), 2′3′ cyclic phosphate (ii) and uncleaved substrate (iii). For panels i), ii) and iii) the top shows Extracted Ion Chromatogram , middle Full MS spectra from the peak in the top panel, and bottom is the deconvoluted intact masses.

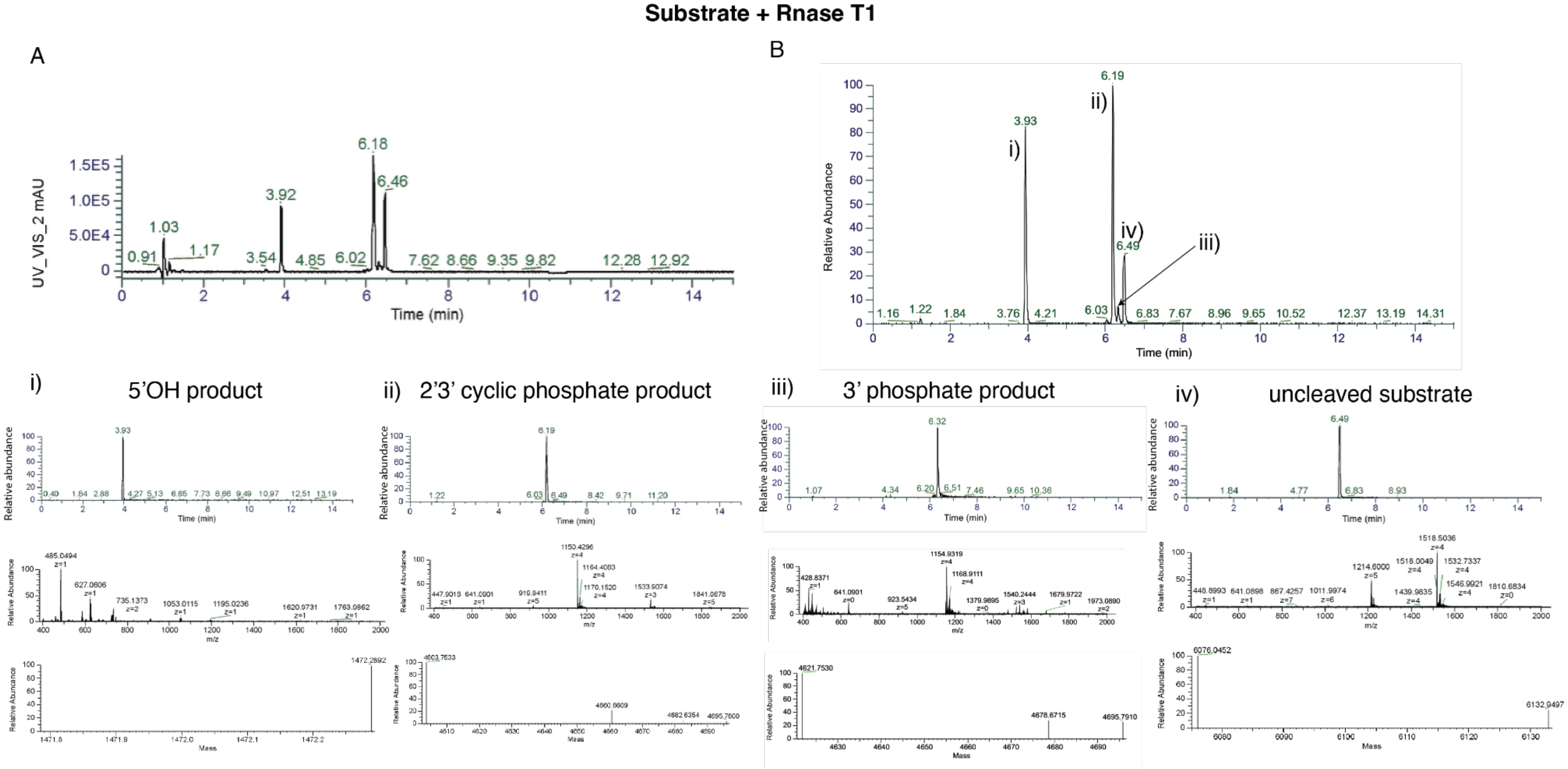


**Figure S45. LC-UV-MS data of substrate cleavage by RNase T1.** Substrate refers to rBKH020. (**A**) LC-UV chromatogram and (**B**) Extracted Ion Chromatogram with the target masses . Further analyses are then shown for each of the 3D peaks: 5′-OH (i), 2′3′ cyclic phosphate (ii), 3′ phosphate (iii) and uncleaved substrate (iv). For panels i), ii), iii) and iv) the top shows Extracted Ion Chromatogram , middle Full MS spectra from the peak in the top panel, and bottom is the deconvoluted intact masses.

### Supplementary Tables

**Table S1. Cryo-EM data collection, refinement and validation statistics.**

| | #1 Map A<br>Fig. S21<br><br>LbuCas13a<br>+crRNA<br><br><br>(EMDB-70952)<br>(PDB 9OX2) | #2 Map B<br>Fig. S21<br><br>LbuCas13a<br>+crRNA<br>+aRNA<br><br>(EMDB-70950)<br>(PDB 9OX0) | #3 Map C<br>Fig. S22<br><br>LbuCas13a<br>+crRNA<br>+$tRNA^{Lys}$<br><br>(EMDB-70953)<br>(PDB 9OX3) | #4 Map D<br>Fig. S22-23<br><br>LbuCas13a<br>+crRNA<br>+aRNA<br>+$tRNA^{Lys}$<br><br>(EMDB-70954)<br>(PDB 9OX4) | #5 Map E<br>Fig. S24<br><br>LbuCas13a<br>+crRNA<br>+aRNA<br>+$tRNA^{Asp}$<br>State 1<br><br>(EMDB-70951<br>(PDB 9OX1) | #6 Map E<br>Fig. S24<br><br>LbuCas13a<br>+crRNA<br>+aRNA<br>+$tRNA^{Asp}$<br>State 2<br><br>(EMDB-70951<br>(PDB 9OX5) |
|---|---|---|---|---|---|---|
| **Data collection and processing** | | | | | | |
| Magnification | 105 | 105 | 105 | 105 | 105 | 105 |
| Voltage (kV) | 300 | 300 | 300 | 300 | 300 | 300 |
| Electron exposure (e–/Å$^2$) | 60 | 60 | 60 | 60 | 60 | 60 |
| Defocus range (μm) | 0.5 – 1.6 | 0.5 – 1.6 | 0.5 – 1.6 | 0.5 – 1.6 | 0.5 – 1.6 | 0.5 – 1.6 |
| Pixel size (Å) | 0.82 | 0.82 | 0.82 | 0.82 | 0.82 | 0.82 |
| Symmetry imposed | C1 | C1 | C1 | C1 | C1 | C1 |
| Initial particle images (no.) | 892998 | 892998 | 613463 | 192576 | 1444332 | 1444332 |
| Final particle images (no.) | 796689 | 94831 | 123767 | 60141 | 296076 | 296076 |
| Map resolution (Å) | 2.32 | 2.63 | 2.53 | 3.10 | 2.62 | 2.62 |
| FSC threshold | 0.143 | 0.143 | 0.143 | 0.143 | 0.143 | 0.143 |
| Map resolution range (Å) | 1.789 - 37.169 | 1.773 - 42.999 | 2.183 - 39.012 | 1.749 - 51.396 | 2.225 - 35.457 | 2.225 - 35.457 |
| **Refinement** | | | | | | |
| Initial model used (PDB code) | 5XWY | 5WXP | 5XWY & 6UGG | 9OX1 | 9OX0 | 9OX0 |
| Model resolution (Å) | 3.20 | 3.09 | 3.20 & 1.95 | 2.62 | 2.63 | 2.63 |
| FSC threshold | 0.143 | 0.143 | 0.143 | 0.143 | 0.143 | 0.143 |
| Model resolution range (Å) | N/A | N/A | N/A | N/A | N/A | N/A |
| Map sharpening *B* factor (Å$^2$) | 85.0 | 72.7 | 77.2 | 90.8 | 93.1 | 93.1 |
| Model composition | 10599 | 11350 | 10940 | 11663 | 11667 | 11667 |
| Non-hydrogen atoms | 1149 | 1147 | 1146 | 1135 | 1135 | 1135 |
| Protein residues | 42 | 79 | 60 | 99 | 99 | 97 |
| RNA nucleotides | | | | | | |
| *B* factors (Å$^2$) | | | | | | |

| | | | | | | |
|---|---|---|---|---|---|---|
| Protein | 40.68 | 48.56 | 43.77 | 45.53 | 45.53 | 45.53 |
| RNA | 38.65 | 36.95 | 64.37 | 38.96 | 39.59 | 39.10 |
| R.m.s. deviations | | | | | | |
| Bond lengths (Å) | 0.002 | 0.002 | 0.003 | 0.002 | 0.002 | 0.002 |
| Bond angles (°) | 0.402 | 0.406 | 0.460 | 0.388 | 0.348 | 0.346 |
| Validation | | | | | | |
| MolProbity score | 1.50 | 1.51 | 1.46 | 1.44 | 1.25 | 1.23 |
| Clashscore | 4.45 | 4.98 | 6.12 | 5.79 | 4.76 | 4.60 |
| Poor rotamers (%) | 0 | 0.09 | 0 | 0.85 | 0.28 | 0.28 |
| Ramachandran plot | | | | | | |
| Favored (%) | 95.98 | 96.33 | 97.37 | 97.34 | 98.05 | 98.23 |
| Allowed (%) | 4.02 | 3.67 | 2.63 | 2.66 | 1.95 | 1.77 |
| Disallowed (%) | 0 | 0 | 0 | 0 | 0 | 0 |

**Table S2. Bacterial strains and plasmids used in this study.**

| Bacterial strain | Description | Source |
| --- | --- | --- |
| *Escherichia coli* | | |
| BL21(DE3) | For recombinant protein expression and generating total E. coli RNA.<br>fhuA2 [lon] ompT gal (λ DE3) [dcm] ΔhsdS<br>λ DE3 = λ sBamHIo ΔEcoRI-B int::(lacI::PlacUV5::T7 gene1) i21 Δnin5 | Merck |
| DH5α | For general cloning.<br>F– φ80lacZΔ M15 Δ (lacZYA-argF) U169 recA1 endA1 hsdR17 (rK– mK+) phoA supE44 λ- thi–1 gyrA96 relA1 | Merck |
| NEB Turbo | For cloning LbuCas13a mutants.<br>F' proA+B+ lacIq ΔlacZM15 / fhuA2 Δ(lac-proAB) glnV galK16 galE15 R(zgb-210::Tn10)TetS endA1 thi-1 Δ(hsdS-mcrB)5 | New England Biolabs |
| NEB 10-beta | For cloning RfxCas13d mutants.<br>Δ(ara-leu) 7697 araD139 fhuA ΔlacX74 galK16 galE15 e14- ϕ80dlacZΔM15 recA1 relA1 endA1 nupG rpsL (StrR) rph spoT1 Δ(mrr-hsdRMS-mcrBC) | New England Biolabs |
| Plasmids | | |
| Name | Description | Source |
| pGJK191 | LbuCas13a expression plasmid. N-terminal hexahistidine tag, MBP and Tev site. T7 promoter, Amp resistance. | Addgene #83482 |
| pGJK189 | LbaCas13a expression plasmid. N-terminal hexahistidine tag, MBP and Tev site. T7 promoter, Amp resistance. | Addgene #91867 |
| pRWC021 | LbuCas13a finger deletion mutant. Cloned using pGJK191 backbone to remove residues 409 - 421. | This study |
| pRWC020 | LbuCas13a catalytically dead mutant. Cloned using pGJK191 backbone to introduce H477N + H1053N. | This study |
| pBKH002 - 005 | LbuCas13a substrate binding pocket mutants. Cloned using KLD with pGJK191 backbone to insert mutations. Mutations are as follows (in order of numbering): R405A, F995A, Q1007A and E1061A | This study |
| pBKH006 | As rBKH002–005 but multiple mutations introduced: K1036A, K1039A, Q1040A and K1042A | This study |
| pBKH007 | As rBKH002–005 but multiple mutations introduced: K409A, | This study |

| | | |
|---|---|---|
| | N414A, K415A and K419A | |
| pBKH008 | As rBKH002–005 but multiple mutations introduced: K1004A, K1011A, K1014A and K1017A | This study |
| pBKH015 - 028 | LbuCas13a catalysis and substrate specificity mutants. Cloned using KLD with pGJK191 backbone to insert mutations. Mutations are as follows (in order of numbering): H477A, R1048A, N1049A, H1053A, N400A, N400G, N1055A, N1055G, N1055D, F995W, N479Q, N479A, N414A and A1052L. | This study |
| pBKH011 | RfxCas13d expression plasmid. C-terminal hexahistidine tag. pET28a+ backbone. | This study |
| pBKH009 | PspCas13b expression plasmid. C-terminal hexahistidine tag. pET29b+ backbone. | This study |
| pNC021 - 023 | RfxCas13d mutants to increase the positive charge of the substrate binding pocket. Cloned using KLD with pBKH011 backbone to insert mutations. Mutations are as follows (in order of numbering): A188K + A191K, E753K + E754K and N897K + E898K + E901K | This study |
| pNC024 - 027 | RfxCas13d mutants that contain residues 408 to 423 of LbuCas13a. Cloned using KLD with pBKH011 backbone to insert mutations. Insertions sites are as follows (in order of numbering): D183 and K185, N188 and A189, A189 and I190, A192 and Q193. | This study |
| pNC028 - 034 | RfxCas13d substrate specificity mutants. Cloned using KLD with pBKH011 backbone to insert mutations. Mutations are as follows (in order of numbering): E248G + Δ249-251, N247A, N247D, N247G, E865A, R869A and E865A + R869A | This study |

**Table S3. RNA used in this study.**

| Name | Description | Sequence (5′ to 3′) | Source |
|---|---|---|---|
| CRISPR RNA | | | |
| rGJK072 | LbuCas13a CRISPR RNA | UAGACCAGCCCAAAAAUGAAGGGCACUAAAACGCAGCGCCUCUUGCAACGAUUAAA | This study |
| rGJK074 | LbuCas13a and RfxCas13d activator RNA | UUUAAUCGUUGCAAGAGGCGCUGCUC | This study |
| rGJK119 | LbuCas13a CRISPR RNA | UAGACCACCCCAAAAAUGAAGGGGACUAAAACUUUCUUUCUUUCCUUUUUCUGCCG | This study |
| rGJK120 | LbuCas13a activator RNA | CGGCAGAAAAAGGAAAGAAAGAAACC | This study |
| rBKH022 | RfxCas13d CRISPR RNA | AACCCCUACCAACUGGUCGGGGUUUGAAACGCAGCGCCUCUUGCAACGAUUAAA | This study |
| Other RNA | | | |
| | | *Activity Assays* | |
| rGJK123 | polyU fluorescent substrate ($U^5$) | **6FAM**/UUUUU/**IowaBlackQ** | East-Seletsky, O'Connell, Burstein, Knott and Doudna (*33*) |
| rBKH005 | polyC fluorescent substrate ($C^5$) | **6FAM**/CCCCC/**IowaBlackQ** | This study |
| rGJK400 | polyGC fluorescent substrate | **6FAM**/CGCGC/**IowaBlackQ** | This study |
| rBKH017 | polyA fluorescent substrate ($A^5$) | **6FAM**/AAAAA/**IowaBlackQ** | This study |
| rRWC041 | $ASL^{Lys(UUU)}$ | **6FAM**/dCGCUUGACUUUUAAUCAAGCdC/**IowaBlackQ** | This study |
| rBKH002 | $tRNA^{Lys(UUU)}$ ACN loop | **6FAM**/CUUUUAA/**IowaBlackQ** | This study |
| rLVA001 | $N^7$ fluorescent substrate | **6FAM**/NNNNNNN/**IowaBlackQ** | This study |

| | | | |
|---|---|---|---|
| rGJK098 | Fluorescent substrate | **6FAM**/AAUAA/**IowaBlackQ** | This study |
| rGJK099 | Fluorescent substrate with pseudouridine | **6FAM**/AApUAA/**IowaBlackQ** | This study |
| | | *HPLC-MS* | |
| rBKH020 | DNA polymer with single RNase cut site | dCdAdTdTdCdGdCdAdTdTdCdCdAdG GdUdCdAdAdG | This study |
| rBKH021 | Control 3′ product for rBKH020 | dUdCdAdAdG | This study |
| | | *RNA Gels* | |
| rRWC009 | Lbu tRNA$^{Lys(UUU)}$ | **6FAM**/dCGAGCCAUUAGCUCAGUC GGUAGAGCACUUGACUUUUAAUC AAGGUGUCACUGGUUCGAUCCCA GUAUGGCUCACCA | This study |
| rRWC043 | Scrambled Lbu tRNA$^{Lys(UUU)}$ | **6FAM**/dCAUCGUUAGACCGUAUCU UCGCCCACAUUGACACUAUGCGA GUGGUUCGAUUCAGUCGUGAUGA GAGACGAUUCACC | This study |
| rRWC023 | ASL$^{Lys(UUU)}$ | **6FAM**/dCGCUUGACUUUUAAUCAA GC | This study |
| rRWC024 | Scrambled ASL$^{Lys(UUU)}$ | **6FAM**/dCGGUCUGUAAACUUAUAU CC | This study |
| rRWC025 | ASL$^{Lys(UUU)}$ with swapped stems | **6FAM**/dCGGAACUCUUUUAAAGUU CC | This study |
| rRWC026 | ASL$^{Phe(GAA)}$ | **6FAM**/dCGAGGGACUGAAAAUCCC UC | This study |
| rRWC027 | Scrambled ASL$^{Phe(GAA)}$ | **6FAM**/dCGCACGUCCAGUAGAUAG AC | This study |
| rRWC028 | ASL$^{Phe(GAA)}$ with swapped stems | **6FAM**/dCGUCCCUCUGAAAAAGGG AC | This study |
| rRWC029 | ASL$^{Lys(UUU)}$ with single deoxy-blocked cut site | **6FAM**/dCGCUUGA**dC**UUUUAAUCA AGC | This study |
| rRWC030 | ASL$^{Lys(UUU)}$ with single deoxy-blocked cut site | **6FAM**/dCGCUUGAC**dU**UUUAAUCA AGC | This study |
| rRWC031 | ASL$^{Lys(UUU)}$ with single deoxy-blocked cut site | **6FAM**/dCGCUUGACU**dU**UUAAUCA AGC | This study |

| | | | |
|---|---|---|---|
| rRWC032 | ASL$^{Lys(UUU)}$ with single deoxy-blocked cut site | **6FAM**/dCGCUUGACUU**d**UUAAUCAAGC | This study |
| rRWC033 | ASL$^{Lys(UUU)}$ with 4 deoxy-blocked cut sites | **6FAM**/dCGCUUGA**dCdUdUdU**UAAUCAAGC | This study |
| rRWC034 | ASL$^{Ph(GAA)}$ with single deoxy-blocked cut site | **6FAM**/dCGAGGGA**dC**UGAAAAUCCCUC | This study |
| rRWC035 | ASL$^{Lys(UUU)}$ with 2 deoxy-blocked cut sites | **6FAM**/dCGCUUGACU**dUdU**UAAUCAAGC | This study |
| rRWC036 | ASL$^{Lys(UUU)}$ with reversed loop | **6FAM**/dCGCUUGAAAUUUUCUCAAGC | This study |
| rRWC037 | ASL$^{Lys(UUU)}$ swapped stems and reversed loop | **6FAM**/dCGGAACUAAUUUUCAGUUCC | This study |
| rRWC038 | ASL$^{Lys(UUU)}$ loop & ASL$^{Phe(GAA)}$ stems | **6FAM**/dCGAGGGACUUUUAAUCCCUC | This study |
| rRWC063 | ASL$^{Asp(GUC)}$ | **6FAM**/dCGCUUGCCUGUCACGCAAGC | |
| *Competition Assay* | | | |
| rRWC039 **CryoEM Substrate** | ASL$^{Lys(UUU)}$ with 2 deoxy-blocked cut sites | GCUUGACU**dUdU**UAAUCAAGC | This study |
| rRWC060 **CryoEM Substrate** | ASL$^{Asp(GUC)}$ with single 2′ F-blocked cut site | GCUUGCCU**fG**UCACGCAAGC | This study |
| rRWC013 | U$^5$ | UUUUU | This study |
| rRWC050 | ASL$^{Lys(UUU)}$ | GCUUGACUUUUAAUCAAGC | This study |
| rRWC051 | ASL$^{Phe(GAA)}$ | GAGGGACUGAAAAUCCCUC | This study |
| rRWC059 | ASL$^{Asp(GUC)}$ | GCUUGCCUGUCACGCAAGC | This study |

*All RNA were purchased from IDT unless stated otherwise

**Table S4. Concentration of aRNA for mutant activity assays.**

| LbuCas13a mutant assays | | |
|---|---|---|
| Substrate used | aRNA concentration | Notes |
| PolyU (rGJK123) | 40 pM | |
| PolyA (rBKH017) | 15 nM | |
| PolyGC (rGJK400) | 100 nM | |
| PolyC (rBKH005) | 100 or 200 nM | For Fig 3. and Fig 2. mutants respectively |
| $ASL^{Lys(UUU)}$ (rRWC041) | 80 pM | |
| $tRNA^{Lys/UUU}$ ACN loop (rBKH002) | 20 pM | |
| PolyU (rGJK123) | 10 nM | $LbuCas13a_{H473N}$ comparison assay |
| RfxCas13d mutant assays | | |
| Substrate used | aRNA concentration | Notes |
| PolyU (rGJK123) | 500 pM | |
| Single U (rGJK098: AAUAA) | 500 pM | |
| $N^7$ (rLVA001:NNNNNNN) | 500 pM | |
